# Pancreatic progenitor epigenome maps prioritize type 2 diabetes risk genes with roles in development

**DOI:** 10.1101/2020.05.18.101071

**Authors:** Ryan J. Geusz, Allen Wang, Joshua Chiou, Joseph J. Lancman, Nichole Wetton, Samy Kefalopoulou, Jinzhao Wang, Yunjiang Qiu, Jian Yan, Anthony Aylward, Bing Ren, P Duc Si Dong, Kyle J. Gaulton, Maike Sander

## Abstract

Genetic variants associated with type 2 diabetes (T2D) risk affect gene regulation in metabolically relevant tissues, such as pancreatic islets. Here, we investigated contributions of regulatory programs active during pancreatic development to T2D risk. Generation of chromatin maps from developmental precursors throughout pancreatic differentiation of human embryonic stem cells (hESCs) identifies enrichment of T2D variants in pancreatic progenitor-specific stretch enhancers that are not active in islets. Genes associated with progenitor-specific stretch enhancers are predicted to regulate developmental processes, most notably tissue morphogenesis. Through gene editing in hESCs, we demonstrate that progenitor-specific enhancers harboring T2D-associated variants regulate cell polarity genes *LAMA1* and *CRB2*. Knockdown of *lama1* or *crb2* in zebrafish embryos causes a defect in pancreas morphogenesis and impairs islet cell development. Together, our findings reveal that a subset of T2D risk variants specifically affects pancreatic developmental programs, suggesting that dysregulation of developmental processes can predispose to T2D.

## INTRODUCTION

Type 2 diabetes (T2D) is a multifactorial metabolic disorder characterized by insulin insensitivity and insufficient insulin secretion by pancreatic beta cells (Halban et al., 2014). Genetic association studies have identified hundreds of loci influencing risk of T2D (Mahajan et al., 2018). However, disease-relevant target genes of T2D risk variants, the mechanisms by which these genes cause disease, and the tissues in which the genes mediate their effects remain poorly understood.

The majority of T2D risk variants map to non-coding sequence, suggesting that genetic risk of T2D is largely mediated through variants affecting transcriptional regulatory activity. Intersection of T2D risk variants with epigenomic data has uncovered enrichment of T2D risk variants in regulatory sites active in specific cell types, predominantly in pancreatic beta cells, including risk variants that affect regulatory activity directly (Chiou et al., 2019; Fuchsberger et al., 2016; Gaulton et al., 2015; Gaulton et al., 2010; Greenwald et al., 2019; Mahajan et al., 2018; Parker et al., 2013; Pasquali et al., 2014; Thurner et al., 2018; Varshney et al., 2017). T2D risk-associated variants are further enriched within large, contiguous regions of islet active chromatin, referred to as stretch or super-enhancers (Parker et al., 2013). These regions of active chromatin preferentially bind islet cell-restricted transcription factors and drive islet-specific gene expression (Parker et al., 2013; Pasquali et al., 2014).

Many genes associated with T2D risk in islets are not uniquely expressed in differentiated islet endocrine cells, but also in pancreatic progenitor cells during embryonic development. For example, T2D risk variants map to *HNF1A, HNF1B, HNF4A, MNX1, NEUROG3, PAX4,* and *PDX1* (Flannick et al., 2019; Mahajan et al., 2018; Steinthorsdottir et al., 2014), which are all transcription factors also expressed in pancreatic –developmental precursors. Studies in model organisms and hESC-based models of pancreatic endocrine cell differentiation have shown that inactivation of these transcription factors causes defects in endocrine cell development, resulting in reduced beta cell numbers (Gaertner et al., 2019). Furthermore, heterozygous mutations for *HNF1A, HNF1B, HNF4A, PAX4,* and *PDX1* are associated with maturity onset diabetes of the young (MODY), which is an autosomal dominant form of diabetes with features similar to T2D (Urakami, 2019). Thus, there is evidence that reduced activity of developmentally expressed transcription factors can cause diabetes later in life.

The role of these transcription factors in T2D and MODY could be explained by their functions in regulating gene expression in mature islet cells. However, it is also possible that their function during endocrine cell development could predispose to diabetes instead of, or in addition to, endocrine cell gene regulation. One conceivable mechanism is that individuals with reduced activity of these transcription factors are born with either fewer beta cells or beta cells more prone to fail under conditions of increased insulin demand. Observations showing that disturbed intrauterine metabolic conditions, such as maternal malnutrition, can lead to reduced beta cell mass and T2D predisposition in the offspring (Lumey et al., 2015; Nielsen et al., 2014; Portha et al., 2011) support the concept that compromised beta cell development could predispose to T2D. However, whether there is T2D genetic risk relevant to the regulation of endocrine cell development independent of gene regulation in mature islet cells has not been explored.

In this study, we investigated the contribution of gene regulatory programs specifically active during pancreatic development to T2D risk. First, we employed a hESC-based differentiation system to generate chromatin maps of hESCs during their stepwise differentiation into pancreatic progenitor cells. We then identified T2D-associated variants localized in active enhancers in developmental precursors but not in mature islets, used genome editing in hESCs to define target genes of pancreatic progenitor-specific enhancers harboring T2D variants, and employed zebrafish genetic models to study the role of two target genes in pancreatic and endocrine cell development.

## RESULTS

### Pancreatic progenitor stretch enhancers are enriched for T2D risk variants

To determine whether there is a development-specific genetic contribution to T2D risk, we generated genome-wide chromatin maps of hESCs during their stepwise differentiation into pancreatic progenitors through four distinct developmental stages: definitive endoderm (DE), gut tube (GT), early pancreatic progenitors (PP1), and late pancreatic progenitors (PP2) (**Figure 1A**). We then used ChromHMM (Ernst & Kellis, 2012) to annotate chromatin states, such as active promoters and enhancers, at all stages of hESC differentiation as well as in primary islets (**Figure 1 – figure supplement 1A,B**).

**Figure 1:**
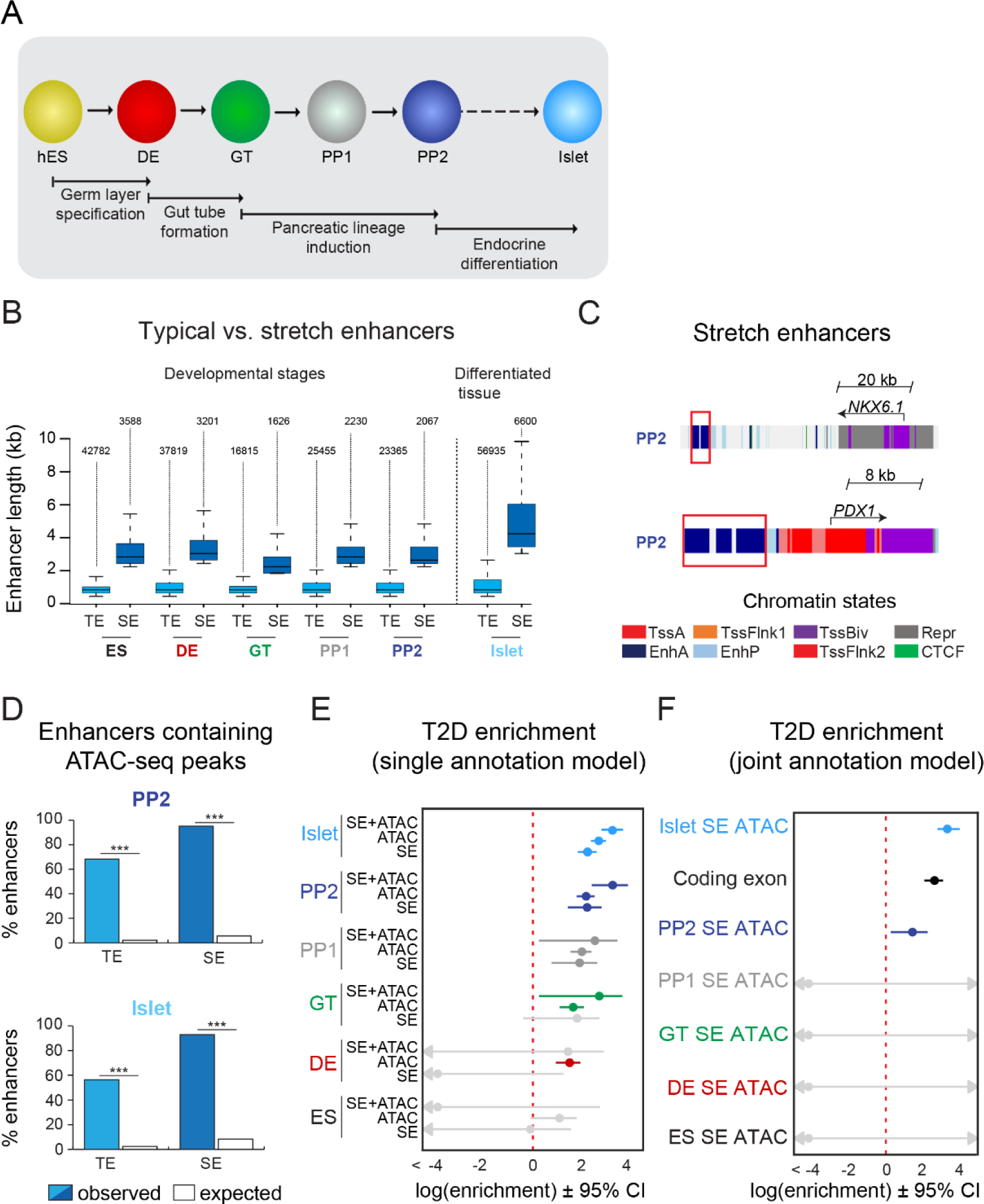
T2D-associated risk variants are enriched in stretch enhancers of pancreatic progenitors independent of islet stretch enhancers. (A) Schematic illustrating the stepwise differentiation of human embryonic stem cells (hES) into pancreatic progenitors (solid arrows) and lineage relationship to islets (dotted arrow). Developmental intermediates include definitive endoderm (DE), gut tube (GT), early pancreatic progenitor (PP1), and late pancreatic progenitor (PP2) cells. (B) Box plots depicting length of typical enhancers (TE) and stretch enhancers (SE) at each developmental stage and in primary human islets. Plots are centered on median, with box encompassing 25th-75th percentile and whiskers extending up to 1.5 interquartile range. Total numbers of enhancers are shown above each box plot. (C) Examples of stretch enhancers near the genes encoding the pancreatic lineage-determining transcription factors NKX6.1 and PDX1, respectively. Chromatin states are based on ChromHMM classifications: TssA, active promoter; TssFlnk, flanking transcription start site; TssBiv, bivalent promoter; Repr, repressed; EnhA, active enhancer; EnhP, poised enhancer. (D) Percentage of TE and SE overlapping with at least one ATAC-seq peak at PP2 or in islets. Enrichment analysis comparing observed and expected overlap based on random genomic regions of the same size and located on the same chromosome averaged over 10,000 iterations (*** p < 1 × 10^−4^; permutation test). (**E**) Genome-wide enrichment of T2D-associated variants in stretch enhancers, ATAC-seq peaks, and ATAC-seq peaks within stretch enhancers for all developmental stages when modelling each annotation separately. Points and lines represent log-scaled enrichment estimates and 95% confidence intervals from fgwas, respectively. (**F**) Genome-wide enrichment of T2D-associated variants in ATAC-seq peaks within stretch enhancers for all developmental stages and coding exons when considering all annotations in a joint model. Points and lines represent log-scaled enrichment estimates and 95% confidence intervals from fgwas, respectively. See also Figure 1 – figure supplement 1.

Large, contiguous regions of active enhancer chromatin, which have been termed stretch- or super-enhancers (Parker et al., 2013; Whyte et al., 2013), are highly enriched for T2D risk variants in islets (Parker et al., 2013; Pasquali et al., 2014). We therefore partitioned active enhancers from each hESC developmental stage and islets into stretch enhancers (SE) and traditional (non-stretch) enhancers (TE) (**Figure 1B**). Consistent with prior observations of SE features (Parker et al., 2013; Whyte et al., 2013), SE comprised a small subset of all active enhancers (7.7%, 7.8%, 8.8%, 8.1%, 8.1%, and 10.4% of active enhancers in ES, DE, GT, PP1, PP2, and islets, respectively; **Figure 1B** and **Figure 1 – figure supplement 1C**) and genes proximal to SE were more highly expressed than genes proximal to TE (p = 4.68 × 10^−7^, 4.64 × 10^−11^, 1.31 × 10^−5^, 8.85 × 10^−9^, 5.34 × 10^−6^, and < 2.2 × 10^−16^ for expression of genes near TE vs SE in ES, DE, GT, PP1, PP2, and islets, respectively; **Figure 1 – figure supplement 1D**). Genes near SE in pancreatic progenitors included transcription factors involved in the regulation of pancreatic cell identity, such as *NKX6.1* and *PDX1* (**Figure 1C**). Since disease-associated variants are preferentially enriched in narrow peaks of accessible chromatin within broader regions of active chromatin (Greenwald et al., 2019; Thurner et al., 2018; Varshney et al., 2017), we next used ATAC-seq to generate genome-wide maps of chromatin accessibility across all time points of differentiation. Nearly all identified SE contained at least one ATAC-seq peak (**Figure 1D** and **Figure 1 – figure supplement 1E,F**). At the PP2 stage, 62.3% of SE harbored one, 32.2% two or three, and 0.7% four or more ATAC-seq peaks (**Figure 1 – figure supplement 1F**). Similar percentages were observed in earlier developmental precursors and islets.

Having annotated accessible chromatin sites within SE, we next tested for enrichment of T2D-associated variants in SE active in mature islets and in pancreatic developmental stages. We observed strongest enrichment of T2D-associated variants in islet SE (log enrichment = 2.18, 95% CI = 1.80, 2.54) and late pancreatic progenitor SE (log enrichment = 2.17, 95% CI = 1.40, 2.74), which was more pronounced when only considering variants in accessible chromatin sites within these elements (islet log enrichment = 3.20, 95% CI = 2.74, 3.60; PP2 log enrichment = 3.18, 95% CI = 2.35, 3.79; **Figure 1E**). Given that a subset of pancreatic progenitor SE is also active in islets, we next determined whether pancreatic progenitor SE contribute to T2D risk independently of islet SE. Variants in accessible chromatin sites of late pancreatic progenitor SE were enriched for T2D association in a joint model including islet SE (islet log enrichment = 2.94, 95% CI = 2.47, 3.35; PP2 log enrichment = 1.27, 95% CI = 0.24, 2.00; **Figure 1F**). We also observed enrichment of variants in accessible chromatin sites of pancreatic progenitor SE after conditioning on islet SE (log enrichment = 0.60, 95% CI = -0.87, 1.48), as well as when excluding pancreatic progenitor SE active in islets (log enrichment = 1.62, 95% CI = <-20, 3.14). Examples of known T2D loci with T2D-associated variants in SE active in pancreatic progenitors but not in islets included *LAMA1* and *PROX1*. These results suggest that a subset of T2D variants may affect disease risk by altering regulatory programs specifically active in pancreatic progenitors.

### Pancreatic progenitor-specific stretch enhancers are near genes that regulate tissue morphogenesis

Having observed enrichment of T2D risk variants in pancreatic progenitor SE independent of islet SE, we next sought to further characterize the regulatory programs of SE with specific function in pancreatic progenitors. We therefore defined a set of pancreatic progenitor-specific stretch enhancers (PSSE) based on the following criteria: (i) annotation as a SE at the PP2 stage, (ii) no classification as a SE at the ES, DE, and GT stages, and (iii) no classification as a TE or SE in islets. Applying these criteria, we identified a total of 492 PSSE genome-wide (**Figure 2A** and **Figure 2 – source data 1**). As expected based on their chromatin state classification, PSSE acquired broad deposition of the active enhancer mark H3K27ac at the PP1 and PP2 stages (**Figure 2B,C**). Coincident with an increase in H3K27ac signal, chromatin accessibility at PSSE also increased (**Figure 2B**), and 93.5% of PSSE contained at least one accessible chromatin site at the PP2 stage (**Figure 2 – figure supplement 1A,B**). Further investigation of PSSE chromatin state dynamics at earlier stages of pancreatic differentiation revealed that PSSE were often poised (defined by H3K4me1 in the absence of H3K27ac) prior to activation (42%, 48%, 63%, and 17% of PSSE in ES, DE, GT, and PP1, respectively; **Figure 2C**), consistent with earlier observations that a poised enhancer state frequently precedes enhancer activation during development (Rada-Iglesias et al., 2011; Wang et al., 2015). Intriguingly, a subset of PSSE was classified as TE earlier in development (13%, 23%, 29%, and 46% of PSSE in ES, DE, GT, and PP1, respectively; **Figure 2C**), suggesting that SE emerge from smaller regions of active chromatin seeded at prior stages of development. During differentiation into mature islet cells, PSSE lost H3K27ac but largely retained H3K4me1 signal (62% of PSSE) (**Figure 2C**), persisting in a poised state in terminally differentiated islet cells.

**Figure 2:**
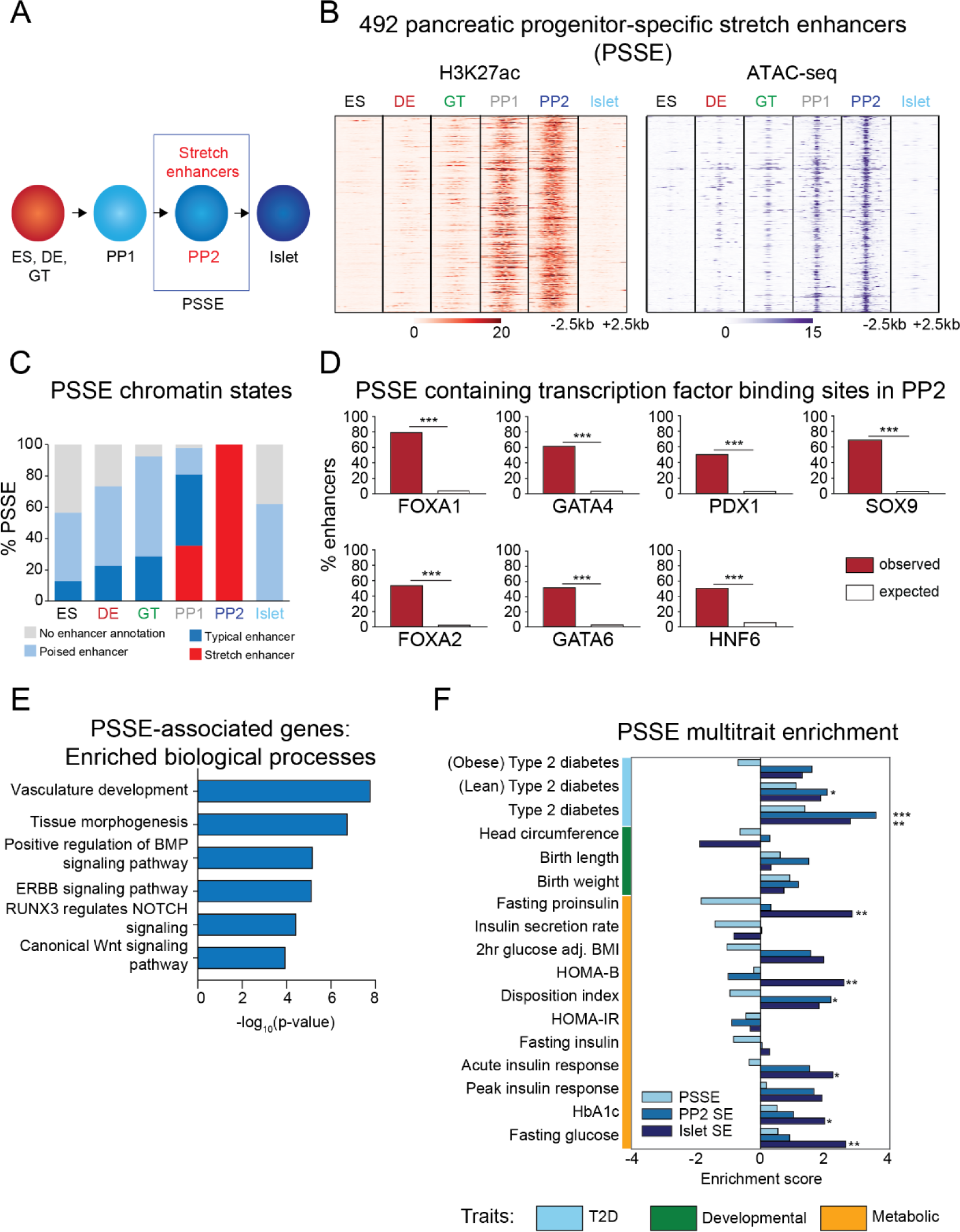
Candidate target genes of pancreatic progenitor-specific stretch enhancers regulate developmental processes. (**A**) Schematic illustrating identification of pancreatic progenitor-specific stretch enhancers (PSSE). (**B**) Heatmap showing density of H3K27ac ChIP-seq and ATAC-seq reads at PSSE, centered on overlapping H3K27ac and ATAC-seq peaks, respectively, and spanning 5 kb in ES, DE, GT, PP1, PP2, and islets. PSSE coordinates in Figure 2 – source data 1. (**C**) Percentage of PSSE exhibiting indicated chromatin states at defined developmental stages and in islets. (**D**) Percentage of PSSE overlapping with at least one ChIP-seq peak at PP2 for the indicated transcription factors. Enrichment analysis comparing observed and expected overlap based on random genomic regions of the same size and located on the same chromosome averaged over 10,000 iterations (*** p < 1 x 10^−4^; permutation test). (**E**) Gene ontology analysis for nearest expressed genes (fragments per kilobase per million fragments mapped (FPKM) ≥ 1 at PP2) to the 492 PSSE. See also Figure 2 – source data 2. (**F**) Enrichment (LD score regression coefficient z-scores) of T2D, developmental, and metabolic GWAS trait-associated variants at accessible chromatin sites in PSSE as compared with PP2 and islet stretch enhancers. Significant enrichment was identified within accessible chromatin at PP2 stretch enhancers for lean type 2 diabetes (Z = 2.06, *p = 3.94 × 10^−2^), at PP2 stretch enhancers for type 2 diabetes (Z = 3.57, ***p = 3.52 × 10^−4^), at islet stretch enhancers for type 2 diabetes (Z = 2.78, **p = 5.46 × 10^−3^), at islet stretch enhancers for fasting proinsulin levels (Z = 2.83, **p = 4.61 × 10^−3^), at islet stretch enhancers for HOMA-B (Z = 2.58, **p = 9.85 × 10^−3^), at PP2 stretch enhancers for disposition index (Z = 2.18, *p = 2.94 × 10^−2^), at islet stretch enhancers for acute insulin response (Z = 2.24, *p = 2.51 × 10^−2^), at islet stretch enhancers for HbA1c (Z = 1.98, *p = 4.72 × 10^−2^), and at islet stretch enhancers for fasting glucose levels (Z = 2.64, **p = 8.31 × 10^−3^). See also Figure 2 – source data 3 and Figure 2 – figure supplement 1.

To gain insight into the transcription factors that regulate PSSE, we conducted motif enrichment analysis of accessible chromatin sites within PSSE (**Figure 2 – figure supplement 1C**). Consistent with the activation of PSSE upon pancreas induction, motifs associated with transcription factors known to regulate pancreatic development (Conrad et al., 2014; Masui et al., 2007) were enriched, including FOXA (p = 1 × 10^−34^), PDX1 (p = 1 × 10^−30^), GATA (p = 1 × 10^−25^), ONECUT (p = 1 × 10^−17^), and RBPJ (p = 1 × 10^−14^), suggesting that pancreatic lineage-determining transcription factors activate PSSE. Analysis of the extent of PSSE overlap with ChIP-seq binding sites for FOXA1, FOXA2, GATA4, GATA6, PDX1, HNF6, and SOX9 at the PP2 stage substantiated this prediction (p < 1 × 10^−4^ for all transcription factors; permutation test; **Figure 2D**).

Annotation of biological functions of predicted target genes for PSSE (nearest gene with FPKM ≥ 1 at PP2 stage) revealed gene ontology terms related to developmental processes, such as tissue morphogenesis (p = 1 × 10^−7^) and vascular development (p = 1 × 10^−8^), as well as developmental signaling pathways, including BMP (p = 1 × 10^−5^), NOTCH (p = 1 × 10^−4^), and canonical Wnt signaling (p = 1 × 10^−4^; **Figure 2E** and **Figure 2 – source data 2**), which have demonstrated roles in pancreas morphogenesis and cell lineage allocation (Ahnfelt-Ronne et al., 2010; Li et al., 2015; Murtaugh, 2008; Sharon et al., 2019; Sui et al., 2013). Consistent with the temporal pattern of H3K27ac deposition at PSSE, transcript levels of PSSE-associated genes increased upon pancreatic lineage induction and peaked at the PP2 stage (p = 1.8 × 10^−8^; **Figure 2 – figure supplement 1D**). Notably, expression of these genes sharply decreased in islets (p < 2.2 × 10^−16^), underscoring the likely role of these genes in regulating pancreatic development but not mature islet function.

### Pancreatic progenitor-specific stretch enhancers are highly specific across T2D-relevant tissues and cell types

We next sought to understand the phenotypic consequences of PSSE activity in the context of T2D pathophysiology. Variants in accessible chromatin sites of PSSE genome-wide were enriched for T2D association (log enrichment = 2.85, 95% CI = <-20, 4.09). We determined enrichment of genetic variants for T2D-related quantitative endophenotypes within accessible chromatin sites of PSSE, as well as all pancreatic progenitor SE (not just progenitor-specific) and islet SE, using LD score regression (Bulik-Sullivan et al., 2015; Finucane et al., 2015). As expected based on prior observations (Parker et al., 2013; Pasquali et al., 2014), we observed enrichment (Z > 1.96) of variants associated with quantitative traits related to insulin secretion and beta cell function within islet SE, exemplified by fasting proinsulin levels, HOMA-B, and acute insulin response (Z = 2.8, Z = 2.6, and Z = 2.2, respectively; **Figure 2F**). Conversely, PSSE showed a trend toward depletion for these traits although the estimates were not significant. We further tested for enrichment in the proportion of variants in PSSE and islet SE nominally associated (p < 0.05) with beta cell function traits compared to background variants. There was significant enrichment of beta cell trait association among islet SE variants (χ^2^ test; p < 0.05 for all beta cell functional traits except for insulin secretion rate), but no corresponding enrichment for PSSE (**Figure 2 – source data 3**).

A prior study found that variants at the *LAMA1* locus had stronger effects on T2D risk among lean relative to obese cases (Perry et al., 2012). Since we identified a PSSE at the *LAMA1* locus, we postulated that variants in PSSE collectively might have differing impact on T2D risk in cases segregated by BMI. We therefore tested PSSE, as well as pancreatic progenitor SE and islet SE, for enrichment of T2D association using GWAS of lean and obese T2D (Perry et al., 2012), using LD score regression (Bulik-Sullivan et al., 2015; Finucane et al., 2015). We observed nominally significant enrichment of variants in pancreatic progenitor SE for T2D among lean cases (Z = 2.1). Variants in PSSE were mildly enriched for T2D among lean (Z = 1.1) and depleted among obese (Z = -0.70) cases, although neither estimate was significant. By comparison, islet SE showed positive enrichment for T2D among both lean (Z = 1.9) and obese cases (Z = 1.3; **Figure 2F**). Together, these results suggest that PSSE may affect T2D risk in a manner distinct from islet SE function.

Having observed little evidence for enrichment of PSSE variants for traits related to beta cell function, we asked whether the enrichment of PSSE for T2D-associated variants could be explained by PSSE activity in T2D-relevant tissues and cell types outside the pancreas. We assessed PSSE activity by measuring H3K27ac signal in 95 representative tissues and cell lines from the ENCODE and Epigenome Roadmap projects (Roadmap Epigenomics et al., 2015). Interestingly, there was group-wide specificity of PSSE to pancreatic progenitors relative to other cells and tissues including those relevant to T2D, such as adipose tissue, skeletal muscle, and liver (**Figure 2 – figure supplement 1E** and **Figure 2 – source data 4**). Since gene regulation in adipocyte precursors also contributes to T2D risk (Claussnitzer et al., 2014), we further examined PSSE specificity with respect to chromatin states during adipogenesis, using data from human adipose stromal cell differentiation stages (hASC1-4) (Mikkelsen et al., 2010; Varshney et al., 2017). PSSE exhibited virtually no active chromatin during adipogenesis (9, 8, 6, and 8 out of the 492 PSSE were active enhancers in hACS-1, hASC-2, hASC-3, and hASC-4, respectively; **Figure 2 – figure supplement 1F**). These findings identify PSSE as highly pancreatic progenitor-specific across T2D-relevant tissues and cell types.

### Identification of pancreatic progenitor-specific stretch enhancers harboring T2D-associated variants

Given the relative specificity of PSSE to pancreatic progenitors, we next sought to identify T2D-associated variants in PSSE at specific loci which may affect pancreatic development. We therefore identified variants in PSSE with evidence of T2D association (at p = 4.7 × 10^−6^) after correcting for the total number of variants in PSSE genome-wide (n = 10,738). In total there were 49 variants in PSSE with T2D association exceeding this threshold mapping to 11 loci (**Figure 3A**). This included variants at 9 loci with known genome-wide significant T2D association (*PROX1*, *ST6GAL1*, *SMARCAD1*, *XKR6*, *INS-IGF2*, *HMGA2*, *SMEK1*, *HMG20A*, and *LAMA1*), as well as at two previously unreported loci with sub-genome-wide significant association, *CRB2* and *PGM1*. To identify candidate target genes of the T2D-associated PSSE in pancreatic progenitors, we analyzed the expression of all genes within the same topologically associated domain (TAD) as the PSSE in PP2 cells and in primary human embryonic pancreas tissue (**Figure 3B** and **Figure 3 – figure supplement 1A**). These expressed genes are candidate effector transcripts of T2D-associated variants in pancreatic progenitors.

**Figure 3:**
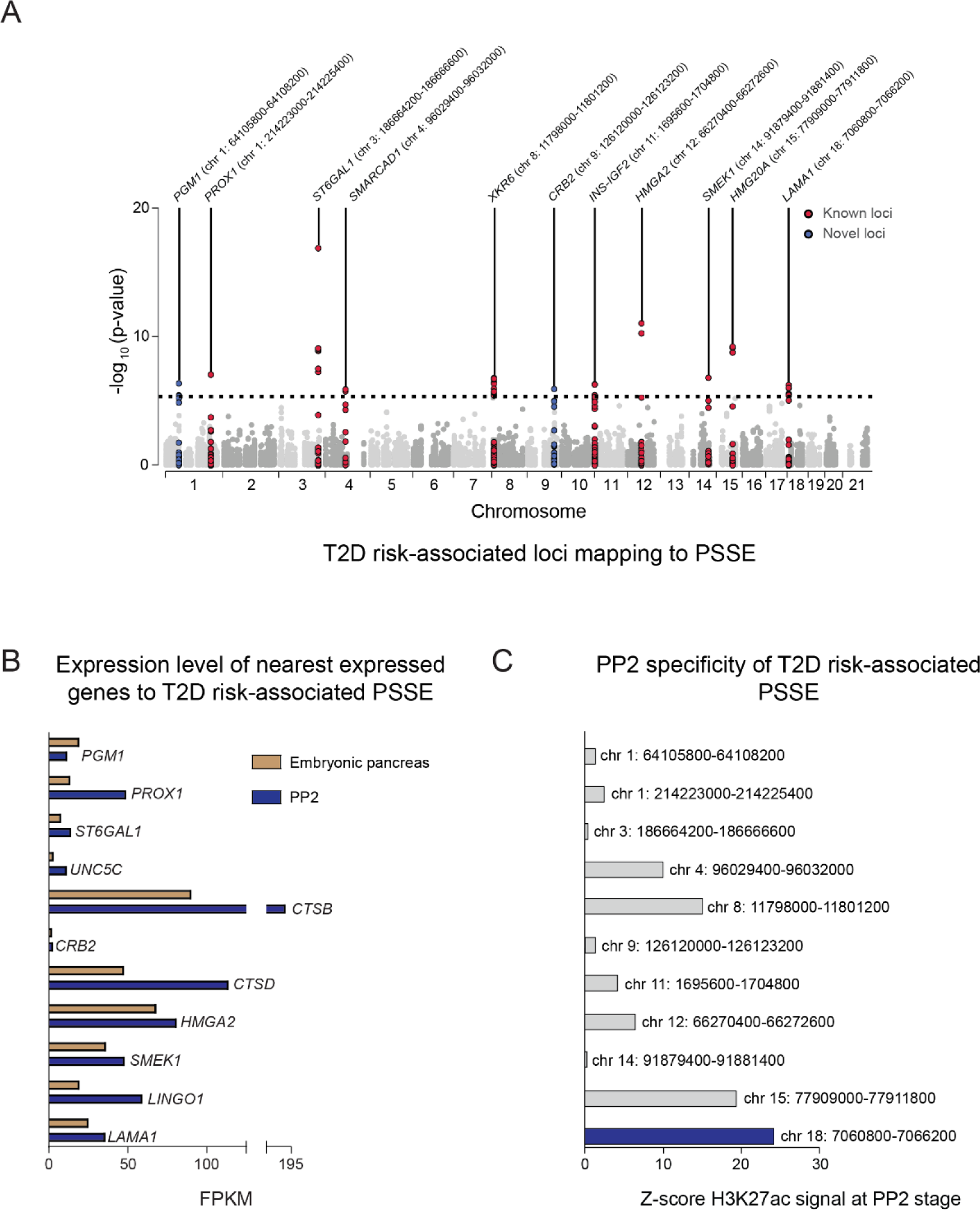
Identification of T2D risk variants associated with pancreatic progenitor-specific stretch enhancers. (**A**) Manhattan plot showing T2D association p-values (from Mahajan et al., 2018) for 10,738 variants mapping within PSSE. The dotted line shows the threshold for Bonferroni correction (p = 4.66 × 10^−6^). Novel loci identified with this threshold and mapping at least 500 kb away from a known locus are highlighted in blue. Chromosomal coordinates of T2D-associated PSSE are indicated. (**B**) mRNA levels (measured in fragments per kilobase per million fragments mapped (FPKM)) at PP2 (blue) and in human embryonic pancreas (54 and 58 days gestation, gold) of nearest expressed (FPMK ≥ 1) gene at PP2 for PSSE harboring T2D variants identified in Fig. 3A. (**C**) PP2 specificity of H3K27ac signal at PSSE harboring T2D variants identified in Fig. 3A. Z-score comparing H3K27ac signal at PP2 to H3K27ac signal in tissues and cell lines from the ENCODE and Epigenome Roadmap projects. See also Figure 3 – figure supplement 1.

As many pancreatic progenitor SE remain poised in mature islets (**Figure 2C**), we considered whether T2D-associated variants in PSSE could have gene regulatory function in islets that is re-activated in the disease state. We therefore assessed overlap of PSSE variants with accessible chromatin of islets from T2D donors (Khetan et al., 2018). None of the strongly T2D-associated variants in PSSE (p = 4.7 × 10^−6^) overlapped an islet accessible chromatin site in T2D islets, arguing against the relevance of PSSE in broadly regulating islet gene activity during T2D.

### A pancreatic progenitor-specific stretch enhancer at *LAMA1* harbors T2D risk variants and regulates *LAMA1* expression selectively in pancreatic progenitors

Variants in a PSSE at the *LAMA1* locus were associated with T2D at genome-wide significance (**Figure 3A**), and *LAMA1* was highly expressed in the human embryonic pancreas (**Figure 3B**). Furthermore, the activity of the PSSE at the *LAMA1* locus was almost exclusively restricted to pancreatic progenitors (**Figure 3 – figure supplement 1B,C**), and was further among the most progenitor-specific across all PSSE harboring T2D risk variants (**Figure 3C**). In addition, reporter gene assays in zebrafish embryos have shown that this enhancer drives gene expression specific to pancreatic progenitors *in vivo* (Cebola et al., 2015). We therefore postulated that the activity of T2D-associated variants within the *LAMA1* PSSE is relevant for gene regulation in pancreatic progenitors, and we sought to characterize the *LAMA1* PSSE in greater depth.

Multiple T2D-associated variants mapped within the *LAMA1* PSSE, and these variants were further in the 99% credible set in fine-mapping data from the DIAMANTE consortium (Mahajan et al., 2018) (**Figure 4A**). No other variants in the 99% credible set mapped in an accessible chromatin site active in islets from either non-diabetic or T2D samples. The PSSE is intronic to the *LAMA1* gene and contains regions of poised chromatin and TE at prior developmental stages (**Figure 4A**). Consistent with its stepwise genesis as a SE throughout development, regions of open chromatin within the *LAMA1* PSSE were already present at the DE and GT stages. Furthermore, pancreatic lineage-determining transcription factors, such as FOXA1, FOXA2, GATA4, GATA6, HNF6, SOX9, and PDX1, were all bound to the PSSE at the PP2 stage (**Figure 4B**). Among credible set variants in the *LAMA1* PSSE, rs10502347 overlapped an ATAC-seq peak as well as ChIP-seq sites for multiple pancreatic lineage-determining transcription factors. Additionally, rs10502347 directly coincided with a SOX9 footprint identified in ATAC-seq data from PP2 cells, and the T2D risk allele C is predicted to disrupt SOX9 binding (**Figure 4B**). Consistent with the collective endophenotype association patterns of PSSE (**Figure 2F**), rs10502347 showed no association with beta cell function (p = 0.81, 0.23, 0.46 for fasting proinsulin levels, HOMA-B, and acute insulin response, respectively; **Figure 4 – figure supplement 1A**). Thus, T2D variant rs10502347 is predicted to affect the binding of pancreatic transcription factors and does not appear to affect beta cell function.

**Figure 4:**
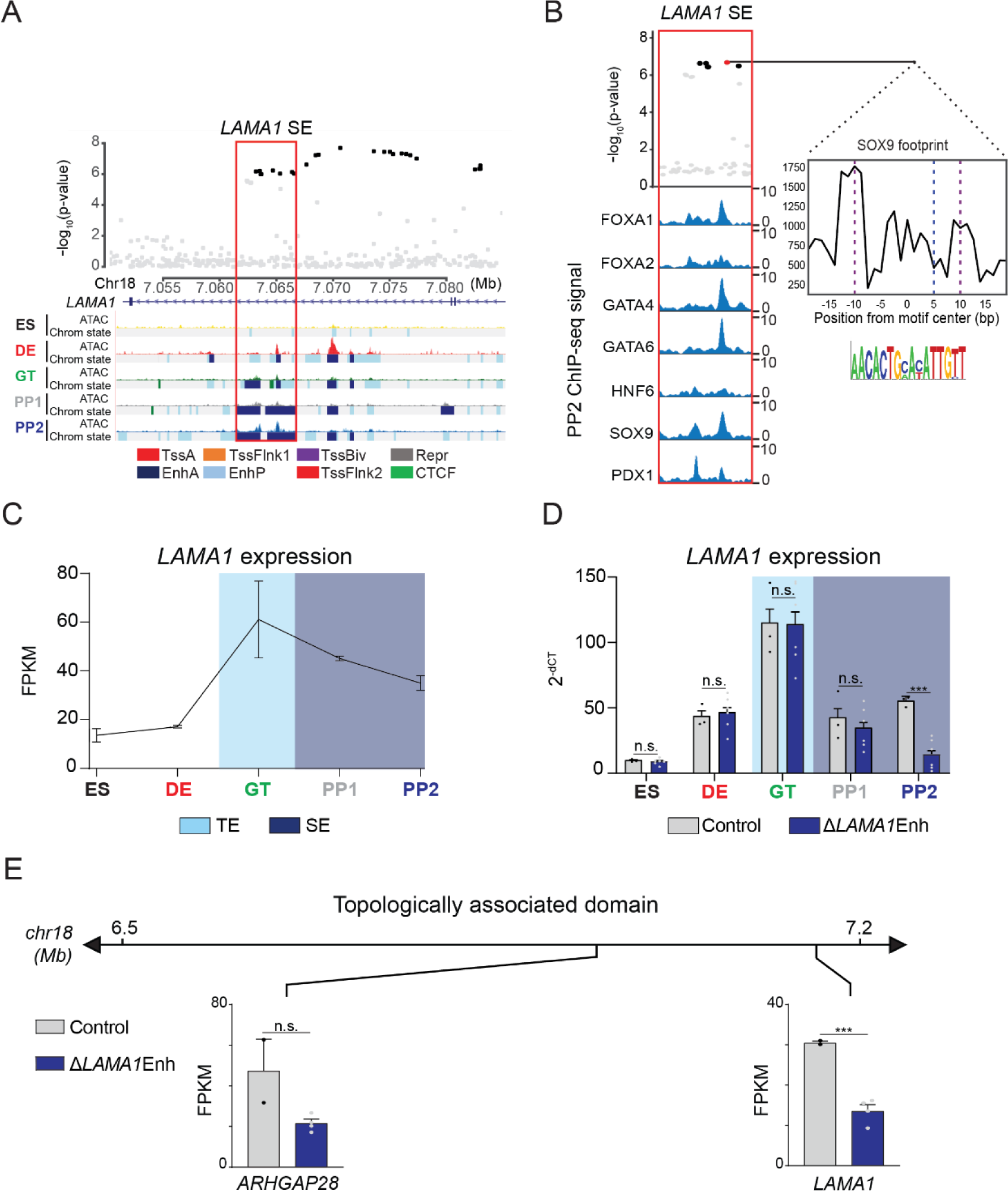
A T2D risk-associated *LAMA1* pancreatic progenitor-specific stretch enhancer regulates *LAMA1* expression specifically in pancreatic progenitors. (**A**) (Top) Locus plots showing T2D association p-values for variants in a 35 kb window (hg19 chr18:7,050,000-7,085,000) at the *LAMA1* locus and *LAMA1* PSSE (red box). Fine mapped variants within the 99% credible set for the *LAMA1* locus are colored black. All other variants are colored light gray. (Bottom) Chromatin states and ATAC-seq signal in ES, DE, GT, PP1, and PP2. TssA, active promoter; TssFlnk, flanking transcription start site; TssBiv, bivalent promoter; Repr, repressed; EnhA, active enhancer; EnhP, poised enhancer. (**B**) FOXA1, FOXA2, GATA4, GATA6, HNF6, SOX9, and PDX1 ChIP-seq profiles at the *LAMA1* PSSE in PP2. The variant rs10502347 (red) overlaps transcription factor binding sites and a predicted ATAC-seq footprint for the SOX9 sequence motif. Purple dotted lines indicate the core binding profile of the average SOX9 footprint genome-wide and the blue dotted line indicates the position of rs10502347 within the SOX9 motif. (**C**) *LAMA1* mRNA expression at each developmental stage determined by RNA-seq, measured in fragments per kilobase per million fragments mapped (FPKM). Data shown as mean ± S.E.M. (n = 3 replicates from independent differentiations). Light blue and purple indicate classification of the *LAMA1* PSSE as typical enhancer (TE) and stretch enhancer (SE), respectively. (**D**) *LAMA1* mRNA expression at each developmental stage determined by qPCR in control and Δ*LAMA1*Enh cells. Data are shown as mean ± S.E.M. (n = 3 replicates from independent differentiations for control cells. Δ*LAMA1*Enh cells represent combined data from 2 clonal lines with 3 replicates for each line from independent differentiations. n = 3 technical replicates for each sample; p = 0.319, 0.594, 0.945, 0.290, and < 1 × 10^−6^ for comparisons in ES, DE, GT, PP1, and PP2, respectively; student’s t-test, 2 sided; *** p < 0.001, n.s., not significant). Light blue and purple indicate classification of the *LAMA1* PSSE as TE and SE, respectively. Plotted points represent average of technical replicates for each differentiation. (**E**) mRNA expression determined by RNA-seq at PP2 of genes expressed in either control or Δ*LAMA1*Enh cells (FPKM ≥ 1 at PP2) and located within the same topologically associated domain as *LAMA1*. Data are shown as mean FPKM ± S.E.M. (n = 2 replicates from independent differentiations for control cells. Δ*LAMA1*Enh cells represent combined data from 2 clonal lines with 2 replicates for each line from independent differentiations. p adj. = 0.389 and 8.11 × 10^−3^ for *ARHGAP28* and *LAMA1*, respectively; DESeq2). See also Figure 4 – figure supplement 1.

Enhancers can control gene expression over large genomic distances, and therefore their target genes cannot be predicted based on proximity alone. To directly assess the function of the *LAMA1* PSSE in regulating gene activity, we utilized CRIPSR-Cas9-mediated genome editing to generate two independent clonal human hESC lines harboring homozygous deletions of the *LAMA1* PSSE (hereafter referred to as Δ*LAMA1*Enh; **Figure 4 – figure supplement 1B**). We examined *LAMA1* expression in Δ*LAMA1*Enh compared to control cells throughout stages of pancreatic differentiation. Consistent with the broad expression of *LAMA1* across developmental and mature tissues, control cells expressed *LAMA1* at all stages (**Figure 4C**). *LAMA1* was expressed at similar levels in Δ*LAMA1*Enh and control cells at early developmental stages, but was significantly reduced in PP2 cells derived from Δ*LAMA1*Enh clones (p = 0.319, 0.594, 0.945, 0.290, and < 1 × 10^−6^ for comparisons in ES, DE, GT, PP1, and PP2, respectively; **Figure 4D**). To next investigate whether the *LAMA1* PSSE regulates other genes at this locus, we examined expression of genes mapping in the same TAD. *ARHGAP28* was the only other expressed gene within the TAD, and albeit not significantly different from controls (p.adj > 0.05), showed a trend toward lower expression in Δ*LAMA1*Enh PP2 cells (**Figure 4E**), raising the possibility that *ARHGAP28* is an additional target gene of the *LAMA1* PSSE. Together, these results demonstrate that while *LAMA1* itself is broadly expressed across developmental stages, the T2D-associated PSSE regulates *LAMA1* expression specifically in pancreatic progenitors.

To determine whether deletion of the *LAMA1* PSSE affects pancreatic development, we generated PP2 stage cells from Δ*LAMA1*Enh and control hESC lines and analyzed pancreatic cell fate commitment by flow cytometry and immunofluorescence staining for PDX1 and NKX6.1 (**Figure 4 – figure supplement 1C,D**). At the PP2 stage, Δ*LAMA1*Enh and control cultures contained similar percentages of PDX1- and NKX6.1-positive cells. Furthermore, mRNA expression of *PDX1*, *NKX6.1*, *PROX1*, *PTF1A*, and *SOX9* was either unaffected or only minimally reduced (p adj. = 3.56 × 10^−2^, 0.224, 0.829, 8.14 × 10^−2^, and 0.142, for comparisons of *PDX1*, *NKX6.1*, *PROX1*, *PTF1A*, and *SOX9* expression, respectively; **Figure 4 – figure supplement 1E**), and the overall gene expression profiles were similar in Δ*LAMA1*Enh and control PP2 cells (**Figure 4 – figure supplement 1F** and **Figure 4 – source data 1,2**). These findings indicate that *in vitro* pancreatic lineage induction is unperturbed in Δ*LAMA1*Enh cells exhibiting reduced *LAMA1* expression.

### Pancreatic progenitor-specific stretch enhancers at the *CRB2* and *PGM1* loci harbor T2D-associated variants

Multiple variants with evidence for T2D association in PSSE mapped outside of known risk loci, such as those mapping to *CRB2* and *PGM1* (**Figure 3A**). As with the *LAMA1* PSSE, PSSE harboring variants at *CRB2* and *PGM1* were intronic to their respective genes, harbored ATAC-seq peaks, and bound pancreatic lineage-determining transcription factors FOXA1, FOXA2, GATA4, GATA6, HNF6, SOX9, and PDX1 (**Figure 5A,B** and **Figure 5 – figure supplement 1A,B**). Compared to the *LAMA1* PSSE, *CRB2* and *PGM1* PSSE were less specific to pancreatic progenitors and exhibited significant H3K27ac signal in several other tissues and cell types, most notably brain, liver, and the digestive tract (**Figure 5 – figure supplement 1C,D**).

**Figure 5:**
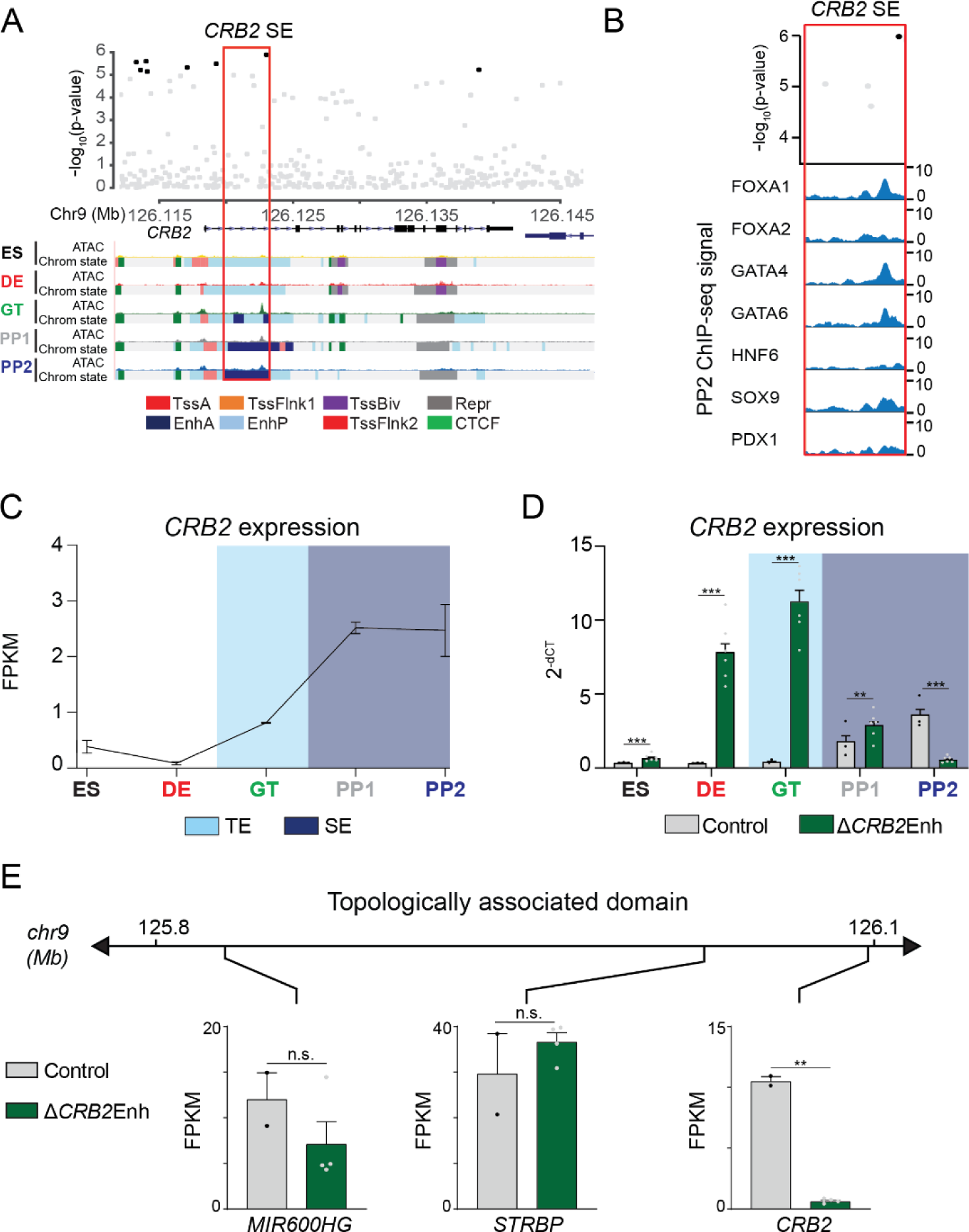
A T2D risk-associated *CRB2* pancreatic progenitor-specific stretch enhancer regulates *CRB2* expression specifically in pancreatic progenitors. (**A**) (Top) Locus plots showing T2D association p-values for variants in a 35 kb window (hg19 chr9:126,112,000-126,147,000) at the *CRB2* locus and *CRB2* PSSE (red box). Fine mapped variants within the 99% credible set for the novel *CRB2* locus are colored black. All other variants are colored light gray. (Bottom) Chromatin states and ATAC-seq signal in ES, DE, GT, PP1, and PP2. TssA, active promoter; TssFlnk, flanking transcription start site; TssBiv, bivalent promoter; Repr, repressed; EnhA, active enhancer; EnhP, poised enhancer. (**B**) FOXA1, FOXA2, GATA4, GATA6, HNF6, SOX9, and PDX1 ChIP-seq profiles at the *CRB2* PSSE in PP2. The variant rs2491353 (black) overlaps with transcription factor binding sites. (**C**) *CRB2* mRNA expression at each developmental stage determined by RNA-seq, measured in fragments per kilobase per million fragments mapped (FPKM). Data shown as mean ± S.E.M. (n = 3 replicates from independent differentiations). Light blue and purple indicate classification of the *CRB2* PSSE as typical enhancer (TE) and stretch enhancer (SE), respectively. Plotted points represent average of technical replicates for each differentiation. (**D**) *CRB2* mRNA expression at each developmental stage determined by qPCR in control and Δ*CRB2*Enh cells. Data are shown as mean ± S.E.M. (n = 3 replicates from independent differentiations for control cells. Δ*CRB2*Enh cells represent combined data from 2 clonal lines with 3 replicates for each line from independent differentiations. n = 3 technical replicates for each sample; p = 7.03 × 10^−4^, < 1 × 10^−6^, < 1 × 10^−6^, 1.46 × 10^−2^, and < 1 × 10^−6^ for comparisons in ES, DE, GT, PP1, and PP2, respectively; student’s t-test, 2 sided; *** p < 0.001 ** p < 0.01). Light blue and purple indicate classification of the *CRB2* PSSE as TE and SE, respectively. (**E**) mRNA expression determined by RNA-seq at PP2 of genes expressed in either control or Δ*CRB2*Enh cells (FPKM ≥ 1 at PP2) and located within the same topologically associated domain as *CRB2*. Data are shown as mean FPKM ± S.E.M. (n = 2 replicates from independent differentiations for control cells. Δ*CRB2*Enh cells represent combined data from 2 clonal lines with 2 replicates for each line from independent differentiations. p adj. = 0.158, 1.00, and 3.51 × 10^−3^, for *MIR600HG*, *STRBP*, and *CRB2*, respectively; DESeq2; **p < 0.01, n.s., not significant). See also Figure 5 – figure supplement 1 and 2.

CRB2 is a component of the Crumbs protein complex involved in the regulation of cell polarity and neuronal, heart, retinal, and kidney development (Alves et al., 2013; Bulgakova & Knust, 2009; Dudok et al., 2016; Jimenez-Amilburu & Stainier, 2019; Slavotinek et al., 2015). However, its role in pancreatic development is unknown. To determine whether the *CRB2* PSSE regulates *CRB2* expression in pancreatic progenitors, we generated two independent hESC clones with homozygous deletions of the *CRB2* PSSE (hereafter referred to as Δ*CRB2*Enh; **Figure 5 – figure supplement 2A**) and performed pancreatic differentiation of Δ*CRB2*Enh and control hESC lines. In control cells, *CRB2* was first expressed at the GT stage and increased markedly at the PP1 stage (**Figure 5C**). This pattern of *CRB2* expression is consistent with H3K27ac deposition at the *CRB2* PSSE in GT stage cells and classification as a SE at the PP1 and PP2 stages (**Figure 5A** and **Figure 5 – figure supplement 1C**). In Δ*CRB2*Enh cells, we observed upregulation of *CRB2* expression at earlier developmental stages, in particular at the DE and GT stages (p < 1 × 10^−6^ at both stages; **Figure 5D**), suggesting that the *CRB2* PSSE may be associated with repressive transcriptional complexes prior to pancreas induction. At the PP2 stage, *CRB2* expression was significantly reduced in Δ*CRB2*Enh cells (p adj. = 3.51 × 10^−3^; **Figure 5D**), whereas the expression of other genes in the same TAD was not affected (p adj. ≥ 0.05; **Figure 5E**). Thus, the *CRB2* PSSE specifically regulates *CRB2* and is required for *CRB2* expression in pancreatic progenitors.

Phenotypic characterization of PP2 stage Δ*CRB2*Enh cells revealed similar percentages of PDX1-and NKX6.1-positive cells as in control cells (**Figure 5 – figure supplement 2B,C**). The expression of pancreatic transcription factors and global gene expression profiles were also similar in Δ*CRB2*Enh and control PP2 cells (**Figure 5 – figure supplement 2D,E** and **Figure 5 – source data 1**). Thus, similar to *LAMA1* PSSE deletion, *CRB2* PSSE deletion does not overtly impair pancreatic lineage induction in the *in vitro* hESC differentiation system.

### *lama1* and *crb2* zebrafish morphants display annular pancreas and decreased beta cell mass

Based on their classification as extracellular matrix and cell polarity proteins, respectively, laminin (encoded by *LAMA1*) and CRB2 are predicted to regulate processes related to tissue morphogenesis, such as cell migration, tissue growth, and cell allocation within the developing organ. Furthermore, PSSE in general were enriched for proximity to genes involved in tissue morphogenesis (**Figure 2E**), suggesting that T2D risk variants acting within PSSE could have roles in pancreas morphogenesis. Since cell migratory processes and niche-specific signaling events are not fully modeled during hESC differentiation, we reasoned that the *in vitro* pancreatic differentiation system might not be suitable for studying laminin and CRB2 function in pancreatic development.

To circumvent these limitations, we employed zebrafish as an *in vivo* vertebrate model to study the effects of reduced *lama1* and *crb2* levels on pancreatic development. The basic organization and cell types in the pancreas as well as the genes regulating endocrine and exocrine pancreas development are highly conserved between zebrafish and mammals (Dong et al., 2008; Field et al., 2003; Kimmel et al., 2015). To analyze pancreatic expression of laminin and Crb proteins, we used *TgBAC(pdx1:eGFP)*^bns13^ embryos, in which eGFP marks the duodenum and developing pancreas, consistent with mammalian *Pdx1* expression (**Figure 6 – Figure supplement 1A**). At 48 hours post-fertilization (hpf), laminin was detected adjacent to pancreatic cells expressing *pdx1:eGFP* (**Figure 6 – Figure supplement 1A**, yellow arrow), whereas Crb was localized in *pdx1:eGFP* pancreatic cells (**Figure 6 – Figure supplement 1B**). Within the developing foregut region, Crb expression was exclusive to the pancreatic anlage (**Figure 6 – Figure supplement 1B**).

To determine the respective functions of *lama1* and *crb2* in pancreatic development, we performed knockdown experiments using anti-sense morpholinos directed against *lama1* and the two zebrafish *crb2* genes, *crb2a* and *crb2b* (Omori & Malicki, 2006; Pollard et al., 2006). We determined effects of these knockdowns in the *Tg(ptf1a:eGFP)^jh1^* embryos to visualize the acinar pancreas, which comprises the majority of the organ. Consistent with prior studies (Pollard et al., 2006), *lama1* morphants exhibited reduced body size and other gross anatomical defects at 78 hpf, whereas *crb2a/b* morphants appeared grossly normal. Both *lama1* and *crb2a/b* morphants displayed an annular pancreas (15 out of 34 *lama1* and 27 out of 69 *crb2a/b* morphants) characterized by pancreatic tissue partially or completely encircling the duodenum (**Figure 6A-D**), a phenotype indicative of impaired migration of pancreatic progenitors during pancreas formation. These findings suggest that both *lama1* and *crb2a/b* control cell migratory processes during early pancreatic development and that reduced levels of *lama1* or *crb2a/b* impair pancreas morphogenesis.

**Figure 6:**
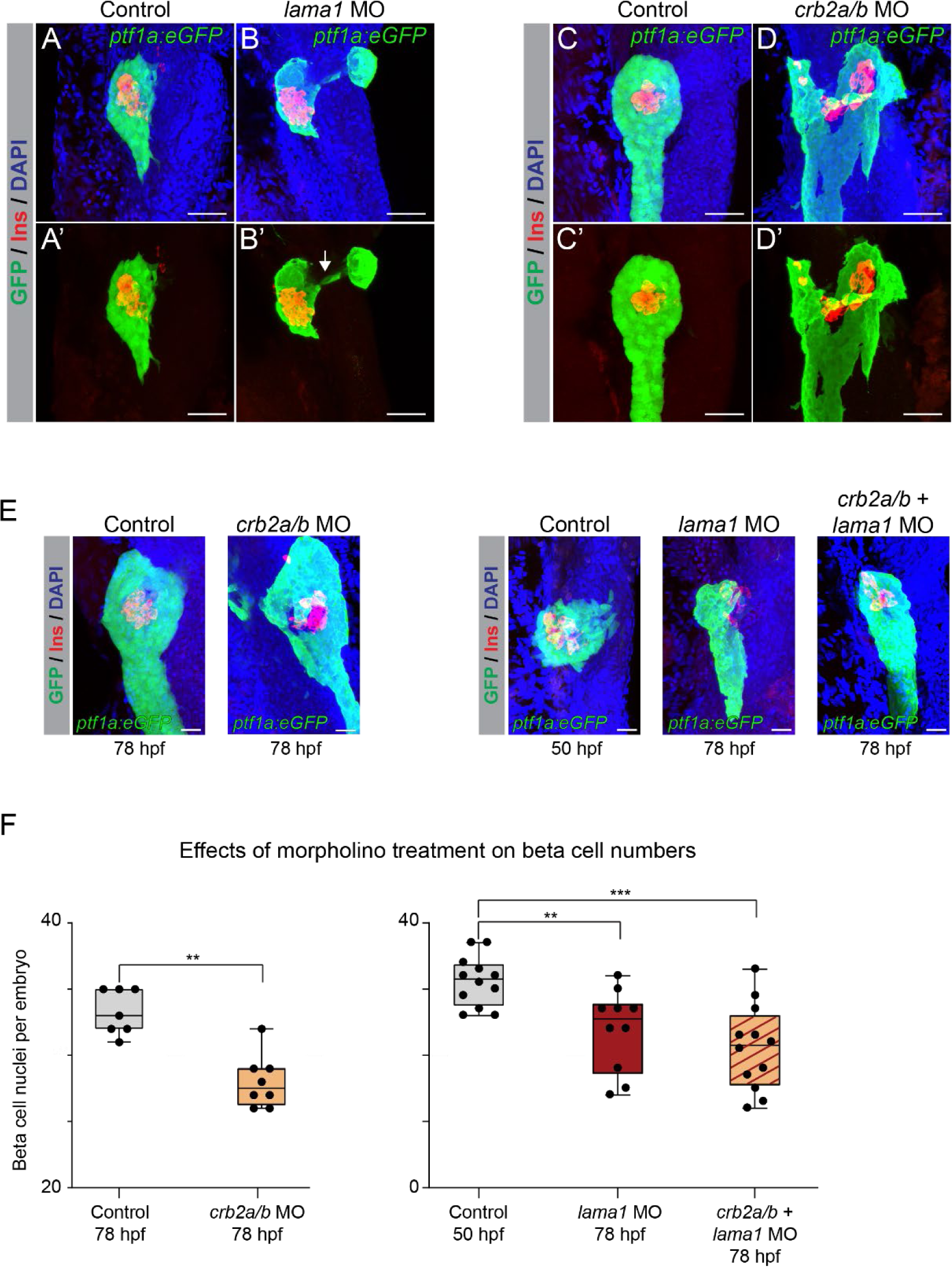
*lama1* and *crb2* regulate pancreas morphogenesis and beta cell differentiation. (**A,B**) Representative 3-dimensional renderings of *Tg(ptf1a:eGFP)* control zebrafish embryos (**A,A’**) and *lama1* morphants (**B,B’**) stained with DAPI (nuclei, blue) and antibody against insulin (red); n ≥ 15 embryos per condition. To account for reduced acinar pancreas size in *lama1* morphants, control embryos were imaged at 50 hours post fertilization (hpf) and *lama1* morphants at 78 hpf. 15 out of 34 *lama1* morphants displayed an annular pancreas with two acinar pancreas domains (green) connected behind the presumptive intestine (**B’**, white arrow). Scale bar, 40 µM. (**C,D**) Representative 3 dimensional renderings of 78 hpf *Tg(ptf1a:eGFP)* control zebrafish embryos (**C,C’**) and *crb2a/b* morphants (**D,D’**) stained with DAPI (nuclei, blue) and antibodies against insulin (red); n ≥ 15 embryos per condition. 27 out of 69 *crb2a/b* morphants displayed an annular pancreas with the acinar pancreas (green) completely surrounding the presumptive intestine. Scale bar, 40 µM. (**E**) Representative 3-dimensional renderings of *Tg(ptf1a:eGFP)* control zebrafish embryos and *crb2a/b, lama1*, or *crb2a/b + lama1* morphants stained with DAPI (nuclei, blue) and antibody against insulin (red). All embryos were imaged at 78 hpf except for controls to *lama1* and *crb2a/b + lama1* morphants, which were imaged at 50 hpf to account for reduced acinar pancreas size of *lama1* morphants. Scale bar, 20 µM. (**F**) Quantification of beta (insulin^+^) cell nuclei per embryo from experiment in (E). p adj. = 4.0 × 10^−3^, 8.0 × 10^−3^, and 2.0 × 10^−4^ for comparison of hfp 78 control (n = 7 embryos) to hfp 78 *crb2a/b* (n = 8), hpf 50 control (n = 12) to hpf 78 *lama1* (n = 10), or *crb2a/b + lama1* (n = 12) morphants, respectively; ANOVA-Dunnett’s multiple comparison test; *** p < 0.001 ** p < 0.01. 5 out of 8 *crb2a/b*, 3 out of 10 *lama1*, and 9 out of 12 *crb2a/b + lama1* morphants displayed an annular pancreas. See also Figure 6 – figure supplement 1 and 2.

To gain insight into the effects of *lama1* and *crb2a/b* knockdown on pancreatic endocrine cell development, we examined beta cell numbers (insulin^+^ cells) at 78 hpf. We also evaluated potential synergistic effects of combined *lama1* and *crb2a/b* knockdown. To account for the reduction in body and pancreas size in *lama1* morphants, we compared cell numbers in 78 hpf *lama1* morphants with 50 hpf control embryos, which have a similarly sized acinar compartment as 78 hpf *lama1* morphants. Beta cell numbers were significantly reduced in both *lama1* and *crb2a/b* morphants (p = 8.0 × 10^−3^ and 4.0 × 10^−3^ for comparisons of *lama1* and *crb2a/b* morphants, respectively; **Figure 6E,F**), showing that reduced *lama1* and *crb2a/b* levels impair beta cell development. Although not significant, morphants with a combined knockdown of *lama1* and *crb2a/b* had a trend toward lower beta cell numbers than individual morphants, suggestive of additive effects (p = 0.42; **Figure 6F**). Furthermore, we found that nearly all *lama1*, *crb2a/b,* and combined *lama1* and *crb2a/b* morphants without an annular pancreas had reduced beta cell numbers, indicating independent roles of *lama1* and *crb2* in pancreas morphogenesis and beta cell differentiation. Finally, to investigate the contributions of individual *crb2* genes to the observed phenotype, we performed knockdown experiments using morpholinos against *crb2a* and *crb2b* alone. Only *crb2b* morphants showed a significant reduction in beta cell numbers (p = 4.4 × 10^−2^; **Figure 6 – Figure supplement 2**), suggesting that *crb2b* is the predominant *crb2* gene required for beta cell development. Combined, these findings demonstrate that *lama1* and *crb2* are regulators of pancreas morphogenesis and beta cell development *in vivo*.

## DISCUSSION

In this study, we identify T2D-associated variants localized within chromatin active in pancreatic progenitors but not islets or other T2D-relevant tissues, suggesting a novel mechanism whereby a subset of T2D risk variants specifically alters pancreatic developmental processes. We link T2D-associated enhancers active in pancreatic progenitors to the regulation of *LAMA1* and *CRB2* and demonstrate a functional requirement in zebrafish for *lama1* and *crb2* in pancreas morphogenesis and endocrine cell formation. Furthermore, we provide a curated list of T2D risk-associated enhancers and candidate effector genes for further exploration of how the regulation of developmental processes in the pancreas can predispose to T2D.

Our analysis identified eleven loci where T2D-associated variants mapped in SE specifically active in pancreatic progenitors. Among these loci was *LAMA1*, which has stronger effects on T2D risk in lean compared to obese individuals (Perry et al., 2012). We also found evidence that variants in PSSE collectively have stronger enrichment for T2D in lean individuals, although the small number of PSSE and limited sample size of the BMI-stratified T2D genetic data prohibits a more robust comparison. There was also a notable lack of enrichment among PSSE variants for association with traits related to insulin secretion and beta cell function. If T2D-associated variants in PSSE indeed confer diabetes susceptibility by affecting beta cell development, the question arises as to why variants associated with traits related to beta cell function are not enriched within PSSE. As genetic association studies of endophenotypes are based on data from non-diabetic subjects, a possible explanation is that variants affecting beta cell developmental processes have no overt phenotypic effect under physiological conditions and contribute to T2D pathogenesis only during the disease process.

Since the genomic position of enhancers and transcription factor binding sites is not well conserved between species (Villar et al., 2015), a human cell model is necessary to identify target genes of enhancers associated with disease risk. By employing enhancer deletion in hESCs, we demonstrate that T2D-associated PSSE at the *LAMA1* and *CRB2* loci regulate *LAMA1* and *CRB2*, respectively, and establish *LAMA1* and *CRB2* as the predominant target gene of their corresponding PSSE within TAD boundaries. By analyzing *LAMA1* and *CRB2* expression throughout the pancreatic differentiation time course, we show that the identified PSSE control *LAMA1* and *CRB2* expression in a temporal manner consistent with the activation pattern of their associated PSSE. While the specific T2D-relevant target genes of the majority of T2D-associated PSSE remain to be identified, it is notable that several are localized within TADs containing genes encoding transcriptional regulators. These include *PROX1* and *GATA4*, which are known to regulate pancreatic development (Shi et al., 2017; Tiyaboonchai et al., 2017; Westmoreland et al., 2012), as well as *HMGA2* and *BCL6* with unknown functions in the pancreas. Our catalogue of T2D-associated PSSE provides a resource to fully characterize the gene regulatory program associated with developmentally mediated T2D risk in the pancreas. Our finding that predicted target genes of PSSE are similarly expressed in hESC-derived pancreatic progenitors and primary human embryonic pancreas (**Figure 3B** and **Figure 3 – figure supplement 1A**) further underscores the utility of the hESC-based system for these studies.

In the embryo, endocrine cells differentiate by delaminating from a polarized epithelium of progenitors governed by local cell-cell and cell-matrix signaling events (Mamidi et al., 2018). These processes are not well-recapitulated in the hESC-based pancreatic differentiation system, highlighting a limitation of this system for studying the function of laminin and CRB2, which are mediators of mechanical signals within an epithelium. Therefore, we analyzed their function in zebrafish as an *in vivo* model. We show that *lama1* or *crb2* knockdown leads to an annular pancreas and reduced beta cell numbers. The beta cell differentiation defect was also evident in embryos not displaying an annular pancreas, suggesting independent mechanisms.

Consistent with our findings in *lama1* morphants, culture of pancreatic progenitors on laminin-based substrates promotes endocrine cell differentiation (Mamidi et al., 2018). During *in vivo* pancreatic development, endothelial cells are an important albeit not the only source of laminin in the pancreas (Heymans et al., 2019; Mamidi et al., 2018; Nikolova et al., 2006). While we do not know the respective contributions of endothelial cell- and pancreatic progenitor cell-derived laminin to the phenotype of *lama1* morphants, the T2D-associated *LAMA1* PSSE is not active in endothelial cells (**Figure 3 – figure supplement 1C**). Furthermore, we found no other T2D-associated variants at the *LAMA1* locus mapping in endothelial cell enhancers or accessible chromatin sites in islets, suggesting that T2D risk is linked to *LAMA1* regulation in pancreatic progenitors.

Similar to laminin, CRB2 has been shown to regulate mechanosignaling (Varelas et al., 2010). Our observation that Crb expression in the developing foregut is specific to *pdx1*-expressing pancreatic epithelial cells suggests that the phenotype of *crb2* morphants reflects a progenitor-autonomous role of Crb2. The similarity in pancreatic phenotype between *lama1* or *crb2* morphants raises the possibility that signals from laminin and Crb2 could converge on the same intracellular pathways in pancreatic progenitors.

Our findings suggest that variation in gene regulation during pancreatic development can predispose to T2D later in life. Several lines of evidence support the concept of a developmental impact on T2D risk. First, human genetic studies have shown a strong correlation between birth weight and adult cardiometabolic traits and disease (Horikoshi et al., 2016). Second, epidemiological studies provide evidence that offspring of mothers who were pregnant during a famine have a higher prevalence of T2D (Lumey et al., 2015). This phenomenon has been experimentally reproduced in rodents, where maternal malnutrition has been shown to cause reduced beta cell mass at birth and to render beta cells more prone to failure under stress (Nielsen et al., 2014). Together, our results provide a strong rationale for further exploration of how genetic variants affecting developmental gene regulation in the pancreas contribute to T2D risk.

## MATERIAL AND METHODS

### Maintenance and differentiation of CyT49 hESCs

Genomic and gene expression analyses (ChIP-seq, ATAC-seq, RNA-seq) for generation of chromatin maps and target gene identification were performed in CyT49 hESCs (male). Propagation of CyT49 hESCs was carried out by passing cells every 3 to 4 days using Accutase™ (eBioscience) for enzymatic cell dissociation, and with 10% (v/v) human AB serum (Valley Biomedical) included in the hESC media the day of passage. hESCs were seeded into tissue culture flasks at a density of 50,000 cells/cm^2^. hESC research was approved by the University of California, San Diego, Institutional Review Board and Embryonic Stem Cell Research oversight committee.

Pancreatic differentiation was performed as previously described (Schulz et al., 2012; Wang et al., 2015; Xie et al., 2013). Briefly, a suspension-based culture format was used to differentiate cells in aggregate form. Undifferentiated aggregates of hESCs were formed by re-suspending dissociated cells in hESC maintenance medium at a concentration of 1 × 10^6^ cells/mL and plating 5.5 mL per well of the cell suspension in 6-well ultra-low attachment plates (Costar). The cells were cultured overnight on an orbital rotator (Innova2000, New Brunswick Scientific) at 95 rpm. After 24 hours the undifferentiated aggregates were washed once with RPMI medium and supplied with 5.5 mL of day 0 differentiation medium. Thereafter, cells were supplied with the fresh medium for the appropriate day of differentiation (see below). Cells were continually rotated at 95 rpm, or 105 rpm on days 4 through 8, and no media change was performed on day 10. Both RPMI (Mediatech) and DMEM High Glucose (HyClone) medium were supplemented with 1X GlutaMAX™ and 1% penicillin/streptomycin. Human activin A, mouse Wnt3a, human KGF, human noggin, and human EGF were purchased from R&D systems. Other added components included FBS (HyClone), B-27® supplement (Life Technologies), Insulin-Transferrin-Selenium (ITS; Life Technologies), TGFβ R1 kinase inhibitor IV (EMD Bioscience), KAAD-Cyclopamine (KC; Toronto Research Chemicals), and the retinoic receptor agonist TTNPB (RA; Sigma Aldrich). Day-specific differentiation media formulations were as follows:

Days 0 and 1: RPMI + 0.2% (v/v) FBS, 100 ng/mL Activin, 50 ng/mL mouse Wnt3a, 1:5000 ITS. Days 1 and 2: RPMI + 0.2% (v/v) FBS, 100ng/mL Activin, 1:5000 ITS
Days 2 and 3: RPMI + 0.2% (v/v) FBS, 2.5 mM TGFβ R1 kinase inhibitor IV, 25ng/mL KGF, 1:1000 ITS
Days 3 – 5: RPMI + 0.2% (v/v) FBS, 25ng/mL KGF, 1:1000 ITS
Days 5 – 8: DMEM + 0.5X B-27® Supplement, 3 nM TTNPB, 0.25 mM KAAD-Cyclopamine, 50ng/mL Noggin
Days 8 – 10: DMEM/B-27, 50ng/mL KGF, 50ng/mL EGF

Cells at D0 correspond to the embryonic stem cell (ES) stage, cells at D2 correspond to the definitive endoderm (DE) stage, cells at D5 correspond to the gut tube (GT) stage, cells at D7 correspond to the early pancreatic progenitor (PP1) stage, and cells at D10 correspond to the late pancreatic progenitor (PP2) stage.

### Maintenance and differentiation of H1 hESCs

*ΔLAMA1Enh* and *ΔCRB2Enh* clonal lines were derived by targeting H1 hESCs (male). Cells were maintained and differentiated as described with some modifications (Jin et al., 2019; Rezania et al., 2014). In brief, hESCs were cultured in mTeSR1 media (Stem Cell Technologies) and propagated by passaging cells every 3 to 4 days using Accutase (eBioscience) for enzymatic cell dissociation. hESC research was approved by the University of California, San Diego, Institutional Review Board and Embryonic Stem Cell Research Oversight Committee.

For differentiation, cells were dissociated using Accutase for 10 min, then reaggregated by plating the cells at a concentration of ∼5.5 e6 cells/well in a low attachment 6-well plate on an orbital shaker (100 rpm) in a 37 °C incubator. The following day, undifferentiated cells were washed in base media (see below) and then differentiated using a multi-step protocol with stage-specific media and daily media changes.

All stage-specific base media were comprised of MCDB 131 medium (Thermo Fisher Scientific) supplemented with NaHCO3, GlutaMAX, D-Glucose, and BSA using the following concentrations:

Stage 1/2 medium: MCDB 131 medium, 1.5 g/L NaHCO3, 1X GlutaMAX, 10 mM D-Glucose, 0.5% BSA
Stage 3/4 medium: MCDB 131 medium, 2.5 g/L NaHCO3, 1X GlutaMAX, 10 mM D-glucose, 2% BSA

Media compositions for each stage were as follows:

Stage 1 (day 0-2): base medium, 100 ng/ml Activin A, 25 ng/ml Wnt3a (day 0). Day 1-2: base medium, 100 ng/ml Activin A
Stage 2 (day 3-5): base medium, 0.25 mM L-Ascorbic Acid (Vitamin C), 50 ng/mL FGF7
Stage 3 (day 6-7): base medium, 0.25 mM L-Ascorbic Acid, 50 ng/mL FGF7, 0.25 µM SANT-1, 1 µM Retinoic Acid, 100 nM LDN193189, 1:200 ITS-X, 200 nM TPB
Stage 4 (day 8-10): base medium, 0.25 mM L-Ascorbic Acid, 2 ng/mL FGF7, 0.25 µM SANT-1, 0.1 µM Retinoic Acid, 200 nM LDN193189, 1:200 ITS-X, 100nM TPB

Cells at D0 correspond to the embryonic stem cell (ES) stage, cells at D3 correspond to the definitive endoderm (DE) stage, cells at D6 correspond to the gut tube (GT) stage, cells at D8 correspond to the early pancreatic progenitor (PP1) stage, and cells at D11 correspond to the late pancreatic progenitor (PP2) stage.

### Generation of Δ*LAMA1*Enh and Δ*CRB2*Enh hESC lines

To generate clonal homozygous *LAMA1*Enh and *CRB2*Enh deletion hESC lines, sgRNAs targeting each relevant enhancer were designed and cloned into Px333-GFP, a modified version of Px333 (Addgene, #64073). The plasmid was transfected into H1 hESCs with XtremeGene 9 (Roche). 24 hours later, 8000 GFP^+^ cells were sorted into a well of six-well plate. Individual colonies that emerged within 5-7 days after transfection were subsequently transferred manually into 48-well plates for expansion, genomic DNA extraction, PCR genotyping, and Sanger sequencing. sgRNA oligos are listed below.

*LAMA1*Enh Upstream Guide: GTCAAATTGCTATAACACGG
*LAMA1*Enh Downstream Guide: CCACTTTAAGTATCTCAGCA
*CRB2*Enh Upstream Guide: ATACAAAGCACGTGAGA
*CRB2*Enh Downstream Guide: GAATGCGGATGACGCCTGAG

### Human tissue

Human embryonic pancreas tissue was obtained from the Birth Defects Research Laboratory of the University of Washington. Studies for use of embryonic human tissue were approved by the Institutional Review Board of the University of California, San Diego. A pancreas from a 54 and 58 day gestation embryo each were pooled for RNA-seq analysis.

### Zebrafish husbandry

Adult zebrafish and embryos were cared for and maintained under standard conditions. All research activity involving zebrafish was reviewed and approved by SBP Medical Discovery Institute Institutional Animal Care and Use Committee. The following transgenic lines were used: *TgBAC(pdx1:eGFP)*^bns13^ (Helker et al., 2019) and *Tg(ptf1a:eGFP)*^jh1^ (Godinho et al., 2005).

### Morpholino injections in zebrafish

The following previously reported morpholinos were injected into the yolk at the 1-cell stage in a final volume of either 0.5 or 1 nl: 0.75 ng lama1-ATG (5’-TCATCCT CATCTCCATCATCGCTCA -3’) (Pollard et al., 2006); 3 ng crb2a-SP, (5’-ACGTTGCCAGTACCTGTGTATCCTG-3’); 3 ng crb2b-SP, (5’-TAAAGATGTCCTACCCAGCTTGAAC-3’) (Omori & Malicki, 2006); 6.75 ng standard control MO (5’-CCTCTTACCTCAGTTACAATTTATA -3’). All morpholinos were obtained from GeneTools, LLC.

### Chromatin Immunoprecipitation Sequencing (ChIP-seq)

ChIP-seq was performed using the ChIP-IT High-Sensitivity kit (Active Motif) according to the manufacturer’s instructions. Briefly, for each cell stage and condition analyzed, 5-10 × 10^6^ cells were harvested and fixed for 15 min in an 11.1% formaldehyde solution. Cells were lysed and homogenized using a Dounce homogenizer and the lysate was sonicated in a Bioruptor® Plus (Diagenode), on high for 3 × 5 min (30 sec on, 30 sec off). Between 10 and 30 µg of the resulting sheared chromatin was used for each immunoprecipitation. Equal quantities of sheared chromatin from each sample were used for immunoprecipitations carried out at the same time. 4 µg of antibody were used for each ChIP-seq assay. Chromatin was incubated with primary antibodies overnight at 4°C on a rotator followed by incubation with Protein G agarose beads for 3 hours at 4 °C on a rotator. Antibodies used were rabbit anti-H3K27ac (Active Motif 39133), rabbit anti-H3K4me1 (Abcam ab8895), goat anti-CTCF (Santa Cruz Biotechnology SC-15914X), goat anti-GATA4 (Santa Cruz SC-1237), rabbit anti-GATA6 (Santa Cruz SC-9055), goat anti-FOXA1 (Abcam Ab5089), goat-anti-FOXA2 (Santa Cruz SC-6554), rabbit anti-PDX1 (BCBC AB1068), rabbit anti-HNF6 (Santa Cruz SC-13050), and rabbit anti-SOX9 (Chemicon AB5535). Reversal of crosslinks and DNA purification were performed according to the ChIP-IT High-Sensitivity instructions, with the modification of incubation at 65 °C for 2-3 hours, rather than at 80 °C for 2 hours. Sequencing libraries were constructed using KAPA DNA Library Preparation Kits for Illumina® (Kapa Biosystems) and library sequencing was performed on either a HiSeq 4000 System (Illumina®) or NovaSeq 6000 System (Illumina®) with single-end reads of either 50 or 75 base pairs (bp). Sequencing was performed by the Institute for Genomic Medicine (IGM) core research facility at the University of California at San Diego (UCSD). Two replicates from independent hESC differentiations were generated for each ChIP-seq experiment.

### ChIP-seq data analysis

ChIP-seq reads were mapped to the human genome consensus build (hg19/GRCh37) and visualized using the UCSC Genome Browser (Kent et al., 2002). Burrows-Wheeler Aligner (BWA) (Li & Durbin, 2009) version 0.7.13 was used to map data to the genome. Unmapped and low-quality (q<15) reads were discarded. SAMtools (Li et al., 2009) was used to remove duplicate sequences and HOMER (Heinz et al., 2010) was used to call peaks using default parameters and to generate tag density plots. Stage- and condition-matched input DNA controls were used as background when calling peaks. The BEDtools suite of programs (Quinlan & Hall, 2010) was used to perform genomic algebra operations. For all ChIP-seq experiments, replicates from two independent hESC differentiations were generated. Tag directories were created for each replicate using HOMER. Directories from each replicate were then combined, and peaks were called from the combined replicates. For histone modifications and CTCF peaks, pearson correlations between each pair of replicates were calculated over the called peaks using the command multiBamSummary from the deepTools2 package (Ramirez et al., 2016). For pancreatic lineage-determining transcription factors (GATA4, GATA6, FOXA1, FOXA2, HNF6, PDX1, SOX9), correlations were calculated for peaks overlapping PSSE. Calculated Pearson correlations are as follow:

|  | H3K4me1 | H3K27ac | CTCF |
| --- | --- | --- | --- |
| ES | 0.90 | 0.91 | 0.87 |
| DE | 0.97 | 0.84 | 0.86 |
| GT | 0.97 | 0.87 | 0.89 |
| PP1 | 0.97 | 0.85 | 0.89 |
| PP2 | 0.98 | 0.87 | 0.87 |

|  | GATA4 | GATA6 | FOXA1 | FOXA2 | HNF6 | PDX1 | SOX9 |
| --- | --- | --- | --- | --- | --- | --- | --- |
| PP2 | 0.86 | 0.82 | 0.87 | 0.80 | 0.95 | 0.64 | 0.86 |

### RNA isolation and sequencing (RNA-seq) and qRT-PCR

RNA was isolated from cell samples using the RNeasy® Micro Kit (Qiagen) according to the manufacturer instructions. For each cell stage and condition analyzed between 0.1 and 1 × 10^6^ cells were collected for RNA extraction. For qRT-PCR, cDNA synthesis was first performed using the iScript™ cDNA Synthesis Kit (Bio-Rad) and 500 ng of isolated RNA per reaction. qRT-PCR reactions were performed in triplicate with 10 ng of template cDNA per reaction using a CFX96™ Real-Time PCR Detection System and the iQ™ SYBR® Green Supermix (Bio-Rad). PCR of the TATA binding protein (TBP) coding sequence was used as an internal control and relative expression was quantified via double delta CT analysis. For RNA-seq, stranded, single-end sequencing libraries were constructed from isolated RNA using the TruSeq® Stranded mRNA Library Prep Kit (Illumina®) and library sequencing was performed on either a HiSeq 4000 System (Illumina®) or NovaSeq 6000 System (Illumina®) with single-end reads of either 50 or 75 base pairs (bp). Sequencing was performed by the Institute for Genomic Medicine (IGM) core research facility at the University of California at San Diego. A complete list of RT-qPCR primer sequences can be found below.

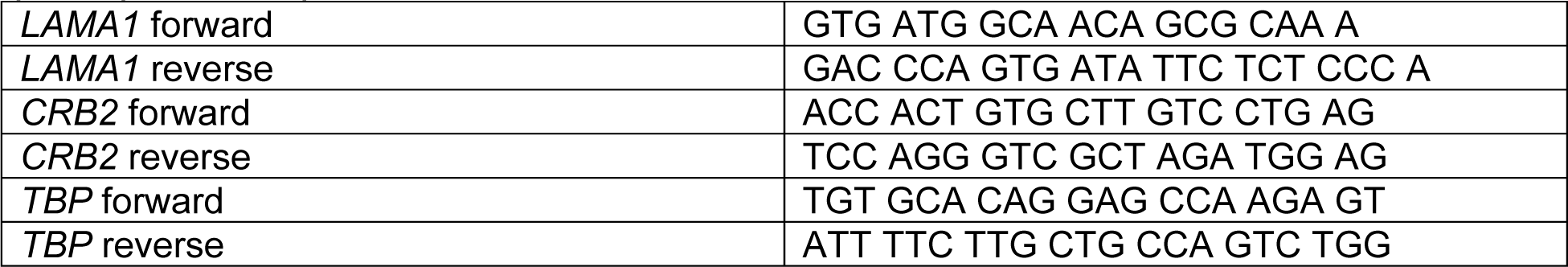

### RNA-seq data analysis

Reads were mapped to the human genome consensus build (hg19/GRCh37) using the Spliced Transcripts Alignment to a Reference (STAR) aligner v2.4 (Dobin et al., 2013). Normalized gene expression (fragments per kilobase per million mapped reads; FPKM) for each sequence file was determined using Cufflinks v2.2.1 (Trapnell et al., 2010) with the parameters: --library-type fr-firststrand --max-bundle-frags 10000000. For all RNA-Seq experiments, replicates from two independent hESC differentiations were generated. Pearson correlations between bam files corresponding to each pair of replicates were calculated, and are as follow:

|  |  |
| --- | --- |
| Δ*LAMA1*Enh clone 1 PP2 | 1.00 |
| Δ*LAMA1*Enh clone 2 PP2 | 0.99 |
| Δ*CRB2*Enh clone 1 PP2 | 0.98 |
| Δ*CRB2*Enh clone 2 PP2 | 0.90 |
| Δ*LAMA1*Enh control PP2 | 0.92 |
| Δ*CRB2*Enh control PP2 | 0.99 |

### Assay for Transposase Accessible Chromatin Sequencing (ATAC-seq)

ATAC-seq (Buenrostro et al., 2013) was performed on approximately 50,000 nuclei. The samples were permeabilized in cold permeabilization buffer (0.2% IGEPAL-CA630 (I8896, Sigma), 1 mM DTT (D9779, Sigma), Protease inhibitor (05056489001, Roche), 5% BSA (A7906, Sigma) in PBS (10010-23, Thermo Fisher Scientific) for 10 minutes on the rotator in the cold room and centrifuged for 5 min at 500 × g at 4°C. The pellet was resuspended in cold tagmentation buffer (33 mM Tris-acetate (pH = 7.8) (BP-152, Thermo Fisher Scientific), 66 mM K-acetate (P5708, Sigma), 11 mM Mg-acetate (M2545, Sigma), 16% DMF (DX1730, EMD Millipore) in Molecular biology water (46000-CM, Corning)) and incubated with tagmentation enzyme (FC-121-1030; Illumina) at 37 °C for 30 min with shaking at 500 rpm. The tagmented DNA was purified using MinElute PCR purification kit (28004, QIAGEN). Libraries were amplified using NEBNext High-Fidelity 2X PCR Master Mix (M0541, NEB) with primer extension at 72°C for 5 minutes, denaturation at 98°C for 30 s, followed by 8 cycles of denaturation at 98°C for 10 s, annealing at 63°C for 30 s and extension at 72°C for 60 s. After the purification of amplified libraries using MinElute PCR purification kit (28004, QIAGEN), double size selection was performed using SPRIselect bead (B23317, Beckman Coulter) with 0.55X beads and 1.5X to sample volume. Finally, libraries were sequenced on HiSeq4000 (Paired-end 50 cycles, Illumina).

### ATAC-seq data analysis

ATAC-seq reads were mapped to the human genome (hg19/GRCh37) using Burrows-Wheeler Aligner (BWA) version 0.7.13 (Li & Durbin, 2009), and visualized using the UCSC Genome Browser (Kent et al., 2002). SAMtools (Li et al., 2009) was used to remove unmapped, low-quality (q<15), and duplicate reads. MACS2 (Zhang et al., 2008) was used to call peaks, with parameters “shift set to 100 bps, smoothing window of 200 bps” and with “nolambda” and “nomodel” flags on. MACS2 was also used to call ATAC-Seq summits, using the same parameters combined with the “call-summits” flag.

For all ATAC-Seq experiments, replicates from two independent hESC differentiations were generated. Bam files for each pair of replicates were merged for downstream analysis using SAMtools, and Pearson correlations between bam files for each individual replicate were calculated over a set of peaks called from the merged bam file. Correlations were performed using the command multiBamSummary from the deepTools2 package (Ramirez et al., 2016) with the “—removeOutliers” flag and are as follow:

|  |  |
| --- | --- |
| ES | 0.95 |
| DE | 0.83 |
| GT | 1.00 |
| PP1 | 1.00 |
| PP2 | 1.00 |

### Hi-C data analysis

Hi-C data were processed as previously described with some modifications (Dixon et al., 2015). Read pairs were aligned to the hg19 reference genome separately using BWA-MEM with default parameters (Li & Durbin, 2009). Specifically, chimeric reads were processed to keep only the 5’ position and reads with low mapping quality (<10) were filtered out. Read pairs were then paired using custom scripts. Picard tools were then used to remove PCR duplicates. Bam files with alignments were further processed into text format as required by Juicebox tools (Durand et al., 2016). Juicebox tools were then applied to generate hic files containing normalized contact matrices. All downstream analysis was based on 10 Kb resolution KR normalized matrices.

Chromatin loops were identified by comparing each pixel with its local background, as described previously (Rao et al., 2014) with some modifications. Specifically, only the donut region around the pixel was compared to model the expected count. Briefly, the KR-normalized contact matrices at 10Kb resolution were used as input for loop calling. For each pixel, distance-corrected contact frequencies were calculated for each surrounding bin and the average of all surrounding bins. The expected counts were then transformed to raw counts by multiplying the counts with the raw-to-KR normalization factor. The probability of observing raw expected counts was calculated using Poisson distribution. All pixels with p-value < 0.01 and distance less than 10 Kb were selected as candidate pixels. Candidate pixels were then filtered to remove pixels without any neighboring candidate pixels since they were likely false positives. Finally, pixels within 20 Kb of each other were collapsed and only the most significant pixel was selected. The collapsed pixels with p-value < 1e-5 were used as the final list of chromatin loops.

### Definition of chromatin states

We collected or generated H3K4me1, H3K27ac, H3K4me1, H3K27me3 and CTCF ChIP-seq data at each developmental stage and in mature islets. Sequence reads were mapped to the human genome hg19 using bwa (version 0.7.12) (Li & Durbin, 2009), and low quality and duplicate reads were filtered using samtools (version 1.3) (Li et al., 2009). Using these reads, we then called chromatin states jointly across all data using chromHMM (version 1.12) (Ernst & Kellis, 2012) and used a 10-state model and 200bp bin size, as models with larger state numbers did not empirically resolve any additional informative states. We then assigned state names based on patterns defined by the NIH Epigenome Roadmap (Roadmap Epigenomics et al., 2015), which included active promoter/TssA (high H3K4me3, high H3K27ac), flanking TSS/TssFlnk1 (high H3K4me3), flanking TSS/TssFlnk 2 (high H3K4me3, high H3K27ac, high H3K4me1), bivalent Tss/TssBiv (high H3K27me3, high H3K4me3), poised enhancer/EnhP (high H3K4me1), insulator/CTCF (high CTCF), active enhancer/EnhA (high H3K27ac, high H3K4me1), repressor (high H3K27me3), and two quiescent (low signal for all assays) states. The state map with assigned names is shown in Figure 1 – figure supplement 1A.

We next defined stretch enhancer elements at each developmental stage and in mature islets. For each active enhancer (EnhA) element, we determined the number of consecutive 200bp bins covered by the enhancer. We then modeled the resulting bin counts for enhancers in each cell type using a Poisson distribution. Enhancers with a p-value less than .001 were labelled as stretch enhancers and otherwise labelled as traditional enhancers.

### Permutation-based significance

A random sampling approach (10,000 iterations) was used to obtain null distributions for enrichment analyses, in order to obtain p-values. Null distributions for enrichments were obtained by randomly shuffling enhancer regions using BEDTools (Quinlan & Hall, 2010) and overlapping with ATAC-seq peaks. P-values < 0.05 were considered significant.

### Assignment of enhancer target genes

Transcriptomes were filtered for genes expressed (FPKM ≥ 1) at each relevant stage, and BEDTools (Quinlan & Hall, 2010) was used to assign each enhancer to the nearest annotated TSS.

### Gene ontology

All gene ontology analyses were performed using Metascape (Zhou et al., 2019) with default parameters.

### Motif enrichment analysis

The findMotifsGenome.pl. command in HOMER (Heinz et al., 2010) was used to identify enriched transcription factor binding motifs. *de novo* motifs were assigned to transcription factors based on suggestions generated by HOMER.

### T2D-relevant trait enrichment analysis

GWAS summary statistics for T2D (Mahajan et al., 2018; Perry et al., 2012), metabolic traits (HOMA-B, HOMA-IR (Dupuis et al., 2010), fasting glucose, fasting insulin (Manning et al., 2012), fasting proinsulin (Strawbridge et al., 2011), 2 hour glucose adjusted for BMI (Saxena et al., 2010), HbA1c, insulin secretion rate, disposition index, acute insulin response, peak insulin response) (Wood et al., 2017), and developmental traits (head circumference (Taal et al., 2012), birth length (van der Valk et al., 2015), birth weight (Horikoshi et al., 2016)) conducted with individuals of European ancestry were obtained from various sources including the MAGIC consortium, EGG consortium, and authors of the studies. Custom LD score annotation files were created for PSSE, PP2 stretch enhancers, and islet stretch enhancers using LD score regression version 1.0.1 (Bulik-Sullivan et al., 2015). Enrichments for GWAS trait-associated variants within PSSE, PP2 stretch enhancers, and islet stretch enhancers were estimated with stratified LD score regression (Finucane et al., 2015). We next determined enrichment in the proportion of variants in accessible chromatin sites within islet SE and PSSE with nominal association to beta cell-related glycemic traits. For each trait, we calculated a 2×2 table of variants mapping in and outside of islet SE or PSSE and with or without nominal association and then determined significance using a chi-square test.

### Adipocyte differentiation analysis

Chromatin states for human adipose stromal cell (hASC) differentiation stages (1-4) were obtained from a published study (Varshney et al., 2017). PSSE were intersected with hASC chromatin states using BEDTools intersect (version 2.26.0) (Quinlan & Hall, 2010) with default parameters.

### Identification of T2D risk loci intersecting PSSE

T2D GWAS summary statistics were obtained from the DIAMANTE consortium (Mahajan et al., 2018). Intersection of variants and PSSE was performed using BEDTools intersect (version 2.26.0) (Quinlan & Hall, 2010) with default parameters. The adjusted significance threshold was set at P < 4.66 × 10^−6^ (Bonferroni correction for 10,738 variants mapping in PSSE). Putative novel loci were defined as those with 1) at least one variant in a PSSE reaching the adjusted significance threshold and 2) mapping at least 500 kb away from a known T2D locus.

### ATAC-seq footprinting analysis

ATAC-seq footprinting was performed as previously described (Aylward et al., 2018). In brief, diploid genomes for CyT49 were created using vcf2diploid (version 0.2.6a) (Rozowsky et al., 2011) and genotypes called from whole genome sequencing and scanned for a compiled database of TF sequence motifs from JASPAR (Mathelier et al., 2016) and ENCODE (Consortium, 2012) with FIMO (version 4.12.0) (Grant et al., 2011) using default parameters for p-value threshold and a 40.9% GC content based on the hg19 human reference genome. Footprints within ATAC-seq peaks were discovered with CENTIPEDE (version 1.2) (Pique-Regi et al., 2011) using cut-site matrices containing Tn5 integration counts within a ±100 bp window around each motif occurrence. Footprints were defined as those with a posterior probability ≥ 0.99.

### Generation of similarity matrices for total transcriptomes

For each replicate, FPKM values corresponding to total transcriptome were filtered for genes expressed (FPKM ≥ 1) in ≥ 1 replicate. For expressed genes, log(FPKM+1) values were used to calculate Pearson correlations.

### Immunofluorescence analysis

Cell aggregates derived from hESCs were allowed to settle in microcentrifuge tubes and washed twice with PBS before fixation with 4% paraformaldehyde (PFA) for 30 min at room temperature. Fixed samples were washed twice with PBS and incubated overnight at 4 °C in 30% (w/v) sucrose in PBS. Samples were then loaded into disposable embedding molds (VWR), covered in Tissue-Tek® O.C.T. Sakura® Finetek compound (VWR) and flash frozen on dry ice to prepare frozen blocks. The blocks were sectioned at 10 µm and sections were placed on Superfrost Plus® (Thermo Fisher) microscope slides and washed with PBS for 10 min. Slide-mounted cell sections were permeabilized and blocked with blocking buffer, consisting of 0.15% (v/v) Triton X-100 (Sigma) and 1% (v/v) normal donkey serum (Jackson Immuno Research Laboratories) in PBS, for 1 hour at room temperature. Slides were then incubated overnight at 4 °C with primary antibody solutions. The following day slides were washed five times with PBS and incubated for 1 hour at room temperature with secondary antibody solutions. Cells were washed five times with PBS before coverslips were applied.

All antibodies were diluted in blocking buffer at the ratios indicated below. Primary antibodies used were goat anti-PDX1 (1:500 dilution, Abcam ab47383); and mouse anti-NKX6.1 (1:300 dilution, Developmental Studies Hybridoma Bank F64A6B4). Secondary antibodies against goat and mouse were Alexa488- and Cy3-conjugated donkey antibodies, respectively (Jackson Immuno Research Laboratories 705-545-003 and 715-165-150, respectively), and were used at dilutions of 1:500 (anti-goat Alexa488) or 1:1000 (anti-mouse Cy3). Cell nuclei were stained with Hoechst 33342 (1:3000, Invitrogen). Representative images were obtained with a Zeiss Axio-Observer-Z1 microscope equipped with a Zeiss ApoTome and AxioCam digital camera. Figures were prepared in Adobe Creative Suite 5.

### Flow cytometry analysis

Cell aggregates derived from hESCs were allowed to settle in microcentrifuge tubes and washed with PBS. Cell aggregates were incubated with Accutase® at room temperature until a single-cell suspension was obtained. Cells were washed with 1 mL ice-cold flow buffer comprised of 0.2% BSA in PBS and centrifuged at 200 g for 5 min. BD Cytofix/Cytoperm™ Plus Fixation/Permeabilization Solution Kit was used to fix and stain cells for flow cytometry according to the manufacturer’s instructions. Briefly, cell pellets were re-suspended in ice-cold BD Fixation/Permeabilization solution (300 µL per microcentrifuge tube). Cells were incubated for 20 min at 4 °C. Cells were washed twice with 1 mL ice-cold 1 × BD Perm/Wash™ Buffer and centrifuged at 10 °C and 200 × g for 5 min. Cells were re-suspended in 50 µL ice-cold 1 × BD Perm/Wash™ Buffer containing diluted antibodies, for each staining performed. Cells were incubated at 4 °C in the dark for 1-3 hrs. Cells were washed with 1.25 mL ice-cold 1X BD Wash Buffer and centrifuged at 200 × g for 5 min. Cell pellets were re-suspended in 300 µL ice-cold flow buffer and analyzed in a FACSCanto™ II (BD Biosciences). Antibodies used were PE-conjugated anti-PDX1 (1:10 dilution, BD Biosciences); and AlexaFluor® 647-conjugated anti-NKX6.1 (1:5 dilution, BD Biosciences). Data were processed using FlowJo software v10.

### Whole mount immunohistochemistry

Zebrafish larvae were fixed and stained according to published protocols (Lancman et al., 2013) using the following antibodies: chicken anti-GFP (1:200; Aves Labs; GFP-1020), guinea pig anti-insulin (1:200; Biomeda; v2024), mouse anti-Crb2a (1:100; ZIRC; zs-4), rabbit anti-panCrb (1:100; provided by Dr. Abbie M. Jensen at University of Massachusetts, Amherst (Hsu & Jensen, 2010)), rabbit anti-laminin (1:100; Sigma;L9393) and DAPI (1:200; 500 mg/ml, Invitrogen; D1306).

### Imaging and quantification of beta cell numbers in zebrafish

To quantify beta cell numbers, 50 and 78 hpf zebrafish larvae were stained for confocal imaging using DAPI and guinea pig anti-insulin antibody (1:200; Biomeda; v2024). Whole mount fluorescent confocal Z-stacks (0.9 μm steps) images were collected for the entire islet with optical slices captured at a focal depth of 1.8 μm. Samples were imaged using a Zeiss 710 confocal microscope running Zen 2010 (Black) software. Final images were generated using Adobe Photoshop CS6 and/or ImageJ64 (vs.1.48b).

### Data sources

The following datasets used in this study were downloaded from the GEO and ArrayExpress repositories:

RNA-seq: Pancreatic differentiation of CyT49 hESC line (E-MTAB-1086); primary islet data (GSE115327)
ChIP-seq: H3K27ac data in primary islets (E-MTAB-1919 and GSE51311); H3K27ac data in CyT49 hESC/DE/GT/PP1/PP2 (GSE54471); H3K4me1 data in CyT49 ES/DE/GT/PP1/PP2 (GSE54471); H3K27me3 and H3K4me3 in CyT49 ES/DE/GT/PP1/PP2 (E-MTAB-1086); PDX1 in CyT49 PP2 (GSE54471); samples from ROADMAP consortium: http://ncbi.nlm.nih.gov/geo/roadmap/epigenomics
ATAC-seq: primary islet data (GSE115327 and PRJN527099); CyT49 GT and PP2 (GSE115327).

Hi-C datasets were generated in collaboration with the Ren laboratory at University of California, San Diego as a component of the 4D Nucleome Project (Dekker et al., 2017) under accession number 4DNES0LVRKBM.

## QUANTIFICATION AND STATISTICAL ANALYSES

Statistical analyses were performed using GraphPad Prism (v8.1.2), and R (v3.6.1). Statistical parameters, such as the value of n, mean, standard deviation (SD), standard error of the mean (SEM), significance level (*p < 0.05, **p < 0.01, and ***p < 0.001), and the statistical tests used, are reported in the figures and figure legends. The ‘‘n’’ refers to the number of independent pancreatic differentiation experiments analyzed (biological replicates).

Statistically significant gene expression changes were determined with DESeq2 (Love et al., 2014).

## DATA AVAILABILITY

All mRNA-seq, ChIP-seq, and ATAC-seq datasets generated for this study have been deposited at GEO under the accession number GSE149148.

## Supporting information

Essential Reagents Table

## ACKNOWLEDGEMENTS

We thank Ileana Matta for assistance with ATAC-seq assays and library preparations, as well as the Sander and Gaulton laboratories for helpful discussions. We also thank Dr. Abbie Jensen at University of Massachusetts, Amherst for the anti-panCrb antibody. We acknowledge support of the UCSD Human Embryonic Stem Cell Core for cell sorting and the K. Jepsen and UCSD IGM Genomic Center for library preparation and sequencing. This work was supported by grant T32 GM008666 (R.J.G.), grant P30 DK064391 (K.J.), R01 DK068471 and U01 DK105541 (M.S., B.R.).

## AUTHOR CONTRIBUTIONS

A.W., K.J.G., and M.S. conceived the project. A.W., R.J.G, J.C., J.J.L., P.D.S.D., K.J.G., and M.S. designed experiments. A.W., R.J.G., J.J.L., N.W., S.K., J.W., and J.Y. performed experiments. A.W., R.J.G, J.C., K.J.G., Y.Q., and A.A. analyzed sequencing data. J.C. and K.J.G. performed GWAS analyses. A.W., R.J.G, J.C., J.J.L., P.D.S.D., K.J.G., and M.S. interpreted data. A.W., R.J.G, J.C., J.J.L., P.D.S.D., K.J.G., and M.S. wrote the manuscript. P.D.S.D., B.R., K.J.G., and M.S. supervised all research.

---

**Figure 1 – figure supplement 1:**
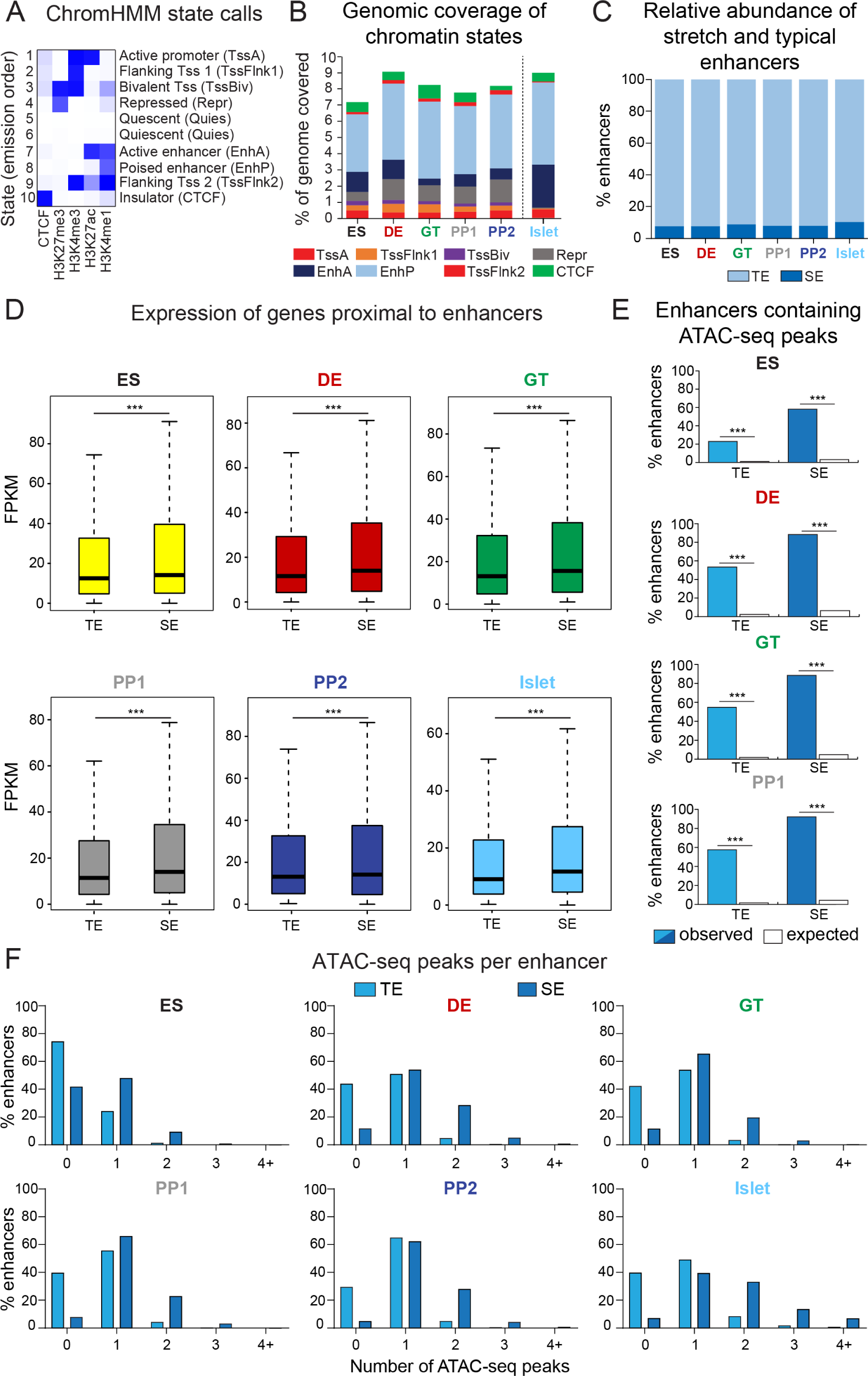
Characterization of typical and stretch enhancers in pancreatic developmental intermediates and islets. (**A**) Diagram illustrating incorporation of histone modification and CTCF ChIP-seq data to generate chromatin state calls via ChromHMM. (**B**) Percentage of the genome covered by defined chromatin states at each developmental stage and in primary human islets. Chromatin states are based on ChromHMM classifications: TssA, active promoter; TssFlnk, flanking transcription start site; TssBiv, bivalent promoter; Repr, repressed; EnhA, active enhancer; EnhP, poised enhancer. (**C**) Percentage of typical (TE) and stretch enhancers (SE) relative to all enhancers at each developmental stage and in islets. (**D**) Box plots showing mRNA levels based on RNA-seq of nearest expressed genes (fragments per kilobase per million fragments mapped (FPKM) ≥ 1) for TE and SE at each developmental stage and in islets (*** p = 4.68 × 10^−7^, 4.64 × 10^−11^, 1.31 × 10^−5^, 8.85 × 10^−9^, 5.34 × 10^−6^, and < 2.2 × 10^−16^ for TE vs SE comparisons in ES, DE, GT, PP1, PP2, and islets, respectively; Wilcoxon rank sum test, 2 sided). Plots are centered on median, with box encompassing 25th-75th percentile and whiskers extending up to 1.5 interquartile range. n = 3 replicates from independent differentiations at ES, DE, GT, PP1, and PP2, respectively; n = 2 islet replicates. (**E**) Percentage of TE and SE overlapping with at least one ATAC-seq peak in ES, DE, GT, or PP1. Enrichment analysis comparing observed and expected overlap based on random genomic regions of the same size and located on the same chromosome averaged over 10,000 iterations (*** p < 1 × 10^−4^; permutation test). (**F**) Percentage of TE and SE overlapping 0, 1, 2, 3, or 4+ ATAC-seq peaks at each developmental stage and in islets.

**Figure 2 – figure supplement 1:**
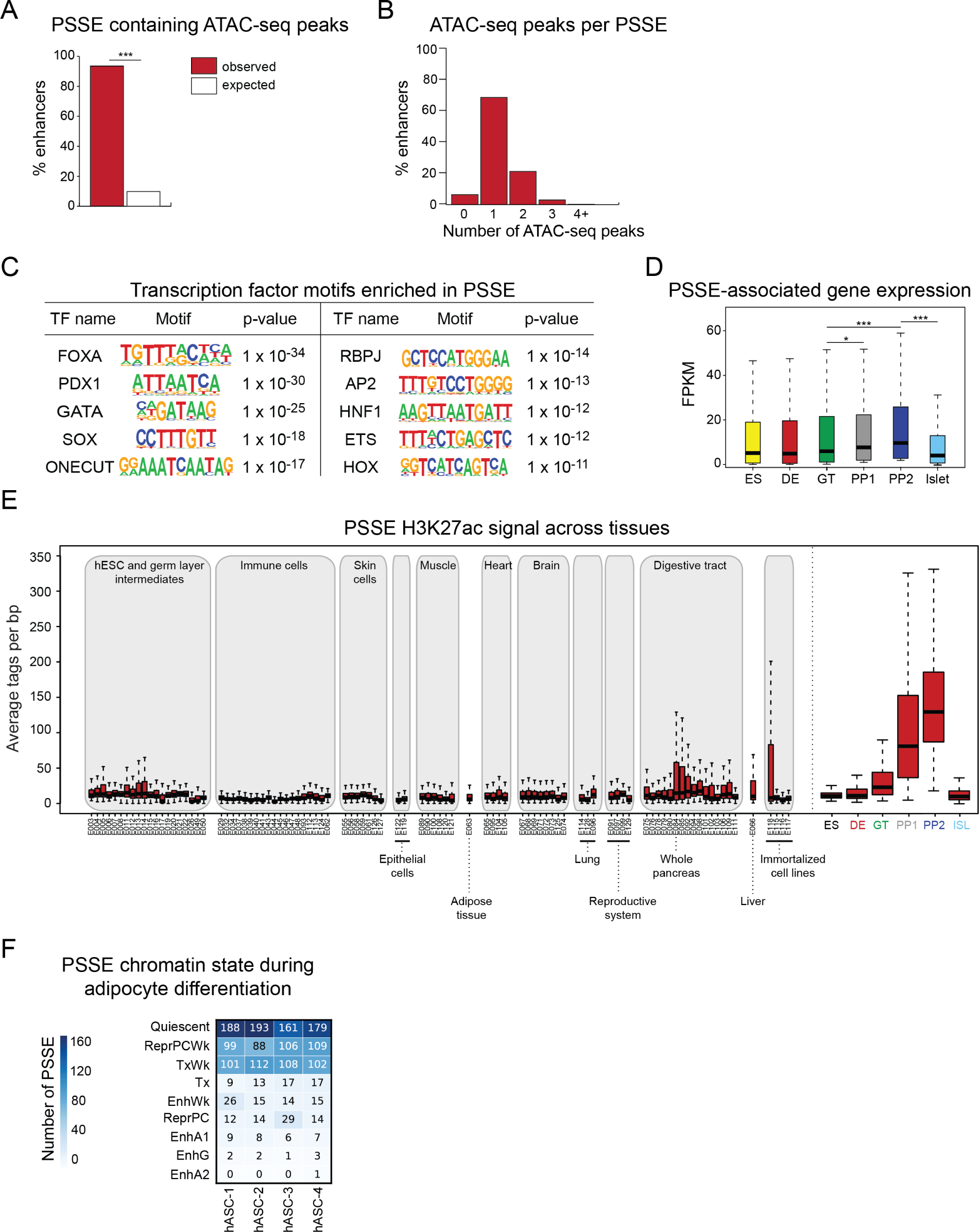
Characterization of pancreatic progenitor-specific stretch enhancers. (**A**) Percentage of PSSE overlapping with at least one ATAC-seq peak at PP2. Enrichment analysis comparing observed and expected overlap based on random genomic regions of the same size and located on the same chromosome averaged over 10,000 iterations (*** p < 1 × 10^−4^; permutation test). (B) Percentage of PSSE overlapping 0, 1, 2, 3, or 4+ ATAC-seq peaks at PP2. (**C**) Enriched transcription factor (TF) binding motifs with associated p-values at ATAC-seq peaks at PP2 intersecting PSSE. Fisher’s exact test, 2 sided, corrected for multiple comparisons. (**D**) Box plots showing mRNA levels based on RNA-seq of nearest expressed genes (fragments per kilobase per million fragments mapped (FPKM) ≥ 1 at PP2) for PSSE at each developmental stage and in islets (* p = 1.10 × 10^−2^ for GT vs PP1; *** p = 1.80 × 10^−8^ for GT vs PP2, p < 2.2 × 10^−16^ for PP2 vs islet; Wilcoxon rank sum test, 2 sided). Plots are centered on median, with box encompassing 25th-75th percentile and whiskers extending up to 1.5 interquartile range. n = 3 replicates from independent differentiations at ES, DE, GT, PP1, and PP2, respectively; n = 2 islet replicates. (**E**) Box plots showing H3K27ac signal at PSSE in tissues and cell lines from the ENCODE and Epigenome Roadmap projects as well as in developmental intermediates and islets (ISL). Plots are centered on median, with box encompassing 25th-75th percentile and whiskers extending up to 1.5 interquartile range. See also Figure 2 – source data 4. (**F**) Number of PSSE overlapping defined chromatin states in human adipose stromal cells from preadipose (hASC1) to mature adipose state (hASC4) (from Varshney, et al 2017). ChromHMM classifications: Quiescent; ReprPCWk, Weak Repressed PolyComb; TxWk, Weak Transcription; Tx, Strong Transcription; EnhWk, Weak Enhancer; ReprPC, Repressed Polycomb; EnhA1, Active Enhancer 1; EnhG, Genic Enhancer; EnhA2, Active Enhancer 2.

**Figure 3 – figure supplement 1:**
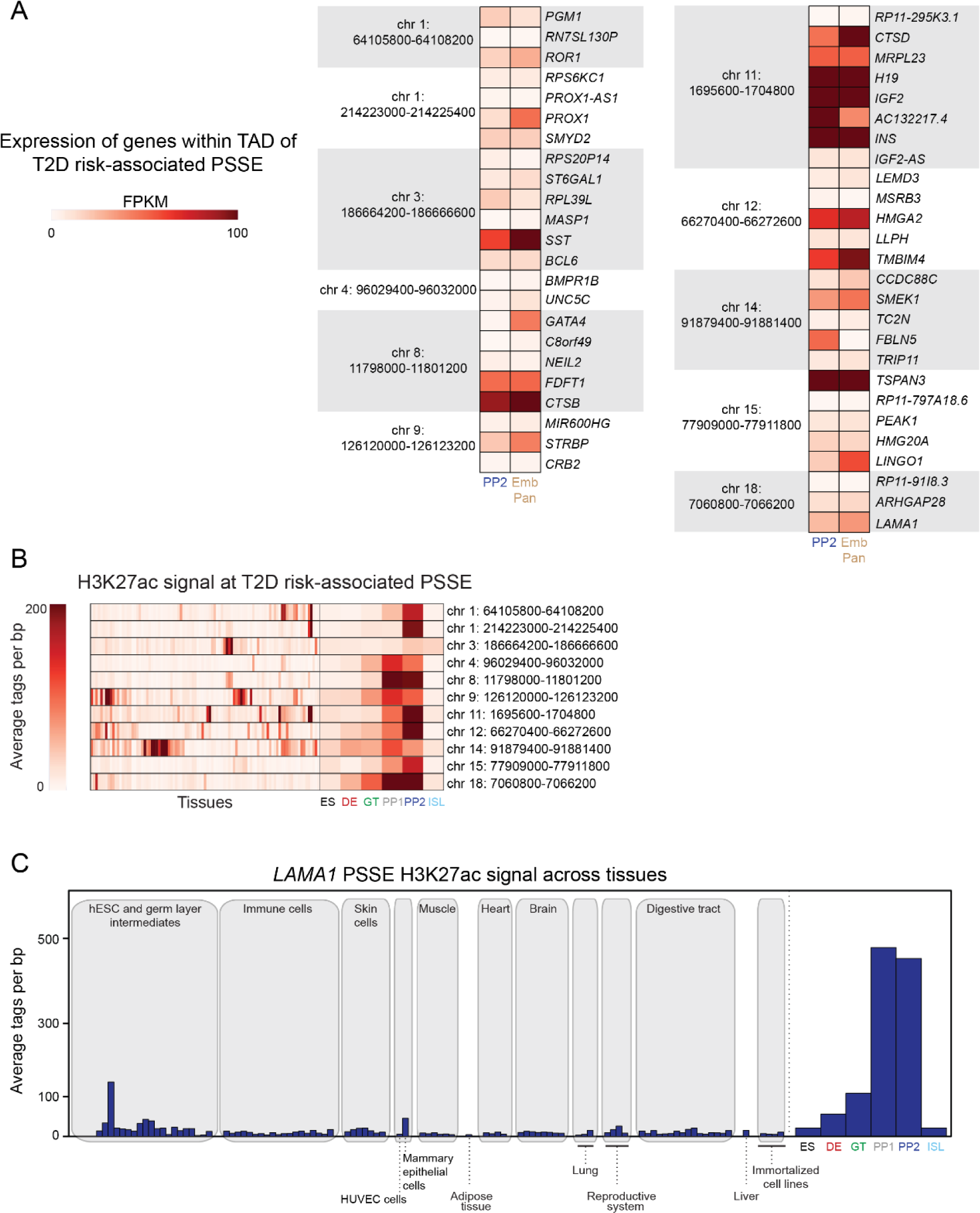
Activity of T2D risk-associated pancreatic progenitor-specific stretch enhancers across human tissues. (**A**) mRNA levels (measured in fragments per kilobase per million fragments mapped (FPKM)) at PP2 and in human embryonic pancreas (54 and 58 days gestation, Emb Pan) of all genes expressed (FPKM ≥ 1) at PP2 and located within topologically associated domains (TADs) containing indicated PSSE harboring T2D variants identified in Fig. 3A. (**B**) Heatmap showing H3K27ac signal at PSSE harboring T2D variants identified in Fig. 3A. Quantification in tissues and cell lines from the ENCODE and Epigenome Roadmap projects (tissues) as well as in developmental intermediates and islets (ISL) is shown. (**C**) H3K27ac signal at *LAMA1*-associated PSSE in tissues and cell lines from the ENCODE and Epigenome Roadmap projects as well as in developmental intermediates and islets.

**Figure 4 – figure supplement 1:**
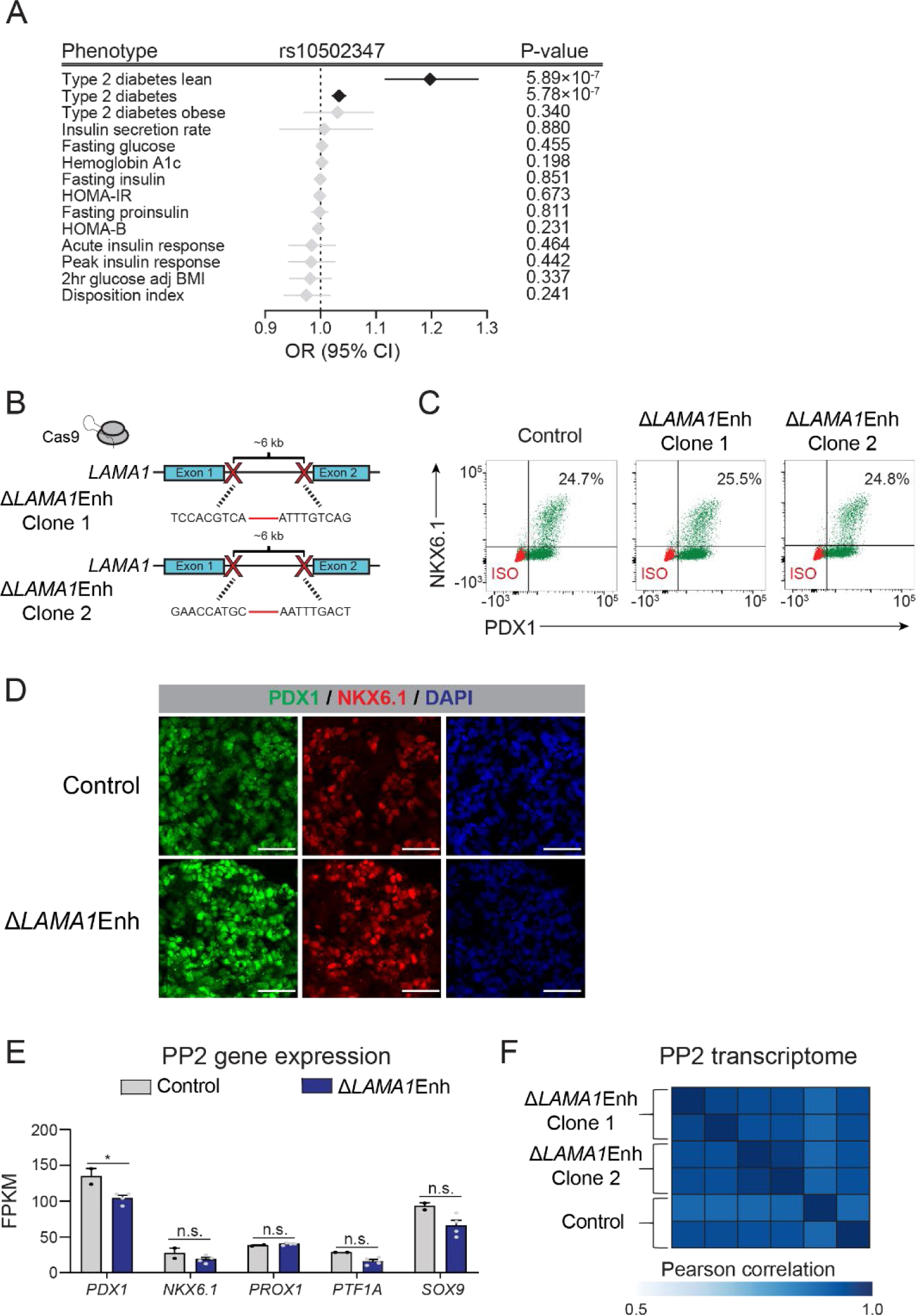
Deletion of the *LAMA1*-associated pancreatic progenitor-specific enhancer does not affect pancreatic lineage specification. (**A**) Odds ratio estimates (points) and 95% CIs (lines) for rs10502347 association with T2D and metabolic GWAS traits. Significant associations are colored black, non-significant are colored light grey. (**B**) Schematic illustrating CRISPR-Cas9-mediated deletion strategy of *LAMA1*-associated PSSE to generate independent Δ*LAMA1*Enh hESC clones with different DNA cleavage products. (**C**) Flow cytometry analysis for NKX6.1 and PDX1 comparing control and Δ*LAMA1*Enh PP2 cells. Isotype control (ISO) for each antibody is shown in red and target protein staining in green. Percentage of cells expressing each protein is indicated (representative experiment, n = 3 independent differentiations). (**D**) Immunofluorescent staining for NKX6.1 and PDX1 in control and Δ*LAMA1*Enh PP2 cells (representative images, n = 2 slides). Scale bar, 50 μm. (**E**) mRNA expression of pancreatic transcription factors determined by RNA-seq in control and Δ*LAMA1*Enh PP2 cells. Data are shown as mean of fragments per kilobase per million fragments mapped (FPKM) ± S.E.M. (n = 2 replicates from independent differentiations for control cells. Δ*LAMA1*Enh cells represent combined data from 2 clonal lines with 2 replicates for each line from independent differentiations. p adj. = 3.56 × 10^−2^, 0.224, 0.829, 8.14 × 10^−2^, and 0.142, for comparisons of *PDX1*, *NKX6.1*, *PROX1*, *PTF1A*, and *SOX9*, respectively; DESeq2; * p adj. < 0.05, n.s., not significant). (**F**) Similarity matrix showing Pearson correlations for normalized transcriptomes (log transformed expression for genes with FPKM ≥ 1 in ≥ 1 replicates) in control and Δ*LAMA1*Enh PP2 cells (n = 2 independent differentiations for control cells and for each Δ*LAMA1*Enh clone). See also Figure 4 – source data 1 and 2.

**Figure 5 – figure supplement 1:**
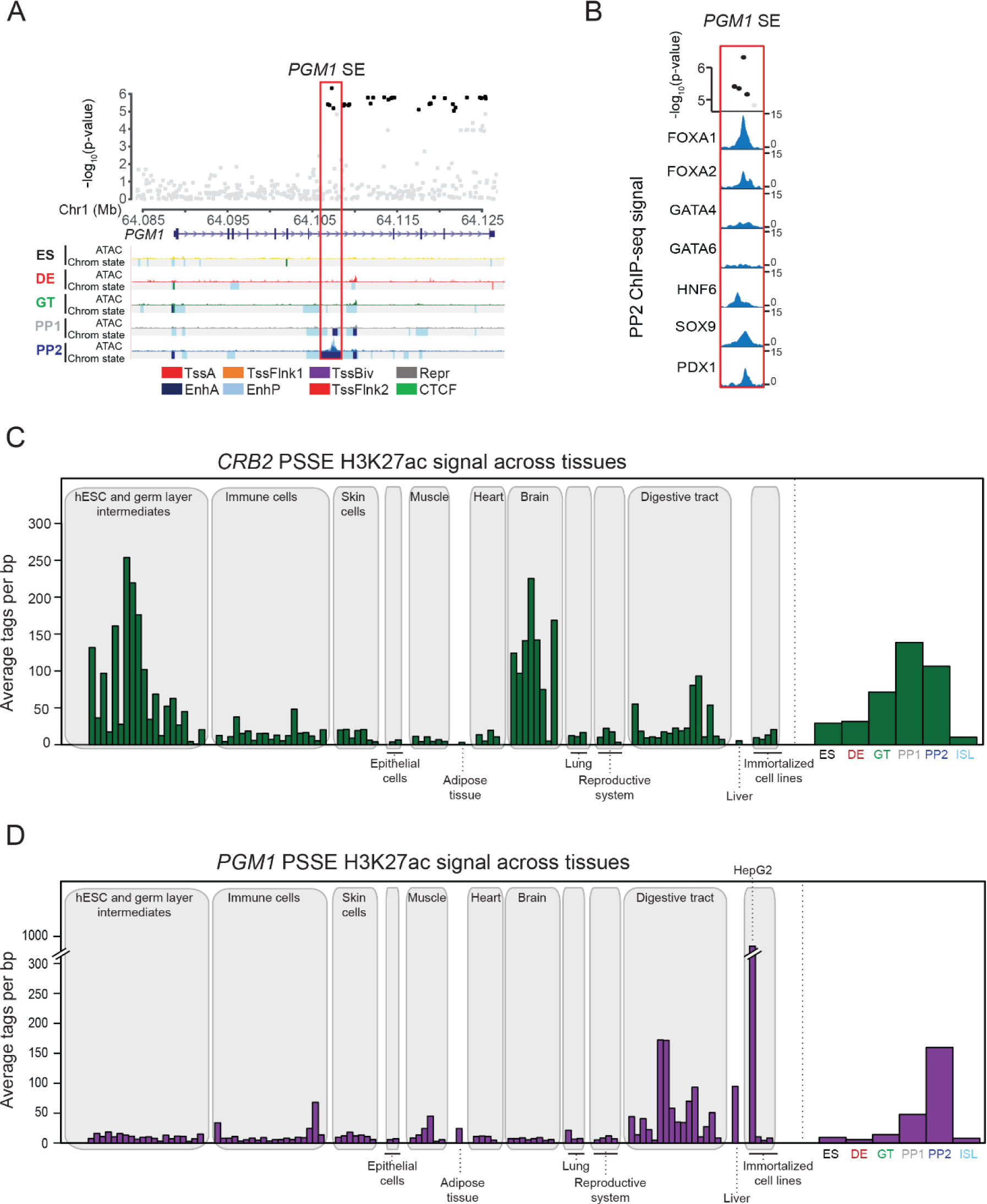
Activity of *CRB2-* and *PGM1-*associated pancreatic progenitor-specific stretch enhancers across human tissues. (**A**) (Top) Locus plots showing T2D association p-values for variants in a 43 kb window (hg19 chr1:64,084,000-64,127,000) at the *PGM1* locus and *PGM1* PSSE (red box). Fine mapped variants within the 99% credible set for the novel *PGM1* locus are colored black. All other variants are colored light gray. (Bottom) Chromatin states and ATAC-seq signal in ES, DE, GT, PP1, and PP2. TssA, active promoter; TssFlnk, flanking transcription start site; TssBiv, bivalent promoter; Repr, repressed; EnhA, active enhancer; EnhP, poised enhancer. (**B**) FOXA1, FOXA2, GATA4, GATA6, HNF6, SOX9, and PDX1 ChIP-seq profiles at the *PGM1* PSSE in PP2 cells. The variants rs2269247, rs2301055, rs2301054, and rs2269246 (black) overlap with transcription factor binding sites. (B) H3K27ac signal at *CRB2-*associated PSSE in tissues and cell lines from the ENCODE and Epigenome Roadmap projects as well as in developmental intermediates and islets (ISL). (**D**) H3K27ac signal at *PGM1-*associated PSSE in tissues and cell lines from the ENCODE and Epigenome Roadmap projects as well as in developmental intermediates and islets.

**Figure 5 – figure supplement 2:**
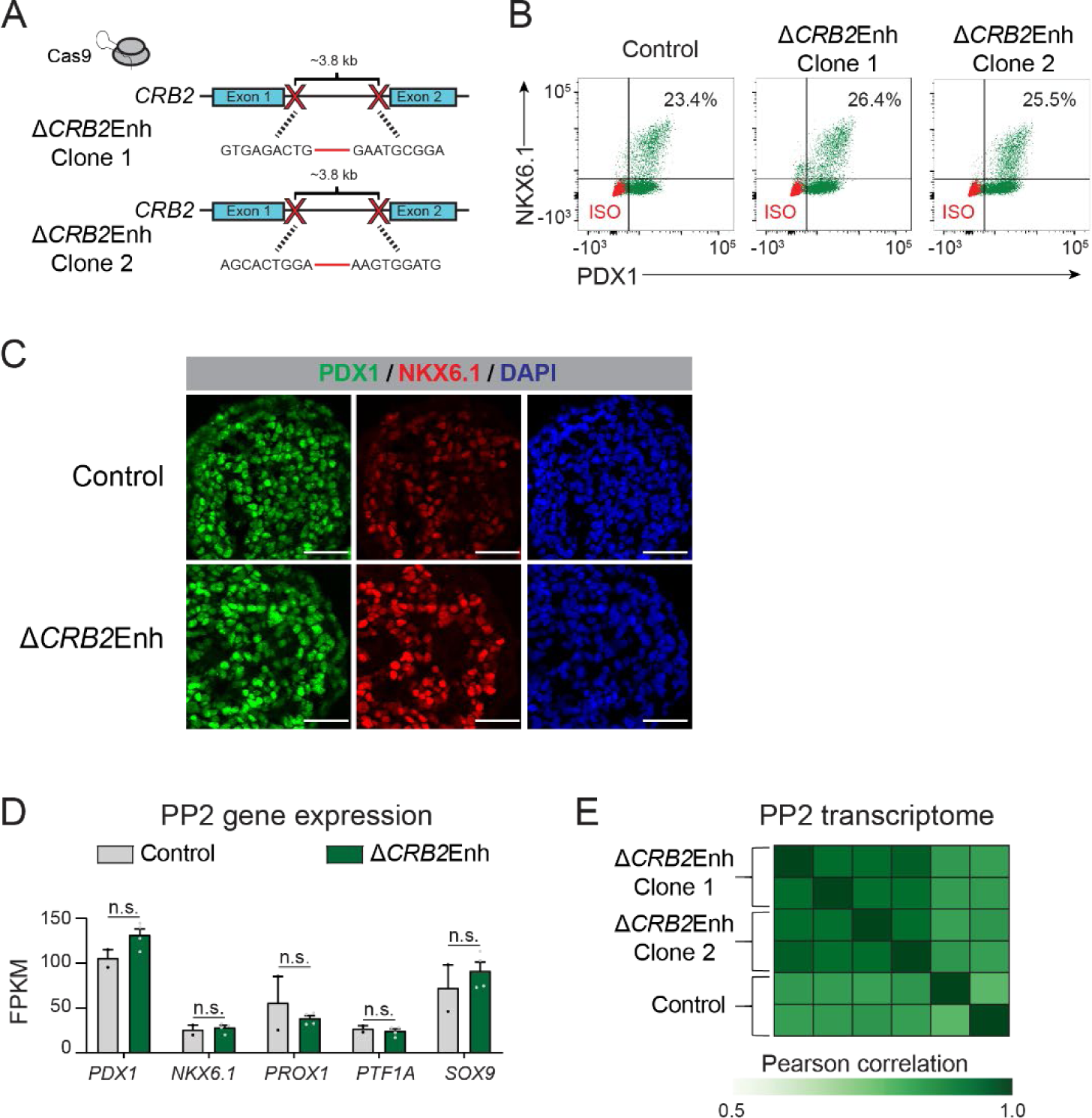
Deletion of the *CRB2*-associated pancreatic progenitor-specific enhancer does not affect pancreatic lineage specification. (**A**) Schematic illustrating CRISPR-Cas9-mediated deletion strategy of *CRB2*-associated PSSE to generate independent Δ*CRB2*Enh hESC clones with different DNA cleavage products. (**B**) Flow cytometry analysis for NKX6.1 and PDX1 comparing control and Δ*CRB2*Enh PP2 cells. Isotype control (ISO) for each antibody is shown in red and target protein staining in green. Percentage of cells expressing each protein is indicated (representative experiment, n = 3 independent differentiations). (**C**) Immunofluorescent staining for NKX6.1 and PDX1 in control and Δ*CRB2*Enh PP2 cells (representative images, n = 2 independent slides). Scale bar, 50 μm. (**D**) mRNA expression of pancreatic transcription factors determined by RNA-seq in control and Δ*CRB2*Enh PP2 cells. Data are shown as mean of fragments per kilobase per million fragments mapped (FPKM) ± S.E.M. (n = 2 replicates from independent differentiations for control cells Δ*CRB2*Enh cells represent combined data from 2 clonal lines with 2 replicates for each line from independent differentiations. p adj. = 1.00, 1.00, 1.00, 1.00, and 1.00, for comparisons of *PDX1*, *NKX6.1*, *PROX1*, *PTF1A*, and *SOX9*, respectively; DESeq2; n.s., not significant). (**E**) Similarity matrix showing Pearson correlations for normalized transcriptomes (log transformed expression for genes with FPKM ≥ 1 in ≥ 1 replicates) in control and Δ*CRB2*Enh PP2 cells (n = 2 independent differentiations for control cells and for each Δ*CRB2*Enh clone). See also Figure 5 – source data 1.

**Figure 6 – figure supplement 1:**
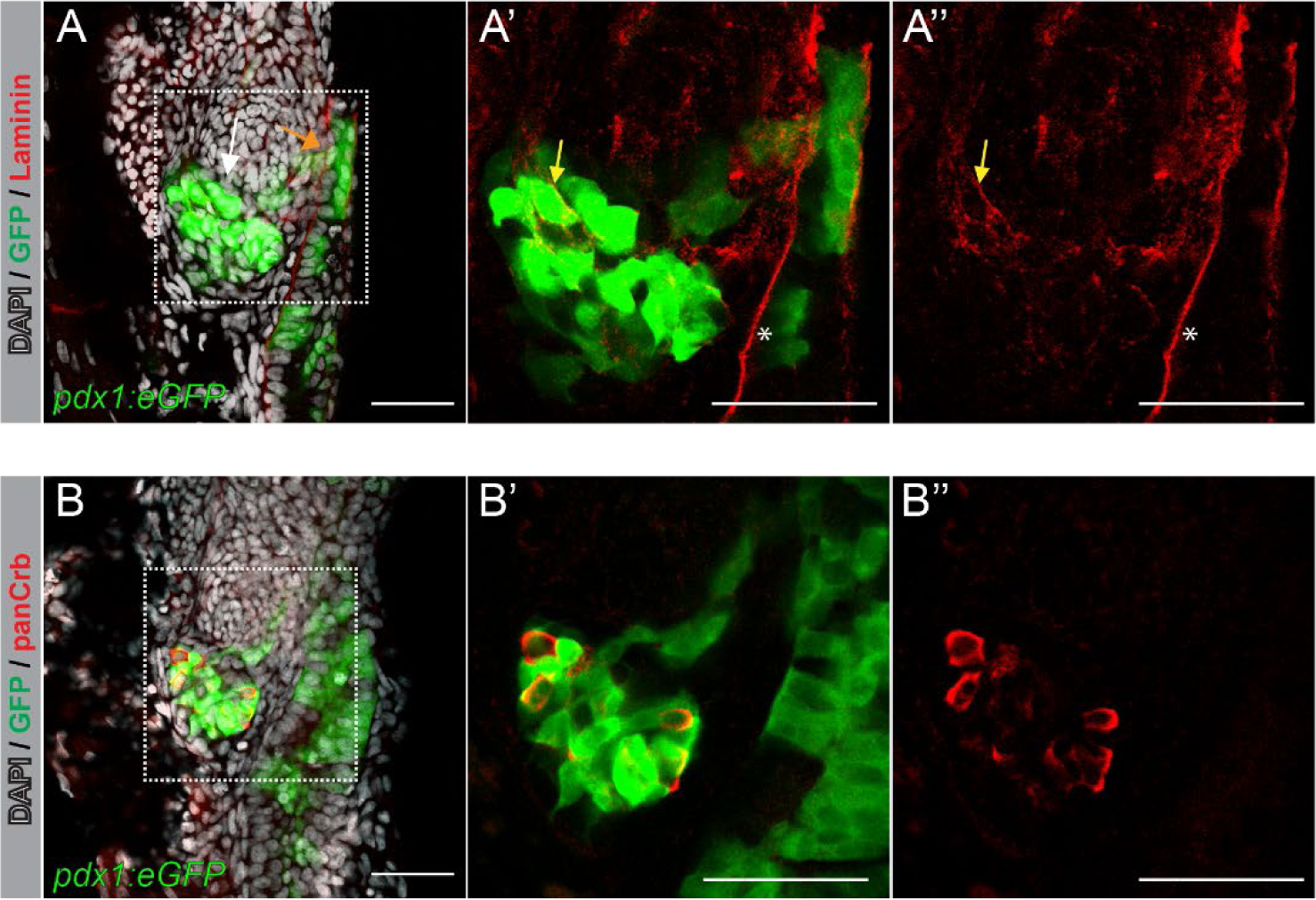
Laminin and Crb are expressed in the embryonic zebrafish pancreas. (**A,B**) 3-dimensional renderings of 48 hours post fertilization (hpf) *TgBAC(pdx1:eGFP)* zebrafish foregut endoderm stained with DAPI (nuclei, grey) and antibodies against laminin (**A**, red) or panCrb (**B**, red); n = 10 embryos stained. (**A-A”**) Representative Z-focal plane image showing pancreatic cells marked by *pdx1:eGFP* (green) expressing laminin (yellow arrow). Laminin is broadly expressed throughout the intestinal lining (star). White and orange arrows indicate developing pancreas and duodenum, respectively. (**B-B”**) Representative Z-focal plane image showing panCrb staining in pancreatic cells expressing *pdx1:eGFP*. Scale bar, 40 µM.

**Figure 6 – figure supplement 2:**
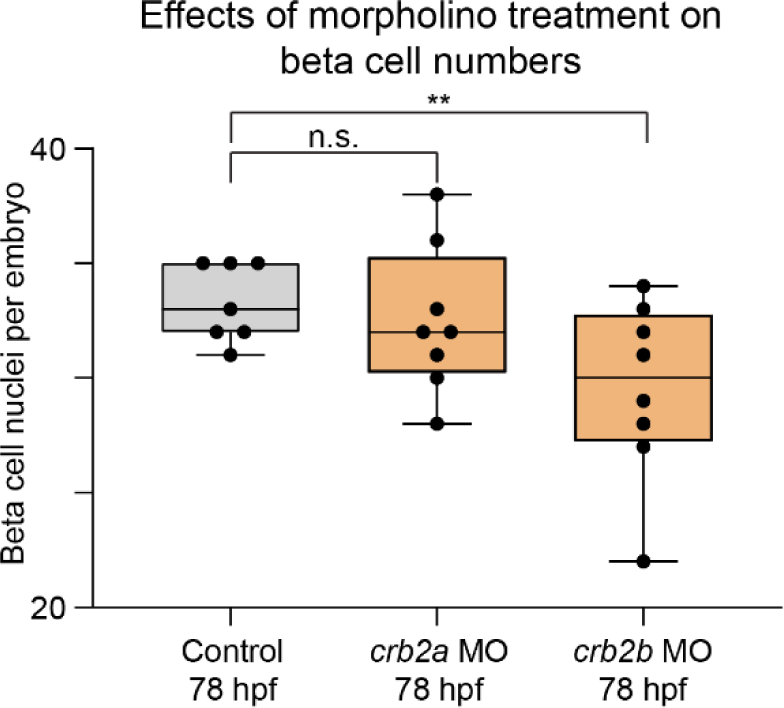
*crb2b* but not *crb2a* regulates pancreatic beta cell differentiation. Quantification of beta (insulin^+^) cell nuclei per embryo in *Tg(ptf1a:eGFP)* control zebrafish embryos and *crb2a* or *crb2b* morphants at 78 hours post fertilization (hpf). p adj. = 0.91 and 4.4 × 10^−2^ for comparison of control (n = 7 embryos) to *crb2a* (n = 8) or *crb2b* (n = 8) morphants, respectively; ANOVA-Dunnett’s multiple comparison test; ** p < 0.01, n.s., not significant. 0 out of 8 *crb2a*, 0 out of 8 *crb2b* morphants displayed an annular pancreas.

## SOURCE DATA

**Figure 2 – source data 1:** Chromosomal coordinates of pancreatic progenitor-specific stretch enhancers (PSSE).

**Figure 2 – source data 2:** Enriched gene ontology terms for PSSE-associated genes.

**Figure 2 – source data 3:** Proportion of variants nominally associated with beta cell functional traits.

**Figure 2 – source data 4:** Tissue identity of downloaded data from ROADMAP consortium.

**Figure 4 – source data 1:** Genes downregulated in Δ*LAMA1*Enh PP2 stage cells compared to control cells (p adj. < .05).

**Figure 4 – source data 2:** Genes upregulated in Δ*LAMA1*Enh PP2 stage cells compared to control cells (p adj. < .05).

**Figure 5 – source data 1:** Genes downregulated in Δ*CRB2*Enh PP2 stage cells compared to control cells (p adj. < .05).

## REFERENCES

Ahnfelt-Ronne, J., Ravassard, P., Pardanaud-Glavieux, C., Scharfmann, R., & Serup, P. (2010, Aug). Mesenchymal bone morphogenetic protein signaling is required for normal pancreas development. Diabetes, 59(8), 1948–1956. https://doi.org/10.2337/db09-1010

Alves, C. H., Bossers, K., Vos, R. M., Essing, A. H., Swagemakers, S., van der Spek, P. J., Verhaagen, J., & Wijnholds, J. (2013). Microarray and morphological analysis of early postnatal CRB2 mutant retinas on a pure C57BL/6J genetic background. PLoS One, 8(12), e82532. https://doi.org/10.1371/journal.pone.0082532

Aylward, A., Chiou, J., Okino, M. L., Kadakia, N., & Gaulton, K. J. (2018, Nov 7). Shared genetic risk contributes to type 1 and type 2 diabetes etiology. Hum Mol Genet. https://doi.org/10.1093/hmg/ddy314

Buenrostro, J. D., Giresi, P. G., Zaba, L. C., Chang, H. Y., & Greenleaf, W. J. (2013, Dec). Transposition of native chromatin for fast and sensitive epigenomic profiling of open chromatin, DNA-binding proteins and nucleosome position. Nat Methods, 10(12), 1213–1218. https://doi.org/10.1038/nmeth.2688

Bulgakova, N. A., & Knust, E. (2009, Aug 1). The Crumbs complex: from epithelial-cell polarity to retinal degeneration. J Cell Sci, 122(Pt 15), 2587–2596. https://doi.org/10.1242/jcs.023648

Bulik-Sullivan, B. K., Loh, P. R., Finucane, H. K., Ripke, S., Yang, J., Schizophrenia Working Group of the Psychiatric Genomics, C., Patterson, N., Daly, M. J., Price, A. L., & Neale, B. M. (2015, Mar). LD Score regression distinguishes confounding from polygenicity in genome-wide association studies. Nat Genet, 47(3), 291–295. https://doi.org/10.1038/ng.3211

Cebola, I., Rodriguez-Segui, S. A., Cho, C. H., Bessa, J., Rovira, M., Luengo, M., Chhatriwala, M., Berry, A., Ponsa-Cobas, J., Maestro, M. A., Jennings, R. E., Pasquali, L., Moran, I., Castro, N., Hanley, N. A., Gomez-Skarmeta, J. L., Vallier, L., & Ferrer, J. (2015, May). TEAD and YAP regulate the enhancer network of human embryonic pancreatic progenitors. Nat Cell Biol, 17(5), 615–626. https://doi.org/10.1038/ncb3160

Chiou, J., Zeng, C., Cheng, Z., Han, J. Y., Schlichting, M., Huang, S., Wang, J., Sui, Y., Deogaygay, A., Okino, M.-L., Qiu, Y., Sun, Y., Kudtarkar, P., Fang, R., Preissl, S., Sander, M., Gorkin, D., & Gaulton, K. J. (2019). Single cell chromatin accessibility reveals pancreatic islet cell type- and state-specific regulatory programs of diabetes risk. bioRxiv, 693671. https://doi.org/10.1101/693671

Claussnitzer, M., Dankel, S. N., Klocke, B., Grallert, H., Glunk, V., Berulava, T., Lee, H., Oskolkov, N., Fadista, J., Ehlers, K., Wahl, S., Hoffmann, C., Qian, K., Ronn, T., Riess, H., Muller-Nurasyid, M., Bretschneider, N., Schroeder, T., Skurk, T., Horsthemke, B., Diagram+Consortium, Spieler, D., Klingenspor, M., Seifert, M., Kern, M. J., Mejhert, N., Dahlman, I., Hansson, O., Hauck, S. M., Bluher, M., Arner, P., Groop, L., Illig, T., Suhre, K., Hsu, Y. H., Mellgren, G., Hauner, H., & Laumen, H. (2014, Jan 16). Leveraging cross-species transcription factor binding site patterns: from diabetes risk loci to disease mechanisms. Cell, 156(1-2), 343–358. https://doi.org/10.1016/j.cell.2013.10.058

Conrad, E., Stein, R., & Hunter, C. S. (2014, Aug). Revealing transcription factors during human pancreatic beta cell development. Trends Endocrinol Metab, 25(8), 407–414. https://doi.org/10.1016/j.tem.2014.03.013

Consortium, E. P. (2012, Sep 6). An integrated encyclopedia of DNA elements in the human genome. Nature, 489(7414), 57–74. https://doi.org/10.1038/nature11247

Dekker, J., Belmont, A. S., Guttman, M., Leshyk, V. O., Lis, J. T., Lomvardas, S., Mirny, L. A., O’Shea, C. C., Park, P. J., Ren, B., Politz, J. C. R., Shendure, J., Zhong, S., & Network, D. N. (2017, Sep 13). The 4D nucleome project. Nature, 549(7671), 219–226. https://doi.org/10.1038/nature23884

Dixon, J. R., Jung, I., Selvaraj, S., Shen, Y., Antosiewicz-Bourget, J. E., Lee, A. Y., Ye, Z., Kim, A., Rajagopal, N., Xie, W., Diao, Y., Liang, J., Zhao, H., Lobanenkov, V. V., Ecker, J. R., Thomson, J. A., & Ren, B. (2015, Feb 19). Chromatin architecture reorganization during stem cell differentiation. Nature, 518(7539), 331–336. https://doi.org/10.1038/nature14222

Dobin, A., Davis, C. A., Schlesinger, F., Drenkow, J., Zaleski, C., Jha, S., Batut, P., Chaisson, M., & Gingeras, T. R. (2013, Jan 1). STAR: ultrafast universal RNA-seq aligner. Bioinformatics, 29(1), 15–21. https://doi.org/10.1093/bioinformatics/bts635

Dong, P. D., Provost, E., Leach, S. D., & Stainier, D. Y. (2008, Jun 1). Graded levels of Ptf1a differentially regulate endocrine and exocrine fates in the developing pancreas. Genes Dev, 22(11), 1445–1450. https://doi.org/10.1101/gad.1663208

Dudok, J. J., Murtaza, M., Henrique Alves, C., Rashbass, P., & Wijnholds, J. (2016, Jul). Crumbs 2 prevents cortical abnormalities in mouse dorsal telencephalon. Neurosci Res, 108, 12–23. https://doi.org/10.1016/j.neures.2016.01.001

Dupuis, J., Langenberg, C., Prokopenko, I., Saxena, R., Soranzo, N., Jackson, A. U., Wheeler, E., Glazer, N. L., Bouatia-Naji, N., Gloyn, A. L., Lindgren, C. M., Magi, R., Morris, A. P., Randall, J., Johnson, T., Elliott, P., Rybin, D., Thorleifsson, G., Steinthorsdottir, V., Henneman, P., Grallert, H., Dehghan, A., Hottenga, J. J., Franklin, C. S., Navarro, P., Song, K., Goel, A., Perry, J. R., Egan, J. M., Lajunen, T., Grarup, N., Sparso, T., Doney, A., Voight, B. F., Stringham, H. M., Li, M., Kanoni, S., Shrader, P., Cavalcanti-Proenca, C., Kumari, M., Qi, L., Timpson, N. J., Gieger, C., Zabena, C., Rocheleau, G., Ingelsson, E., An, P., O’Connell, J., Luan, J., Elliott, A., McCarroll, S. A., Payne, F., Roccasecca, R. M., Pattou, F., Sethupathy, P., Ardlie, K., Ariyurek, Y., Balkau, B., Barter, P., Beilby, J. P., Ben-Shlomo, Y., Benediktsson, R., Bennett, A. J., Bergmann, S., Bochud, M., Boerwinkle, E., Bonnefond, A., Bonnycastle, L. L., Borch-Johnsen, K., Bottcher, Y., Brunner, E., Bumpstead, S. J., Charpentier, G., Chen, Y. D., Chines, P., Clarke, R., Coin, L. J., Cooper, M. N., Cornelis, M., Crawford, G., Crisponi, L., Day, I. N., de Geus, E. J., Delplanque, J., Dina, C., Erdos, M. R., Fedson, A. C., Fischer-Rosinsky, A., Forouhi, N. G., Fox, C. S., Frants, R., Franzosi, M. G., Galan, P., Goodarzi, M. O., Graessler, J., Groves, C. J., Grundy, S., Gwilliam, R., Gyllensten, U., Hadjadj, S., Hallmans, G., Hammond, N., Han, X., Hartikainen, A. L., Hassanali, N., Hayward, C., Heath, S. C., Hercberg, S., Herder, C., Hicks, A. A., Hillman, D. R., Hingorani, A. D., Hofman, A., Hui, J., Hung, J., Isomaa, B., Johnson, P. R., Jorgensen, T., Jula, A., Kaakinen, M., Kaprio, J., Kesaniemi, Y. A., Kivimaki, M., Knight, B., Koskinen, S., Kovacs, P., Kyvik, K. O., Lathrop, G. M., Lawlor, D. A., Le Bacquer, O., Lecoeur, C., Li, Y., Lyssenko, V., Mahley, R., Mangino, M., Manning, A. K., Martinez-Larrad, M. T., McAteer, J. B., McCulloch, L. J., McPherson, R., Meisinger, C., Melzer, D., Meyre, D., Mitchell, B. D., Morken, M. A., Mukherjee, S., Naitza, S., Narisu, N., Neville, M. J., Oostra, B. A., Orru, M., Pakyz, R., Palmer, C. N., Paolisso, G., Pattaro, C., Pearson, D., Peden, J. F., Pedersen, N. L., Perola, M., Pfeiffer, A. F., Pichler, I., Polasek, O., Posthuma, D., Potter, S. C., Pouta, A., Province, M. A., Psaty, B. M., Rathmann, W., Rayner, N. W., Rice, K., Ripatti, S., Rivadeneira, F., Roden, M., Rolandsson, O., Sandbaek, A., Sandhu, M., Sanna, S., Sayer, A. A., Scheet, P., Scott, L. J., Seedorf, U., Sharp, S. J., Shields, B., Sigurethsson, G., Sijbrands, E. J., Silveira, A., Simpson, L., Singleton, A., Smith, N. L., Sovio, U., Swift, A., Syddall, H., Syvanen, A. C., Tanaka, T., Thorand, B., Tichet, J., Tonjes, A., Tuomi, T., Uitterlinden, A. G., van Dijk, K. W., van Hoek, M., Varma, D., Visvikis-Siest, S., Vitart, V., Vogelzangs, N., Waeber, G., Wagner, P. J., Walley, A., Walters, G. B., Ward, K. L., Watkins, H., Weedon, M. N., Wild, S. H., Willemsen, G., Witteman, J. C., Yarnell, J. W., Zeggini, E., Zelenika, D., Zethelius, B., Zhai, G., Zhao, J. H., Zillikens, M. C., Consortium, D., Consortium, G., Global, B. C., Borecki, I. B., Loos, R. J., Meneton, P., Magnusson, P. K., Nathan, D. M., Williams, G. H., Hattersley, A. T., Silander, K., Salomaa, V., Smith, G. D., Bornstein, S. R., Schwarz, P., Spranger, J., Karpe, F., Shuldiner, A. R., Cooper, C., Dedoussis, G. V., Serrano-Rios, M., Morris, A. D., Lind, L., Palmer, L. J., Hu, F. B., Franks, P. W., Ebrahim, S., Marmot, M., Kao, W. H., Pankow, J. S., Sampson, M. J., Kuusisto, J., Laakso, M., Hansen, T., Pedersen, O., Pramstaller, P. P., Wichmann, H. E., Illig, T., Rudan, I., Wright, A. F., Stumvoll, M., Campbell, H., Wilson, J. F., Anders Hamsten on behalf of Procardis, C., investigators, M., Bergman, R. N., Buchanan, T. A., Collins, F. S., Mohlke, K. L., Tuomilehto, J., Valle, T. T., Altshuler, D., Rotter, J. I., Siscovick, D. S., Penninx, B. W., Boomsma, D. I., Deloukas, P., Spector, T. D., Frayling, T. M., Ferrucci, L., Kong, A., Thorsteinsdottir, U., Stefansson, K., van Duijn, C. M., Aulchenko, Y. S., Cao, A., Scuteri, A., Schlessinger, D., Uda, M., Ruokonen, A., Jarvelin, M. R., Waterworth, D. M., Vollenweider, P., Peltonen, L., Mooser, V., Abecasis, G. R., Wareham, N. J., Sladek, R., Froguel, P., Watanabe, R. M., Meigs, J. B., Groop, L., Boehnke, M., McCarthy, M. I., Florez, J. C., & Barroso, I. (2010, Feb). New genetic loci implicated in fasting glucose homeostasis and their impact on type 2 diabetes risk. Nat Genet, 42(2), 105–116. https://doi.org/10.1038/ng.520

Durand, N. C., Robinson, J. T., Shamim, M. S., Machol, I., Mesirov, J. P., Lander, E. S., & Aiden, E. L. (2016, Jul). Juicebox Provides a Visualization System for Hi-C Contact Maps with Unlimited Zoom. Cell Syst, 3(1), 99–101. https://doi.org/10.1016/j.cels.2015.07.012

Ernst, J., & Kellis, M. (2012, Feb 28). ChromHMM: automating chromatin-state discovery and characterization. Nat Methods, 9(3), 215–216. https://doi.org/10.1038/nmeth.1906

Field, H. A., Dong, P. D., Beis, D., & Stainier, D. Y. (2003, Sep 1). Formation of the digestive system in zebrafish. II. Pancreas morphogenesis. Dev Biol, 261(1), 197–208. https://doi.org/10.1016/s0012-1606(03)00308-7

Finucane, H. K., Bulik-Sullivan, B., Gusev, A., Trynka, G., Reshef, Y., Loh, P. R., Anttila, V., Xu, H., Zang, C., Farh, K., Ripke, S., Day, F. R., ReproGen, C., Schizophrenia Working Group of the Psychiatric Genomics, C., Consortium, R., Purcell, S., Stahl, E., Lindstrom, S., Perry, J. R., Okada, Y., Raychaudhuri, S., Daly, M. J., Patterson, N., Neale, B. M., & Price, A. L. (2015, Nov). Partitioning heritability by functional annotation using genome-wide association summary statistics. Nat Genet, 47(11), 1228–1235. https://doi.org/10.1038/ng.3404

Flannick, J., Mercader, J. M., Fuchsberger, C., Udler, M. S., Mahajan, A., Wessel, J., Teslovich, T. M., Caulkins, L., Koesterer, R., Barajas-Olmos, F., Blackwell, T. W., Boerwinkle, E., Brody, J. A., Centeno-Cruz, F., Chen, L., Chen, S., Contreras-Cubas, C., Cordova, E., Correa, A., Cortes, M., DeFronzo, R. A., Dolan, L., Drews, K. L., Elliott, A., Floyd, J. S., Gabriel, S., Garay-Sevilla, M. E., Garcia-Ortiz, H., Gross, M., Han, S., Heard-Costa, N. L., Jackson, A. U., Jorgensen, M. E., Kang, H. M., Kelsey, M., Kim, B. J., Koistinen, H. A., Kuusisto, J., Leader, J. B., Linneberg, A., Liu, C. T., Liu, J., Lyssenko, V., Manning, A. K., Marcketta, A., Malacara-Hernandez, J. M., Martinez-Hernandez, A., Matsuo, K., Mayer-Davis, E., Mendoza-Caamal, E., Mohlke, K. L., Morrison, A. C., Ndungu, A., Ng, M. C. Y., O’Dushlaine, C., Payne, A. J., Pihoker, C., Broad Genomics, P., Post, W. S., Preuss, M., Psaty, B. M., Vasan, R. S., Rayner, N. W., Reiner, A. P., Revilla-Monsalve, C., Robertson, N. R., Santoro, N., Schurmann, C., So, W. Y., Soberon, X., Stringham, H. M., Strom, T. M., Tam, C. H. T., Thameem, F., Tomlinson, B., Torres, J. M., Tracy, R. P., van Dam, R. M., Vujkovic, M., Wang, S., Welch, R. P., Witte, D. R., Wong, T. Y., Atzmon, G., Barzilai, N., Blangero, J., Bonnycastle, L. L., Bowden, D. W., Chambers, J. C., Chan, E., Cheng, C. Y., Cho, Y. S., Collins, F. S., de Vries, P. S., Duggirala, R., Glaser, B., Gonzalez, C., Gonzalez, M. E., Groop, L., Kooner, J. S., Kwak, S. H., Laakso, M., Lehman, D. M., Nilsson, P., Spector, T. D., Tai, E. S., Tuomi, T., Tuomilehto, J., Wilson, J. G., Aguilar-Salinas, C. A., Bottinger, E., Burke, B., Carey, D. J., Chan, J. C. N., Dupuis, J., Frossard, P., Heckbert, S. R., Hwang, M. Y., Kim, Y. J., Kirchner, H. L., Lee, J. Y., Lee, J., Loos, R. J. F., Ma, R. C. W., Morris, A. D., O’Donnell, C. J., Palmer, C. N. A., Pankow, J., Park, K. S., Rasheed, A., Saleheen, D., Sim, X., Small, K. S., Teo, Y. Y., Haiman, C., Hanis, C. L., Henderson, B. E., Orozco, L., Tusie-Luna, T., Dewey, F. E., Baras, A., Gieger, C., Meitinger, T., Strauch, K., Lange, L., Grarup, N., Hansen, T., Pedersen, O., Zeitler, P., Dabelea, D., Abecasis, G., Bell, G. I., Cox, N. J., Seielstad, M., Sladek, R., Meigs, J. B., Rich, S. S., Rotter, J. I., Discov, E. H. R. C., Charge, LuCamp, ProDiGy, GoT2D, Esp, Sigma, T. D., T2D, G., Amp T2D, G., Altshuler, D., Burtt, N. P., Scott, L. J., Morris, A. P., Florez, J. C., McCarthy, M. I., & Boehnke, M. (2019, Jun). Exome sequencing of 20,791 cases of type 2 diabetes and 24,440 controls. Nature, 570(7759), 71–76. https://doi.org/10.1038/s41586-019-1231-2

Fuchsberger, C., Flannick, J., Teslovich, T. M., Mahajan, A., Agarwala, V., Gaulton, K. J., Ma, C., Fontanillas, P., Moutsianas, L., McCarthy, D. J., Rivas, M. A., Perry, J. R. B., Sim, X., Blackwell, T. W., Robertson, N. R., Rayner, N. W., Cingolani, P., Locke, A. E., Tajes, J. F., Highland, H. M., Dupuis, J., Chines, P. S., Lindgren, C. M., Hartl, C., Jackson, A. U., Chen, H., Huyghe, J. R., van de Bunt, M., Pearson, R. D., Kumar, A., Muller-Nurasyid, M., Grarup, N., Stringham, H. M., Gamazon, E. R., Lee, J., Chen, Y., Scott, R. A., Below, J. E., Chen, P., Huang, J., Go, M. J., Stitzel, M. L., Pasko, D., Parker, S. C. J., Varga, T. V., Green, T., Beer, N. L., Day-Williams, A. G., Ferreira, T., Fingerlin, T., Horikoshi, M., Hu, C., Huh, I., Ikram, M. K., Kim, B. J., Kim, Y., Kim, Y. J., Kwon, M. S., Lee, J., Lee, S., Lin, K. H., Maxwell, T. J., Nagai, Y., Wang, X., Welch, R. P., Yoon, J., Zhang, W., Barzilai, N., Voight, B. F., Han, B. G., Jenkinson, C. P., Kuulasmaa, T., Kuusisto, J., Manning, A., Ng, M. C. Y., Palmer, N. D., Balkau, B., Stancakova, A., Abboud, H. E., Boeing, H., Giedraitis, V., Prabhakaran, D., Gottesman, O., Scott, J., Carey, J., Kwan, P., Grant, G., Smith, J. D., Neale, B. M., Purcell, S., Butterworth, A. S., Howson, J. M. M., Lee, H. M., Lu, Y., Kwak, S. H., Zhao, W., Danesh, J., Lam, V. K. L., Park, K. S., Saleheen, D., So, W. Y., Tam, C. H. T., Afzal, U., Aguilar, D., Arya, R., Aung, T., Chan, E., Navarro, C., Cheng, C. Y., Palli, D., Correa, A., Curran, J. E., Rybin, D., Farook, V. S., Fowler, S. P., Freedman, B. I., Griswold, M., Hale, D. E., Hicks, P. J., Khor, C. C., Kumar, S., Lehne, B., Thuillier, D., Lim, W. Y., Liu, J., van der Schouw, Y. T., Loh, M., Musani, S. K., Puppala, S., Scott, W. R., Yengo, L., Tan, S. T., Taylor, H. A., Jr., Thameem, F., Wilson, G., Sr., Wong, T. Y., Njolstad, P. R., Levy, J. C., Mangino, M., Bonnycastle, L. L., Schwarzmayr, T., Fadista, J., Surdulescu, G. L., Herder, C., Groves, C. J., Wieland, T., Bork-Jensen, J., Brandslund, I., Christensen, C., Koistinen, H. A., Doney, A. S. F., Kinnunen, L., Esko, T., Farmer, A. J., Hakaste, L., Hodgkiss, D., Kravic, J., Lyssenko, V., Hollensted, M., Jorgensen, M. E., Jorgensen, T., Ladenvall, C., Justesen, J. M., Karajamaki, A., Kriebel, J., Rathmann, W., Lannfelt, L., Lauritzen, T., Narisu, N., Linneberg, A., Melander, O., Milani, L., Neville, M., Orho-Melander, M., Qi, L., Qi, Q., Roden, M., Rolandsson, O., Swift, A., Rosengren, A. H., Stirrups, K., Wood, A. R., Mihailov, E., Blancher, C., Carneiro, M. O., Maguire, J., Poplin, R., Shakir, K., Fennell, T., DePristo, M., de Angelis, M. H., Deloukas, P., Gjesing, A. P., Jun, G., Nilsson, P., Murphy, J., Onofrio, R., Thorand, B., Hansen, T., Meisinger, C., Hu, F. B., Isomaa, B., Karpe, F., Liang, L., Peters, A., Huth, C., O’Rahilly, S. P., Palmer, C. N. A., Pedersen, O., Rauramaa, R., Tuomilehto, J., Salomaa, V., Watanabe, R. M., Syvanen, A. C., Bergman, R. N., Bharadwaj, D., Bottinger, E. P., Cho, Y. S., Chandak, G. R., Chan, J. C. N., Chia, K. S., Daly, M. J., Ebrahim, S. B., Langenberg, C., Elliott, P., Jablonski, K. A., Lehman, D. M., Jia, W., Ma, R. C. W., Pollin, T. I., Sandhu, M., Tandon, N., Froguel, P., Barroso, I., Teo, Y. Y., Zeggini, E., Loos, R. J. F., Small, K. S., Ried, J. S., DeFronzo, R. A., Grallert, H., Glaser, B., Metspalu, A., Wareham, N. J., Walker, M., Banks, E., Gieger, C., Ingelsson, E., Im, H. K., Illig, T., Franks, P. W., Buck, G., Trakalo, J., Buck, D., Prokopenko, I., Magi, R., Lind, L., Farjoun, Y., Owen, K. R., Gloyn, A. L., Strauch, K., Tuomi, T., Kooner, J. S., Lee, J. Y., Park, T., Donnelly, P., Morris, A. D., Hattersley, A. T., Bowden, D. W., Collins, F. S., Atzmon, G., Chambers, J. C., Spector, T. D., Laakso, M., Strom, T. M., Bell, G. I., Blangero, J., Duggirala, R., Tai, E. S., McVean, G., Hanis, C. L., Wilson, J. G., Seielstad, M., Frayling, T. M., Meigs, J. B., Cox, N. J., Sladek, R., Lander, E. S., Gabriel, S., Burtt, N. P., Mohlke, K. L., Meitinger, T., Groop, L., Abecasis, G., Florez, J. C., Scott, L. J., Morris, A. P., Kang, H. M., Boehnke, M., Altshuler, D., & McCarthy, M. I. (2016, Aug 4). The genetic architecture of type 2 diabetes. Nature, 536(7614), 41–47. https://doi.org/10.1038/nature18642

Gaertner, B., Carrano, A. C., & Sander, M. (2019, Nov 1). Human stem cell models: lessons for pancreatic development and disease. Genes Dev, 33(21-22), 1475–1490. https://doi.org/10.1101/gad.331397.119

Gaulton, K. J., Ferreira, T., Lee, Y., Raimondo, A., Magi, R., Reschen, M. E., Mahajan, A., Locke, A., Rayner, N. W., Robertson, N., Scott, R. A., Prokopenko, I., Scott, L. J., Green, T., Sparso, T., Thuillier, D., Yengo, L., Grallert, H., Wahl, S., Franberg, M., Strawbridge, R. J., Kestler, H., Chheda, H., Eisele, L., Gustafsson, S., Steinthorsdottir, V., Thorleifsson, G., Qi, L., Karssen, L. C., van Leeuwen, E. M., Willems, S. M., Li, M., Chen, H., Fuchsberger, C., Kwan, P., Ma, C., Linderman, M., Lu, Y., Thomsen, S. K., Rundle, J. K., Beer, N. L., van de Bunt, M., Chalisey, A., Kang, H. M., Voight, B. F., Abecasis, G. R., Almgren, P., Baldassarre, D., Balkau, B., Benediktsson, R., Bluher, M., Boeing, H., Bonnycastle, L. L., Bottinger, E. P., Burtt, N. P., Carey, J., Charpentier, G., Chines, P. S., Cornelis, M. C., Couper, D. J., Crenshaw, A. T., van Dam, R. M., Doney, A. S., Dorkhan, M., Edkins, S., Eriksson, J. G., Esko, T., Eury, E., Fadista, J., Flannick, J., Fontanillas, P., Fox, C., Franks, P. W., Gertow, K., Gieger, C., Gigante, B., Gottesman, O., Grant, G. B., Grarup, N., Groves, C. J., Hassinen, M., Have, C. T., Herder, C., Holmen, O. L., Hreidarsson, A. B., Humphries, S. E., Hunter, D. J., Jackson, A. U., Jonsson, A., Jorgensen, M. E., Jorgensen, T., Kao, W. H., Kerrison, N. D., Kinnunen, L., Klopp, N., Kong, A., Kovacs, P., Kraft, P., Kravic, J., Langford, C., Leander, K., Liang, L., Lichtner, P., Lindgren, C. M., Lindholm, E., Linneberg, A., Liu, C. T., Lobbens, S., Luan, J., Lyssenko, V., Mannisto, S., McLeod, O., Meyer, J., Mihailov, E., Mirza, G., Muhleisen, T. W., Muller-Nurasyid, M., Navarro, C., Nothen, M. M., Oskolkov, N. N., Owen, K. R., Palli, D., Pechlivanis, S., Peltonen, L., Perry, J. R., Platou, C. G., Roden, M., Ruderfer, D., Rybin, D., van der Schouw, Y. T., Sennblad, B., Sigurethsson, G., Stancakova, A., Steinbach, G., Storm, P., Strauch, K., Stringham, H. M., Sun, Q., Thorand, B., Tikkanen, E., Tonjes, A., Trakalo, J., Tremoli, E., Tuomi, T., Wennauer, R., Wiltshire, S., Wood, A. R., Zeggini, E., Dunham, I., Birney, E., Pasquali, L., Ferrer, J., Loos, R. J., Dupuis, J., Florez, J. C., Boerwinkle, E., Pankow, J. S., van Duijn, C., Sijbrands, E., Meigs, J. B., Hu, F. B., Thorsteinsdottir, U., Stefansson, K., Lakka, T. A., Rauramaa, R., Stumvoll, M., Pedersen, N. L., Lind, L., Keinanen-Kiukaanniemi, S. M., Korpi-Hyovalti, E., Saaristo, T. E., Saltevo, J., Kuusisto, J., Laakso, M., Metspalu, A., Erbel, R., Jocke, K. H., Moebus, S., Ripatti, S., Salomaa, V., Ingelsson, E., Boehm, B. O., Bergman, R. N., Collins, F. S., Mohlke, K. L., Koistinen, H., Tuomilehto, J., Hveem, K., Njolstad, I., Deloukas, P., Donnelly, P. J., Frayling, T. M., Hattersley, A. T., de Faire, U., Hamsten, A., Illig, T., Peters, A., Cauchi, S., Sladek, R., Froguel, P., Hansen, T., Pedersen, O., Morris, A. D., Palmer, C. N., Kathiresan, S., Melander, O., Nilsson, P. M., Groop, L. C., Barroso, I., Langenberg, C., Wareham, N. J., O’Callaghan, C. A., Gloyn, A. L., Altshuler, D., Boehnke, M., Teslovich, T. M., McCarthy, M. I., Morris, A. P., Replication, D. I. G., & Meta-analysis, C. (2015, Dec). Genetic fine mapping and genomic annotation defines causal mechanisms at type 2 diabetes susceptibility loci. Nat Genet, 47(12), 1415–1425. https://doi.org/10.1038/ng.3437

Gaulton, K. J., Nammo, T., Pasquali, L., Simon, J. M., Giresi, P. G., Fogarty, M. P., Panhuis, T. M., Mieczkowski, P., Secchi, A., Bosco, D., Berney, T., Montanya, E., Mohlke, K. L., Lieb, J. D., & Ferrer, J. (2010, Mar). A map of open chromatin in human pancreatic islets. Nat Genet, 42(3), 255–259. https://doi.org/10.1038/ng.530

Godinho, L., Mumm, J. S., Williams, P. R., Schroeter, E. H., Koerber, A., Park, S. W., Leach, S. D., & Wong, R. O. (2005, Nov). Targeting of amacrine cell neurites to appropriate synaptic laminae in the developing zebrafish retina. Development, 132(22), 5069–5079. https://doi.org/10.1242/dev.02075

Grant, C. E., Bailey, T. L., & Noble, W. S. (2011, Apr 1). FIMO: scanning for occurrences of a given motif. Bioinformatics, 27(7), 1017–1018. https://doi.org/10.1093/bioinformatics/btr064

Greenwald, W. W., Chiou, J., Yan, J., Qiu, Y., Dai, N., Wang, A., Nariai, N., Aylward, A., Han, J. Y., Kadakia, N., Regue, L., Okino, M. L., Drees, F., Kramer, D., Vinckier, N., Minichiello, L., Gorkin, D., Avruch, J., Frazer, K. A., Sander, M., Ren, B., & Gaulton, K. J. (2019, May 7). Pancreatic islet chromatin accessibility and conformation reveals distal enhancer networks of type 2 diabetes risk. Nat Commun, 10(1), 2078. https://doi.org/10.1038/s41467-019-09975-4

Halban, P. A., Polonsky, K. S., Bowden, D. W., Hawkins, M. A., Ling, C., Mather, K. J., Powers, A. C., Rhodes, C. J., Sussel, L., & Weir, G. C. (2014, Jun). beta-cell failure in type 2 diabetes: postulated mechanisms and prospects for prevention and treatment. Diabetes Care, 37(6), 1751–1758. https://doi.org/10.2337/dc14-0396

Heinz, S., Benner, C., Spann, N., Bertolino, E., Lin, Y. C., Laslo, P., Cheng, J. X., Murre, C., Singh, H., & Glass, C. K. (2010, May 28). Simple combinations of lineage-determining transcription factors prime cis-regulatory elements required for macrophage and B cell identities. Mol Cell, 38(4), 576–589. https://doi.org/10.1016/j.molcel.2010.05.004

Helker, C. S. M., Mullapudi, S. T., Mueller, L. M., Preussner, J., Tunaru, S., Skog, O., Kwon, H. B., Kreuder, F., Lancman, J. J., Bonnavion, R., Dong, P. D. S., Looso, M., Offermanns, S., Korsgren, O., Spagnoli, F. M., & Stainier, D. Y. R. (2019, Jul 24). A whole organism small molecule screen identifies novel regulators of pancreatic endocrine development. Development, 146(14). https://doi.org/10.1242/dev.172569

Heymans, C., Degosserie, J., Spourquet, C., & Pierreux, C. E. (2019, Feb 25). Pancreatic acinar differentiation is guided by differential laminin deposition. Sci Rep, 9(1), 2711. https://doi.org/10.1038/s41598-019-39077-6

Horikoshi, M., Beaumont, R. N., Day, F. R., Warrington, N. M., Kooijman, M. N., Fernandez-Tajes, J., Feenstra, B., van Zuydam, N. R., Gaulton, K. J., Grarup, N., Bradfield, J. P., Strachan, D. P., Li-Gao, R., Ahluwalia, T. S., Kreiner, E., Rueedi, R., Lyytikainen, L. P., Cousminer, D. L., Wu, Y., Thiering, E., Wang, C. A., Have, C. T., Hottenga, J. J., Vilor-Tejedor, N., Joshi, P. K., Boh, E. T. H., Ntalla, I., Pitkanen, N., Mahajan, A., van Leeuwen, E. M., Joro, R., Lagou, V., Nodzenski, M., Diver, L. A., Zondervan, K. T., Bustamante, M., Marques-Vidal, P., Mercader, J. M., Bennett, A. J., Rahmioglu, N., Nyholt, D. R., Ma, R. C. W., Tam, C. H. T., Tam, W. H., Group, C. C. H. W., Ganesh, S. K., van Rooij, F. J., Jones, S. E., Loh, P. R., Ruth, K. S., Tuke, M. A., Tyrrell, J., Wood, A. R., Yaghootkar, H., Scholtens, D. M., Paternoster, L., Prokopenko, I., Kovacs, P., Atalay, M., Willems, S. M., Panoutsopoulou, K., Wang, X., Carstensen, L., Geller, F., Schraut, K. E., Murcia, M., van Beijsterveldt, C. E., Willemsen, G., Appel, E. V. R., Fonvig, C. E., Trier, C., Tiesler, C. M., Standl, M., Kutalik, Z., Bonas-Guarch, S., Hougaard, D. M., Sanchez, F., Torrents, D., Waage, J., Hollegaard, M. V., de Haan, H. G., Rosendaal, F. R., Medina-Gomez, C., Ring, S. M., Hemani, G., McMahon, G., Robertson, N. R., Groves, C. J., Langenberg, C., Luan, J., Scott, R. A., Zhao, J. H., Mentch, F. D., MacKenzie, S. M., Reynolds, R. M., Early Growth Genetics, C., Lowe, W. L., Jr., Tonjes, A., Stumvoll, M., Lindi, V., Lakka, T. A., van Duijn, C. M., Kiess, W., Korner, A., Sorensen, T. I., Niinikoski, H., Pahkala, K., Raitakari, O. T., Zeggini, E., Dedoussis, G. V., Teo, Y. Y., Saw, S. M., Melbye, M., Campbell, H., Wilson, J. F., Vrijheid, M., de Geus, E. J., Boomsma, D. I., Kadarmideen, H. N., Holm, J. C., Hansen, T., Sebert, S., Hattersley, A. T., Beilin, L. J., Newnham, J. P., Pennell, C. E., Heinrich, J., Adair, L. S., Borja, J. B., Mohlke, K. L., Eriksson, J. G., Widen, E. E., Kahonen, M., Viikari, J. S., Lehtimaki, T., Vollenweider, P., Bonnelykke, K., Bisgaard, H., Mook-Kanamori, D. O., Hofman, A., Rivadeneira, F., Uitterlinden, A. G., Pisinger, C., Pedersen, O., Power, C., Hypponen, E., Wareham, N. J., Hakonarson, H., Davies, E., Walker, B. R., Jaddoe, V. W., Jarvelin, M. R., Grant, S. F., Vaag, A. A., Lawlor, D. A., Frayling, T. M., Davey Smith, G., Morris, A. P., Ong, K. K., Felix, J. F., Timpson, N. J., Perry, J. R., Evans, D. M., McCarthy, M. I., & Freathy, R. M. (2016, Oct 13). Genome-wide associations for birth weight and correlations with adult disease. Nature, 538(7624), 248–252. https://doi.org/10.1038/nature19806

Hsu, Y. C., & Jensen, A. M. (2010, Jul 29). Multiple domains in the Crumbs Homolog 2a (Crb2a) protein are required for regulating rod photoreceptor size. BMC Cell Biol, 11, 60. https://doi.org/10.1186/1471-2121-11-60

Jimenez-Amilburu, V., & Stainier, D. Y. R. (2019, May 7). The transmembrane protein Crb2a regulates cardiomyocyte apicobasal polarity and adhesion in zebrafish. Development, 146(9). https://doi.org/10.1242/dev.171207

Jin, W., Mulas, F., Gaertner, B., Sui, Y., Wang, J., Matta, I., Zeng, C., Vinckier, N., Wang, A., Nguyen-Ngoc, K. V., Chiou, J., Kaestner, K. H., Frazer, K. A., Carrano, A. C., Shih, H. P., & Sander, M. (2019, Nov 22). A Network of microRNAs Acts to Promote Cell Cycle Exit and Differentiation of Human Pancreatic Endocrine Cells. iScience, 21, 681–694. https://doi.org/10.1016/j.isci.2019.10.063

Kent, W. J., Sugnet, C. W., Furey, T. S., Roskin, K. M., Pringle, T. H., Zahler, A. M., & Haussler, D. (2002, Jun). The human genome browser at UCSC. Genome Res, 12(6), 996–1006. https://doi.org/10.1101/gr.229102

Khetan, S., Kursawe, R., Youn, A., Lawlor, N., Jillette, A., Marquez, E. J., Ucar, D., & Stitzel, M. L. (2018, Nov). Type 2 Diabetes-Associated Genetic Variants Regulate Chromatin Accessibility in Human Islets. Diabetes, 67(11), 2466–2477. https://doi.org/10.2337/db18-0393

Kimmel, R. A., Dobler, S., Schmitner, N., Walsen, T., Freudenblum, J., & Meyer, D. (2015, Sep 18). Diabetic pdx1-mutant zebrafish show conserved responses to nutrient overload and anti-glycemic treatment. Sci Rep, 5, 14241. https://doi.org/10.1038/srep14241

Lancman, J. J., Zvenigorodsky, N., Gates, K. P., Zhang, D., Solomon, K., Humphrey, R. K., Kuo, T., Setiawan, L., Verkade, H., Chi, Y. I., Jhala, U. S., Wright, C. V., Stainier, D. Y., & Dong, P. D. (2013, Jul). Specification of hepatopancreas progenitors in zebrafish by hnf1ba and wnt2bb. Development, 140(13), 2669–2679. https://doi.org/10.1242/dev.090993

Li, H., & Durbin, R. (2009, Jul 15). Fast and accurate short read alignment with Burrows-Wheeler transform. Bioinformatics, 25(14), 1754–1760. https://doi.org/10.1093/bioinformatics/btp324

Li, H., Handsaker, B., Wysoker, A., Fennell, T., Ruan, J., Homer, N., Marth, G., Abecasis, G., Durbin, R., & Genome Project Data Processing, S. (2009, Aug 15). The Sequence Alignment/Map format and SAMtools. Bioinformatics, 25(16), 2078–2079. https://doi.org/10.1093/bioinformatics/btp352

Li, X. Y., Zhai, W. J., & Teng, C. B. (2015, Dec 30). Notch Signaling in Pancreatic Development. Int J Mol Sci, 17(1). https://doi.org/10.3390/ijms17010048

Love, M. I., Huber, W., & Anders, S. (2014). Moderated estimation of fold change and dispersion for RNA-seq data with DESeq2. Genome Biol, 15(12), 550. https://doi.org/10.1186/s13059-014-0550-8

Lumey, L. H., Khalangot, M. D., & Vaiserman, A. M. (2015, Oct). Association between type 2 diabetes and prenatal exposure to the Ukraine famine of 1932-33: a retrospective cohort study. Lancet Diabetes Endocrinol, 3(10), 787–794. https://doi.org/10.1016/S2213-8587(15)00279-X

Mahajan, A., Taliun, D., Thurner, M., Robertson, N. R., Torres, J. M., Rayner, N. W., Payne, A. J., Steinthorsdottir, V., Scott, R. A., Grarup, N., Cook, J. P., Schmidt, E. M., Wuttke, M., Sarnowski, C., Magi, R., Nano, J., Gieger, C., Trompet, S., Lecoeur, C., Preuss, M. H., Prins, B. P., Guo, X., Bielak, L. F., Below, J. E., Bowden, D. W., Chambers, J. C., Kim, Y. J., Ng, M. C. Y., Petty, L. E., Sim, X., Zhang, W., Bennett, A. J., Bork-Jensen, J., Brummett, C. M., Canouil, M., Ec Kardt, K. U., Fischer, K., Kardia, S. L. R., Kronenberg, F., Lall, K., Liu, C. T., Locke, A. E., Luan, J., Ntalla, I., Nylander, V., Schonherr, S., Schurmann, C., Yengo, L., Bottinger, E. P., Brandslund, I., Christensen, C., Dedoussis, G., Florez, J. C., Ford, I., Franco, O. H., Frayling, T. M., Giedraitis, V., Hackinger, S., Hattersley, A. T., Herder, C., Ikram, M. A., Ingelsson, M., Jorgensen, M. E., Jorgensen, T., Kriebel, J., Kuusisto, J., Ligthart, S., Lindgren, C. M., Linneberg, A., Lyssenko, V., Mamakou, V., Meitinger, T., Mohlke, K. L., Morris, A. D., Nadkarni, G., Pankow, J. S., Peters, A., Sattar, N., Stancakova, A., Strauch, K., Taylor, K. D., Thorand, B., Thorleifsson, G., Thorsteinsdottir, U., Tuomilehto, J., Witte, D. R., Dupuis, J., Peyser, P. A., Zeggini, E., Loos, R. J. F., Froguel, P., Ingelsson, E., Lind, L., Groop, L., Laakso, M., Collins, F. S., Jukema, J. W., Palmer, C. N. A., Grallert, H., Metspalu, A., Dehghan, A., Kottgen, A., Abecasis, G. R., Meigs, J. B., Rotter, J. I., Marchini, J., Pedersen, O., Hansen, T., Langenberg, C., Wareham, N. J., Stefansson, K., Gloyn, A. L., Morris, A. P., Boehnke, M., & McCarthy, M. I. (2018, Nov). Fine-mapping type 2 diabetes loci to single-variant resolution using high-density imputation and islet-specific epigenome maps. Nat Genet, 50(11), 1505–1513. https://doi.org/10.1038/s41588-018-0241-6

Mamidi, A., Prawiro, C., Seymour, P. A., de Lichtenberg, K. H., Jackson, A., Serup, P., & Semb, H. (2018, Dec). Mechanosignalling via integrins directs fate decisions of pancreatic progenitors. Nature, 564(7734), 114–118. https://doi.org/10.1038/s41586-018-0762-2

Manning, A. K., Hivert, M. F., Scott, R. A., Grimsby, J. L., Bouatia-Naji, N., Chen, H., Rybin, D., Liu, C. T., Bielak, L. F., Prokopenko, I., Amin, N., Barnes, D., Cadby, G., Hottenga, J. J., Ingelsson, E., Jackson, A. U., Johnson, T., Kanoni, S., Ladenvall, C., Lagou, V., Lahti, J., Lecoeur, C., Liu, Y., Martinez-Larrad, M. T., Montasser, M. E., Navarro, P., Perry, J. R., Rasmussen-Torvik, L. J., Salo, P., Sattar, N., Shungin, D., Strawbridge, R. J., Tanaka, T., van Duijn, C. M., An, P., de Andrade, M., Andrews, J. S., Aspelund, T., Atalay, M., Aulchenko, Y., Balkau, B., Bandinelli, S., Beckmann, J. S., Beilby, J. P., Bellis, C., Bergman, R. N., Blangero, J., Boban, M., Boehnke, M., Boerwinkle, E., Bonnycastle, L. L., Boomsma, D. I., Borecki, I. B., Bottcher, Y., Bouchard, C., Brunner, E., Budimir, D., Campbell, H., Carlson, O., Chines, P. S., Clarke, R., Collins, F. S., Corbaton-Anchuelo, A., Couper, D., de Faire, U., Dedoussis, G. V., Deloukas, P., Dimitriou, M., Egan, J. M., Eiriksdottir, G., Erdos, M. R., Eriksson, J. G., Eury, E., Ferrucci, L., Ford, I., Forouhi, N. G., Fox, C. S., Franzosi, M. G., Franks, P. W., Frayling, T. M., Froguel, P., Galan, P., de Geus, E., Gigante, B., Glazer, N. L., Goel, A., Groop, L., Gudnason, V., Hallmans, G., Hamsten, A., Hansson, O., Harris, T. B., Hayward, C., Heath, S., Hercberg, S., Hicks, A. A., Hingorani, A., Hofman, A., Hui, J., Hung, J., Jarvelin, M. R., Jhun, M. A., Johnson, P. C., Jukema, J. W., Jula, A., Kao, W. H., Kaprio, J., Kardia, S. L., Keinanen-Kiukaanniemi, S., Kivimaki, M., Kolcic, I., Kovacs, P., Kumari, M., Kuusisto, J., Kyvik, K. O., Laakso, M., Lakka, T., Lannfelt, L., Lathrop, G. M., Launer, L. J., Leander, K., Li, G., Lind, L., Lindstrom, J., Lobbens, S., Loos, R. J., Luan, J., Lyssenko, V., Magi, R., Magnusson, P. K., Marmot, M., Meneton, P., Mohlke, K. L., Mooser, V., Morken, M. A., Miljkovic, I., Narisu, N., O’Connell, J., Ong, K. K., Oostra, B. A., Palmer, L. J., Palotie, A., Pankow, J. S., Peden, J. F., Pedersen, N. L., Pehlic, M., Peltonen, L., Penninx, B., Pericic, M., Perola, M., Perusse, L., Peyser, P. A., Polasek, O., Pramstaller, P. P., Province, M. A., Raikkonen, K., Rauramaa, R., Rehnberg, E., Rice, K., Rotter, J. I., Rudan, I., Ruokonen, A., Saaristo, T., Sabater-Lleal, M., Salomaa, V., Savage, D. B., Saxena, R., Schwarz, P., Seedorf, U., Sennblad, B., Serrano-Rios, M., Shuldiner, A. R., Sijbrands, E. J., Siscovick, D. S., Smit, J. H., Small, K. S., Smith, N. L., Smith, A. V., Stancakova, A., Stirrups, K., Stumvoll, M., Sun, Y. V., Swift, A. J., Tonjes, A., Tuomilehto, J., Trompet, S., Uitterlinden, A. G., Uusitupa, M., Vikstrom, M., Vitart, V., Vohl, M. C., Voight, B. F., Vollenweider, P., Waeber, G., Waterworth, D. M., Watkins, H., Wheeler, E., Widen, E., Wild, S. H., Willems, S. M., Willemsen, G., Wilson, J. F., Witteman, J. C., Wright, A. F., Yaghootkar, H., Zelenika, D., Zemunik, T., Zgaga, L., Replication, D. I. G., Meta-analysis, C., Multiple Tissue Human Expression Resource, C., Wareham, N. J., McCarthy, M. I., Barroso, I., Watanabe, R. M., Florez, J. C., Dupuis, J., Meigs, J. B., & Langenberg, C. (2012, May 13). A genome-wide approach accounting for body mass index identifies genetic variants influencing fasting glycemic traits and insulin resistance. Nat Genet, 44(6), 659–669. https://doi.org/10.1038/ng.2274

Masui, T., Long, Q., Beres, T. M., Magnuson, M. A., & MacDonald, R. J. (2007, Oct 15). Early pancreatic development requires the vertebrate Suppressor of Hairless (RBPJ) in the PTF1 bHLH complex. Genes Dev, 21(20), 2629–2643. https://doi.org/10.1101/gad.1575207

Mathelier, A., Fornes, O., Arenillas, D. J., Chen, C. Y., Denay, G., Lee, J., Shi, W., Shyr, C., Tan, G., Worsley-Hunt, R., Zhang, A. W., Parcy, F., Lenhard, B., Sandelin, A., & Wasserman, W. W. (2016, Jan 4). JASPAR 2016: a major expansion and update of the open-access database of transcription factor binding profiles. Nucleic Acids Res, 44(D1), D110–115. https://doi.org/10.1093/nar/gkv1176

Mikkelsen, T. S., Xu, Z., Zhang, X., Wang, L., Gimble, J. M., Lander, E. S., & Rosen, E. D. (2010, Oct 1). Comparative epigenomic analysis of murine and human adipogenesis. Cell, 143(1), 156–169. https://doi.org/10.1016/j.cell.2010.09.006

Murtaugh, L. C. (2008, Apr). The what, where, when and how of Wnt/beta-catenin signaling in pancreas development. Organogenesis, 4(2), 81–86. https://doi.org/10.4161/org.4.2.5853

Nielsen, J. H., Haase, T. N., Jaksch, C., Nalla, A., Sostrup, B., Nalla, A. A., Larsen, L., Rasmussen, M., Dalgaard, L. T., Gaarn, L. W., Thams, P., Kofod, H., & Billestrup, N. (2014, Nov). Impact of fetal and neonatal environment on beta cell function and development of diabetes. Acta Obstet Gynecol Scand, 93(11), 1109–1122. https://doi.org/10.1111/aogs.12504

Nikolova, G., Jabs, N., Konstantinova, I., Domogatskaya, A., Tryggvason, K., Sorokin, L., Fassler, R., Gu, G., Gerber, H. P., Ferrara, N., Melton, D. A., & Lammert, E. (2006, Mar). The vascular basement membrane: a niche for insulin gene expression and Beta cell proliferation. Dev Cell, 10(3), 397–405. https://doi.org/10.1016/j.devcel.2006.01.015

Omori, Y., & Malicki, J. (2006, May 23). oko meduzy and related crumbs genes are determinants of apical cell features in the vertebrate embryo. Curr Biol, 16(10), 945–957. https://doi.org/10.1016/j.cub.2006.03.058

Parker, S. C., Stitzel, M. L., Taylor, D. L., Orozco, J. M., Erdos, M. R., Akiyama, J. A., van Bueren, K. L., Chines, P. S., Narisu, N., Program, N. C. S., Black, B. L., Visel, A., Pennacchio, L. A., Collins, F. S., National Institutes of Health Intramural Sequencing Center Comparative Sequencing Program, A., & Authors, N. C. S. P. (2013, Oct 29). Chromatin stretch enhancer states drive cell-specific gene regulation and harbor human disease risk variants. Proc Natl Acad Sci U S A, 110(44), 17921–17926. https://doi.org/10.1073/pnas.1317023110

Pasquali, L., Gaulton, K. J., Rodriguez-Segui, S. A., Mularoni, L., Miguel-Escalada, I., Akerman, I., Tena, J. J., Moran, I., Gomez-Marin, C., van de Bunt, M., Ponsa-Cobas, J., Castro, N., Nammo, T., Cebola, I., Garcia-Hurtado, J., Maestro, M. A., Pattou, F., Piemonti, L., Berney, T., Gloyn, A. L., Ravassard, P., Skarmeta, J. L. G., Muller, F., McCarthy, M. I., & Ferrer, J. (2014, Feb). Pancreatic islet enhancer clusters enriched in type 2 diabetes risk-associated variants. Nat Genet, 46(2), 136–143. https://doi.org/10.1038/ng.2870

Perry, J. R., Voight, B. F., Yengo, L., Amin, N., Dupuis, J., Ganser, M., Grallert, H., Navarro, P., Li, M., Qi, L., Steinthorsdottir, V., Scott, R. A., Almgren, P., Arking, D. E., Aulchenko, Y., Balkau, B., Benediktsson, R., Bergman, R. N., Boerwinkle, E., Bonnycastle, L., Burtt, N. P., Campbell, H., Charpentier, G., Collins, F. S., Gieger, C., Green, T., Hadjadj, S., Hattersley, A. T., Herder, C., Hofman, A., Johnson, A. D., Kottgen, A., Kraft, P., Labrune, Y., Langenberg, C., Manning, A. K., Mohlke, K. L., Morris, A. P., Oostra, B., Pankow, J., Petersen, A. K., Pramstaller, P. P., Prokopenko, I., Rathmann, W., Rayner, W., Roden, M., Rudan, I., Rybin, D., Scott, L. J., Sigurdsson, G., Sladek, R., Thorleifsson, G., Thorsteinsdottir, U., Tuomilehto, J., Uitterlinden, A. G., Vivequin, S., Weedon, M. N., Wright, A. F., Magic, Consortium, D., Consortium, G., Hu, F. B., Illig, T., Kao, L., Meigs, J. B., Wilson, J. F., Stefansson, K., van Duijn, C., Altschuler, D., Morris, A. D., Boehnke, M., McCarthy, M. I., Froguel, P., Palmer, C. N., Wareham, N. J., Groop, L., Frayling, T. M., & Cauchi, S. (2012, May). Stratifying type 2 diabetes cases by BMI identifies genetic risk variants in LAMA1 and enrichment for risk variants in lean compared to obese cases. PLoS Genet, 8(5), e1002741. https://doi.org/10.1371/journal.pgen.1002741

Pique-Regi, R., Degner, J. F., Pai, A. A., Gaffney, D. J., Gilad, Y., & Pritchard, J. K. (2011, Mar). Accurate inference of transcription factor binding from DNA sequence and chromatin accessibility data. Genome Res, 21(3), 447–455. https://doi.org/10.1101/gr.112623.110

Pollard, S. M., Parsons, M. J., Kamei, M., Kettleborough, R. N., Thomas, K. A., Pham, V. N., Bae, M. K., Scott, A., Weinstein, B. M., & Stemple, D. L. (2006, Jan 1). Essential and overlapping roles for laminin alpha chains in notochord and blood vessel formation. Dev Biol, 289(1), 64–76. https://doi.org/10.1016/j.ydbio.2005.10.006

Portha, B., Chavey, A., & Movassat, J. (2011). Early-life origins of type 2 diabetes: fetal programming of the beta-cell mass. Exp Diabetes Res, 2011, 105076. https://doi.org/10.1155/2011/105076

Quinlan, A. R., & Hall, I. M. (2010, Mar 15). BEDTools: a flexible suite of utilities for comparing genomic features. Bioinformatics, 26(6), 841–842. https://doi.org/10.1093/bioinformatics/btq033

Rada-Iglesias, A., Bajpai, R., Swigut, T., Brugmann, S. A., Flynn, R. A., & Wysocka, J. (2011, Feb 10). A unique chromatin signature uncovers early developmental enhancers in humans. Nature, 470(7333), 279–283. https://doi.org/10.1038/nature09692

Ramirez, F., Ryan, D. P., Gruning, B., Bhardwaj, V., Kilpert, F., Richter, A. S., Heyne, S., Dundar, F., & Manke, T. (2016, Jul 8). deepTools2: a next generation web server for deep-sequencing data analysis. Nucleic Acids Res, 44(W1), W160–165. https://doi.org/10.1093/nar/gkw257

Rao, S. S., Huntley, M. H., Durand, N. C., Stamenova, E. K., Bochkov, I. D., Robinson, J. T., Sanborn, A. L., Machol, I., Omer, A. D., Lander, E. S., & Aiden, E. L. (2014, Dec 18). A 3D map of the human genome at kilobase resolution reveals principles of chromatin looping. Cell, 159(7), 1665–1680. https://doi.org/10.1016/j.cell.2014.11.021

Rezania, A., Bruin, J. E., Arora, P., Rubin, A., Batushansky, I., Asadi, A., O’Dwyer, S., Quiskamp, N., Mojibian, M., Albrecht, T., Yang, Y. H., Johnson, J. D., & Kieffer, T. J. (2014, Nov). Reversal of diabetes with insulin-producing cells derived in vitro from human pluripotent stem cells. Nat Biotechnol, 32(11), 1121–1133. https://doi.org/10.1038/nbt.3033

Roadmap Epigenomics, C., Kundaje, A., Meuleman, W., Ernst, J., Bilenky, M., Yen, A., Heravi-Moussavi, A., Kheradpour, P., Zhang, Z., Wang, J., Ziller, M. J., Amin, V., Whitaker, J. W., Schultz, M. D., Ward, L. D., Sarkar, A., Quon, G., Sandstrom, R. S., Eaton, M. L., Wu, Y. C., Pfenning, A. R., Wang, X., Claussnitzer, M., Liu, Y., Coarfa, C., Harris, R. A., Shoresh, N., Epstein, C. B., Gjoneska, E., Leung, D., Xie, W., Hawkins, R. D., Lister, R., Hong, C., Gascard, P., Mungall, A. J., Moore, R., Chuah, E., Tam, A., Canfield, T. K., Hansen, R. S., Kaul, R., Sabo, P. J., Bansal, M. S., Carles, A., Dixon, J. R., Farh, K. H., Feizi, S., Karlic, R., Kim, A. R., Kulkarni, A., Li, D., Lowdon, R., Elliott, G., Mercer, T. R., Neph, S. J., Onuchic, V., Polak, P., Rajagopal, N., Ray, P., Sallari, R. C., Siebenthall, K. T., Sinnott-Armstrong, N. A., Stevens, M., Thurman, R. E., Wu, J., Zhang, B., Zhou, X., Beaudet, A. E., Boyer, L. A., De Jager, P. L., Farnham, P. J., Fisher, S. J., Haussler, D., Jones, S. J., Li, W., Marra, M. A., McManus, M. T., Sunyaev, S., Thomson, J. A., Tlsty, T. D., Tsai, L. H., Wang, W., Waterland, R. A., Zhang, M. Q., Chadwick, L. H., Bernstein, B. E., Costello, J. F., Ecker, J. R., Hirst, M., Meissner, A., Milosavljevic, A., Ren, B., Stamatoyannopoulos, J. A., Wang, T., & Kellis, M. (2015, Feb 19). Integrative analysis of 111 reference human epigenomes. Nature, 518(7539), 317–330. https://doi.org/10.1038/nature14248

Rozowsky, J., Abyzov, A., Wang, J., Alves, P., Raha, D., Harmanci, A., Leng, J., Bjornson, R., Kong, Y., Kitabayashi, N., Bhardwaj, N., Rubin, M., Snyder, M., & Gerstein, M. (2011, Aug 2). AlleleSeq: analysis of allele-specific expression and binding in a network framework. Mol Syst Biol, 7, 522. https://doi.org/10.1038/msb.2011.54

Saxena, R., Hivert, M. F., Langenberg, C., Tanaka, T., Pankow, J. S., Vollenweider, P., Lyssenko, V., Bouatia-Naji, N., Dupuis, J., Jackson, A. U., Kao, W. H., Li, M., Glazer, N. L., Manning, A. K., Luan, J., Stringham, H. M., Prokopenko, I., Johnson, T., Grarup, N., Boesgaard, T. W., Lecoeur, C., Shrader, P., O’Connell, J., Ingelsson, E., Couper, D. J., Rice, K., Song, K., Andreasen, C. H., Dina, C., Kottgen, A., Le Bacquer, O., Pattou, F., Taneera, J., Steinthorsdottir, V., Rybin, D., Ardlie, K., Sampson, M., Qi, L., van Hoek, M., Weedon, M. N., Aulchenko, Y. S., Voight, B. F., Grallert, H., Balkau, B., Bergman, R. N., Bielinski, S. J., Bonnefond, A., Bonnycastle, L. L., Borch-Johnsen, K., Bottcher, Y., Brunner, E., Buchanan, T. A., Bumpstead, S. J., Cavalcanti-Proenca, C., Charpentier, G., Chen, Y. D., Chines, P. S., Collins, F. S., Cornelis, M., G, J. C., Delplanque, J., Doney, A., Egan, J. M., Erdos, M. R., Firmann, M., Forouhi, N. G., Fox, C. S., Goodarzi, M. O., Graessler, J., Hingorani, A., Isomaa, B., Jorgensen, T., Kivimaki, M., Kovacs, P., Krohn, K., Kumari, M., Lauritzen, T., Levy-Marchal, C., Mayor, V., McAteer, J. B., Meyre, D., Mitchell, B. D., Mohlke, K. L., Morken, M. A., Narisu, N., Palmer, C. N., Pakyz, R., Pascoe, L., Payne, F., Pearson, D., Rathmann, W., Sandbaek, A., Sayer, A. A., Scott, L. J., Sharp, S. J., Sijbrands, E., Singleton, A., Siscovick, D. S., Smith, N. L., Sparso, T., Swift, A. J., Syddall, H., Thorleifsson, G., Tonjes, A., Tuomi, T., Tuomilehto, J., Valle, T. T., Waeber, G., Walley, A., Waterworth, D. M., Zeggini, E., Zhao, J. H., consortium, G., investigators, M., Illig, T., Wichmann, H. E., Wilson, J. F., van Duijn, C., Hu, F. B., Morris, A. D., Frayling, T. M., Hattersley, A. T., Thorsteinsdottir, U., Stefansson, K., Nilsson, P., Syvanen, A. C., Shuldiner, A. R., Walker, M., Bornstein, S. R., Schwarz, P., Williams, G. H., Nathan, D. M., Kuusisto, J., Laakso, M., Cooper, C., Marmot, M., Ferrucci, L., Mooser, V., Stumvoll, M., Loos, R. J., Altshuler, D., Psaty, B. M., Rotter, J. I., Boerwinkle, E., Hansen, T., Pedersen, O., Florez, J. C., McCarthy, M. I., Boehnke, M., Barroso, I., Sladek, R., Froguel, P., Meigs, J. B., Groop, L., Wareham, N. J., & Watanabe, R. M. (2010, Feb). Genetic variation in GIPR influences the glucose and insulin responses to an oral glucose challenge. Nat Genet, 42(2), 142–148. https://doi.org/10.1038/ng.521

Schulz, T. C., Young, H. Y., Agulnick, A. D., Babin, M. J., Baetge, E. E., Bang, A. G., Bhoumik, A., Cepa, I., Cesario, R. M., Haakmeester, C., Kadoya, K., Kelly, J. R., Kerr, J., Martinson, L. A., McLean, A. B., Moorman, M. A., Payne, J. K., Richardson, M., Ross, K. G., Sherrer, E. S., Song, X., Wilson, A. Z., Brandon, E. P., Green, C. E., Kroon, E. J., Kelly, O. G., D’Amour, K. A., & Robins, A. J. (2012). A scalable system for production of functional pancreatic progenitors from human embryonic stem cells. PLoS One, 7(5), e37004. https://doi.org/10.1371/journal.pone.0037004

Sharon, N., Vanderhooft, J., Straubhaar, J., Mueller, J., Chawla, R., Zhou, Q., Engquist, E. N., Trapnell, C., Gifford, D. K., & Melton, D. A. (2019, May 21). Wnt Signaling Separates the Progenitor and Endocrine Compartments during Pancreas Development. Cell Rep, 27(8), 2281–2291 e2285. https://doi.org/10.1016/j.celrep.2019.04.083

Shi, Z. D., Lee, K., Yang, D., Amin, S., Verma, N., Li, Q. V., Zhu, Z., Soh, C. L., Kumar, R., Evans, T., Chen, S., & Huangfu, D. (2017, May 4). Genome Editing in hPSCs Reveals GATA6 Haploinsufficiency and a Genetic Interaction with GATA4 in Human Pancreatic Development. Cell Stem Cell, 20(5), 675–688 e676. https://doi.org/10.1016/j.stem.2017.01.001

Slavotinek, A., Kaylor, J., Pierce, H., Cahr, M., DeWard, S. J., Schneidman-Duhovny, D., Alsadah, A., Salem, F., Schmajuk, G., & Mehta, L. (2015, Jan 8). CRB2 mutations produce a phenotype resembling congenital nephrosis, Finnish type, with cerebral ventriculomegaly and raised alpha-fetoprotein. Am J Hum Genet, 96(1), 162–169. https://doi.org/10.1016/j.ajhg.2014.11.013

Steinthorsdottir, V., Thorleifsson, G., Sulem, P., Helgason, H., Grarup, N., Sigurdsson, A., Helgadottir, H. T., Johannsdottir, H., Magnusson, O. T., Gudjonsson, S. A., Justesen, J. M., Harder, M. N., Jorgensen, M. E., Christensen, C., Brandslund, I., Sandbaek, A., Lauritzen, T., Vestergaard, H., Linneberg, A., Jorgensen, T., Hansen, T., Daneshpour, M. S., Fallah, M. S., Hreidarsson, A. B., Sigurdsson, G., Azizi, F., Benediktsson, R., Masson, G., Helgason, A., Kong, A., Gudbjartsson, D. F., Pedersen, O., Thorsteinsdottir, U., & Stefansson, K. (2014, Mar). Identification of low-frequency and rare sequence variants associated with elevated or reduced risk of type 2 diabetes. Nat Genet, 46(3), 294–298. https://doi.org/10.1038/ng.2882

Strawbridge, R. J., Dupuis, J., Prokopenko, I., Barker, A., Ahlqvist, E., Rybin, D., Petrie, J. R., Travers, M. E., Bouatia-Naji, N., Dimas, A. S., Nica, A., Wheeler, E., Chen, H., Voight, B. F., Taneera, J., Kanoni, S., Peden, J. F., Turrini, F., Gustafsson, S., Zabena, C., Almgren, P., Barker, D. J., Barnes, D., Dennison, E. M., Eriksson, J. G., Eriksson, P., Eury, E., Folkersen, L., Fox, C. S., Frayling, T. M., Goel, A., Gu, H. F., Horikoshi, M., Isomaa, B., Jackson, A. U., Jameson, K. A., Kajantie, E., Kerr-Conte, J., Kuulasmaa, T., Kuusisto, J., Loos, R. J., Luan, J., Makrilakis, K., Manning, A. K., Martinez-Larrad, M. T., Narisu, N., Nastase Mannila, M., Ohrvik, J., Osmond, C., Pascoe, L., Payne, F., Sayer, A. A., Sennblad, B., Silveira, A., Stancakova, A., Stirrups, K., Swift, A. J., Syvanen, A. C., Tuomi, T., van ’t Hooft, F. M., Walker, M., Weedon, M. N., Xie, W., Zethelius, B., Consortium, D., Consortium, G., Mu, T. C., Consortium, C. A., Consortium, C. D., Ongen, H., Malarstig, A., Hopewell, J. C., Saleheen, D., Chambers, J., Parish, S., Danesh, J., Kooner, J., Ostenson, C. G., Lind, L., Cooper, C. C., Serrano-Rios, M., Ferrannini, E., Forsen, T. J., Clarke, R., Franzosi, M. G., Seedorf, U., Watkins, H., Froguel, P., Johnson, P., Deloukas, P., Collins, F. S., Laakso, M., Dermitzakis, E. T., Boehnke, M., McCarthy, M. I., Wareham, N. J., Groop, L., Pattou, F., Gloyn, A. L., Dedoussis, G. V., Lyssenko, V., Meigs, J. B., Barroso, I., Watanabe, R. M., Ingelsson, E., Langenberg, C., Hamsten, A., & Florez, J. C. (2011, Oct). Genome-wide association identifies nine common variants associated with fasting proinsulin levels and provides new insights into the pathophysiology of type 2 diabetes. Diabetes, 60(10), 2624–2634. https://doi.org/10.2337/db11-0415

Sui, L., Geens, M., Sermon, K., Bouwens, L., & Mfopou, J. K. (2013, Oct). Role of BMP signaling in pancreatic progenitor differentiation from human embryonic stem cells. Stem Cell Rev Rep, 9(5), 569–577. https://doi.org/10.1007/s12015-013-9435-6

Taal, H. R., Pourcain, B. S., Thiering, E., Das, S., Mook-Kanamori, D. O., Warrington, N. M., Kaakinen, M., Kreiner-Moller, E., Bradfield, J. P., Freathy, R. M., Geller, F., Guxens, M., Cousminer, D. L., Kerkhof, M., Timpson, N. J., Ikram, M. A., Beilin, L. J., Bonnelykke, K., Buxton, J. L., Charoen, P., Chawes, B. L. K., Eriksson, J., Evans, D. M., Hofman, A., Kemp, J. P., Kim, C. E., Klopp, N., Lahti, J., Lye, S. J., McMahon, G., Mentch, F. D., Muller, M., O’Reilly, P. F., Prokopenko, I., Rivadeneira, F., Steegers, E. A. P., Sunyer, J., Tiesler, C., Yaghootkar, H., Cohorts for, H., Aging Research in Genetic Epidemiology, C., Breteler, M. M. B., Debette, S., Fornage, M., Gudnason, V., Launer, L. J., van der Lugt, A., Mosley, T. H., Seshadri, S., Smith, A. V., Vernooij, M. W., Early, G., Lifecourse Epidemiology, c., Blakemore, A. I., Chiavacci, R. M., Feenstra, B., Fernandez-Benet, J., Grant, S. F. A., Hartikainen, A. L., van der Heijden, A. J., Iniguez, C., Lathrop, M., McArdle, W. L., Molgaard, A., Newnham, J. P., Palmer, L. J., Palotie, A., Pouta, A., Ring, S. M., Sovio, U., Standl, M., Uitterlinden, A. G., Wichmann, H. E., Vissing, N. H., DeCarli, C., van Duijn, C. M., McCarthy, M. I., Koppelman, G. H., Estivill, X., Hattersley, A. T., Melbye, M., Bisgaard, H., Pennell, C. E., Widen, E., Hakonarson, H., Smith, G. D., Heinrich, J., Jarvelin, M. R., Early Growth Genetics, C., & Jaddoe, V. W. V. (2012, Apr 15). Common variants at 12q15 and 12q24 are associated with infant head circumference. Nat Genet, 44(5), 532–538. https://doi.org/10.1038/ng.2238

Thurner, M., van de Bunt, M., Torres, J. M., Mahajan, A., Nylander, V., Bennett, A. J., Gaulton, K. J., Barrett, A., Burrows, C., Bell, C. G., Lowe, R., Beck, S., Rakyan, V. K., Gloyn, A. L., & McCarthy, M. I. (2018, Feb 7). Integration of human pancreatic islet genomic data refines regulatory mechanisms at Type 2 Diabetes susceptibility loci. Elife, 7. https://doi.org/10.7554/eLife.31977

Tiyaboonchai, A., Cardenas-Diaz, F. L., Ying, L., Maguire, J. A., Sim, X., Jobaliya, C., Gagne, A. L., Kishore, S., Stanescu, D. E., Hughes, N., De Leon, D. D., French, D. L., & Gadue, P. (2017, Mar 14). GATA6 Plays an Important Role in the Induction of Human Definitive Endoderm, Development of the Pancreas, and Functionality of Pancreatic beta Cells. Stem Cell Reports, 8(3), 589–604. https://doi.org/10.1016/j.stemcr.2016.12.026

Trapnell, C., Williams, B. A., Pertea, G., Mortazavi, A., Kwan, G., van Baren, M. J., Salzberg, S. L., Wold, B. J., & Pachter, L. (2010, May). Transcript assembly and quantification by RNA-Seq reveals unannotated transcripts and isoform switching during cell differentiation. Nat Biotechnol, 28(5), 511–515. https://doi.org/10.1038/nbt.1621

Urakami, T. (2019). Maturity-onset diabetes of the young (MODY): current perspectives on diagnosis and treatment. Diabetes Metab Syndr Obes, 12, 1047–1056. https://doi.org/10.2147/DMSO.S179793

van der Valk, R. J., Kreiner-Moller, E., Kooijman, M. N., Guxens, M., Stergiakouli, E., Saaf, A., Bradfield, J. P., Geller, F., Hayes, M. G., Cousminer, D. L., Korner, A., Thiering, E., Curtin, J. A., Myhre, R., Huikari, V., Joro, R., Kerkhof, M., Warrington, N. M., Pitkanen, N., Ntalla, I., Horikoshi, M., Veijola, R., Freathy, R. M., Teo, Y. Y., Barton, S. J., Evans, D. M., Kemp, J. P., St Pourcain, B., Ring, S. M., Davey Smith, G., Bergstrom, A., Kull, I., Hakonarson, H., Mentch, F. D., Bisgaard, H., Chawes, B., Stokholm, J., Waage, J., Eriksen, P., Sevelsted, A., Melbye, M., Early, G., Lifecourse Epidemiology, C., van Duijn, C. M., Medina-Gomez, C., Hofman, A., de Jongste, J. C., Taal, H. R., Uitterlinden, A. G., Genetic Investigation of, A. T. C., Armstrong, L. L., Eriksson, J., Palotie, A., Bustamante, M., Estivill, X., Gonzalez, J. R., Llop, S., Kiess, W., Mahajan, A., Flexeder, C., Tiesler, C. M., Murray, C. S., Simpson, A., Magnus, P., Sengpiel, V., Hartikainen, A. L., Keinanen-Kiukaanniemi, S., Lewin, A., Da Silva Couto Alves, A., Blakemore, A. I., Buxton, J. L., Kaakinen, M., Rodriguez, A., Sebert, S., Vaarasmaki, M., Lakka, T., Lindi, V., Gehring, U., Postma, D. S., Ang, W., Newnham, J. P., Lyytikainen, L. P., Pahkala, K., Raitakari, O. T., Panoutsopoulou, K., Zeggini, E., Boomsma, D. I., Groen-Blokhuis, M., Ilonen, J., Franke, L., Hirschhorn, J. N., Pers, T. H., Liang, L., Huang, J., Hocher, B., Knip, M., Saw, S. M., Holloway, J. W., Melen, E., Grant, S. F., Feenstra, B., Lowe, W. L., Widen, E., Sergeyev, E., Grallert, H., Custovic, A., Jacobsson, B., Jarvelin, M. R., Atalay, M., Koppelman, G. H., Pennell, C. E., Niinikoski, H., Dedoussis, G. V., McCarthy, M. I., Frayling, T. M., Sunyer, J., Timpson, N. J., Rivadeneira, F., Bonnelykke, K., Jaddoe, V. W., & Early Growth Genetics, C. (2015, Feb 15). A novel common variant in DCST2 is associated with length in early life and height in adulthood. Hum Mol Genet, 24(4), 1155–1168. https://doi.org/10.1093/hmg/ddu510

Varelas, X., Samavarchi-Tehrani, P., Narimatsu, M., Weiss, A., Cockburn, K., Larsen, B. G., Rossant, J., & Wrana, J. L. (2010, Dec 14). The Crumbs complex couples cell density sensing to Hippo-dependent control of the TGF-beta-SMAD pathway. Dev Cell, 19(6), 831–844. https://doi.org/10.1016/j.devcel.2010.11.012

Varshney, A., Scott, L. J., Welch, R. P., Erdos, M. R., Chines, P. S., Narisu, N., Albanus, R. D., Orchard, P., Wolford, B. N., Kursawe, R., Vadlamudi, S., Cannon, M. E., Didion, J. P., Hensley, J., Kirilusha, A., Program, N. C. S., Bonnycastle, L. L., Taylor, D. L., Watanabe, R., Mohlke, K. L., Boehnke, M., Collins, F. S., Parker, S. C., & Stitzel, M. L. (2017, Feb 28). Genetic regulatory signatures underlying islet gene expression and type 2 diabetes. Proc Natl Acad Sci U S A, 114(9), 2301–2306. https://doi.org/10.1073/pnas.1621192114

Villar, D., Berthelot, C., Aldridge, S., Rayner, T. F., Lukk, M., Pignatelli, M., Park, T. J., Deaville, R., Erichsen, J. T., Jasinska, A. J., Turner, J. M., Bertelsen, M. F., Murchison, E. P., Flicek, P., & Odom, D. T. (2015, Jan 29). Enhancer evolution across 20 mammalian species. Cell, 160(3), 554–566. https://doi.org/10.1016/j.cell.2015.01.006

Wang, A., Yue, F., Li, Y., Xie, R., Harper, T., Patel, N. A., Muth, K., Palmer, J., Qiu, Y., Wang, J., Lam, D. K., Raum, J. C., Stoffers, D. A., Ren, B., & Sander, M. (2015, Apr 2). Epigenetic priming of enhancers predicts developmental competence of hESC-derived endodermal lineage intermediates. Cell Stem Cell, 16(4), 386–399. https://doi.org/10.1016/j.stem.2015.02.013

Westmoreland, J. J., Kilic, G., Sartain, C., Sirma, S., Blain, J., Rehg, J., Harvey, N., & Sosa-Pineda, B. (2012, Apr). Pancreas-specific deletion of Prox1 affects development and disrupts homeostasis of the exocrine pancreas. Gastroenterology, 142(4), 999–1009 e1006. https://doi.org/10.1053/j.gastro.2011.12.007

Whyte, W. A., Orlando, D. A., Hnisz, D., Abraham, B. J., Lin, C. Y., Kagey, M. H., Rahl, P. B., Lee, T. I., & Young, R. A. (2013, Apr 11). Master transcription factors and mediator establish super-enhancers at key cell identity genes. Cell, 153(2), 307–319. https://doi.org/10.1016/j.cell.2013.03.035

Wood, A. R., Jonsson, A., Jackson, A. U., Wang, N., van Leewen, N., Palmer, N. D., Kobes, S., Deelen, J., Boquete-Vilarino, L., Paananen, J., Stancakova, A., Boomsma, D. I., de Geus, E. J. C., Eekhoff, E. M. W., Fritsche, A., Kramer, M., Nijpels, G., Simonis-Bik, A., van Haeften, T. W., Mahajan, A., Boehnke, M., Bergman, R. N., Tuomilehto, J., Collins, F. S., Mohlke, K. L., Banasik, K., Groves, C. J., McCarthy, M. I., Diabetes Research on Patient, S., Pearson, E. R., Natali, A., Mari, A., Buchanan, T. A., Taylor, K. D., Xiang, A. H., Gjesing, A. P., Grarup, N., Eiberg, H., Pedersen, O., Chen, Y. D., Laakso, M., Norris, J. M., Smith, U., Wagenknecht, L. E., Baier, L., Bowden, D. W., Hansen, T., Walker, M., Watanabe, R. M., t Hart, L. M., Hanson, R. L., & Frayling, T. M. (2017, Aug). A Genome-Wide Association Study of IVGTT-Based Measures of First-Phase Insulin Secretion Refines the Underlying Physiology of Type 2 Diabetes Variants. Diabetes, 66(8), 2296–2309. https://doi.org/10.2337/db16-1452

Xie, R., Everett, L. J., Lim, H. W., Patel, N. A., Schug, J., Kroon, E., Kelly, O. G., Wang, A., D’Amour, K. A., Robins, A. J., Won, K. J., Kaestner, K. H., & Sander, M. (2013, Feb 7). Dynamic chromatin remodeling mediated by polycomb proteins orchestrates pancreatic differentiation of human embryonic stem cells. Cell Stem Cell, 12(2), 224–237. https://doi.org/10.1016/j.stem.2012.11.023

Zhang, Y., Liu, T., Meyer, C. A., Eeckhoute, J., Johnson, D. S., Bernstein, B. E., Nusbaum, C., Myers, R. M., Brown, M., Li, W., & Liu, X. S. (2008). Model-based analysis of ChIP-Seq (MACS). Genome Biol, 9(9), R137. https://doi.org/10.1186/gb-2008-9-9-r137

Zhou, Y., Zhou, B., Pache, L., Chang, M., Khodabakhshi, A. H., Tanaseichuk, O., Benner, C., & Chanda, S. K. (2019, Apr 3). Metascape provides a biologist-oriented resource for the analysis of systems-level datasets. Nat Commun, 10(1), 1523. https://doi.org/10.1038/s41467-019-09234-6

