## Supplementary material for "Pancreatic progenitor epigenome maps prioritize type 2 diabetes risk genes with roles in development": Essential Reagents Table

**Key Resources Table**

| Reagent type (species) or resource | Designation | Source or reference | Identifiers | Additional information |
|---|---|---|---|---|
| Antibody | APC Mouse IgG1, κ Isotype Control | BD Pharmingen™ | Cat# 555751, RRID:AB_398613 | Flow cytometry (1:100) |
| Antibody | chicken anti-GFP | Aves Labs | Cat# GFP-1020, RRID:AB_10000240 | Immunohistochemistry (1:200) |
| Antibody | Cy3-conjugated donkey anti-mouse | Jackson ImmunoResearch Labs | Cat# 715-165-150, RRID:AB_2340813 | Immunofluorescence (1:1000) |
| Antibody | DyLight 488-conjugated donkey anti-goat | Jackson ImmunoResearch Labs | Cat# 705-545-003, RRID:AB_2340428 | Immunofluorescence (1:500) |
| Antibody | goat anti-CTCF | Santa Cruz Biotechnology | Cat# SC-15914X, RRID:AB_2086899 | ChIP-seq (4 ug) |
| Antibody | goat anti-FOXA1 | Abcam | Cat# ab5089, RRID:AB_304744 | ChIP-seq (4 ug) |
| Antibody | goat anti-FOXA2 | Santa Cruz Biotechnology | Cat# sc-6554, RRID:AB_2262810 | ChIP-seq (4 ug) |

| | | | | |
|---|---|---|---|---|
| Antibody | goat anti-GATA4 | Santa Cruz Biotechnology | Cat# sc-1237, RRID:AB_2108747 | ChIP-seq (4 ug) |
| Antibody | goat anti-PDX1 | Abcam | Cat# ab47383, RRID:AB_2162359 | Immunofluorescence (1:500) |
| Antibody | guinea pig anti-Insulin | Biomeda | Cat# v2024 | Immunohistochemistry (1:200) |
| Antibody | mouse anti-Crb2a | ZIRC | Cat# Zs-4 | Immunohistochemistry (1:100) |
| Antibody | mouse anti-GATA6 | Santa Cruz Biotechnology | Cat# sc-9055, RRID:AB_2108768 | ChIP-seq (4 ug) |
| Antibody | mouse anti-NKX6.1 | Developmental Studies Hybridoma Bank | Cat# F64A6B4, RRID:AB_532380 | Immunofluorescence (1:300) |
| Antibody | mouse anti-NKX6.1-Alexa Fluor® 647 | BD Biosciences | Cat# 563338, RRID:AB_2738144 | Flow cytometry (1:5) |
| Antibody | mouse anti-PDX1-PE | BD Biosciences | Cat# 562161, RRID:AB_10893589 | Flow cytometry (1:10) |

| | | | | |
|---|---|---|---|---|
| Antibody | PE Mouse IgG1, κ Isotype Control | BD Pharmingen™ | Cat# 555749, RRID:AB_396091 | Flow cytometry (1:100) |
| Antibody | rabbit anti-H3K27ac | Active Motif | Cat# 39133, RRID:AB_2561016 | ChIP-seq (4 ug) |
| Antibody | rabbit anti-H3K4me1 | Abcam | Cat# ab8895, RRID:AB_306847 | ChIP-seq (4 ug) |
| Antibody | rabbit anti-HNF6 | Santa Cruz Biotechnology | Cat# sc-13050, RRID:AB_2251852 | ChIP-seq (4 ug) |
| Antibody | rabbit anti-laminin | Sigma | Cat# L9393, RRID:AB_477163 | Immunohistochemistry (1:100) |
| Antibody | rabbit anti-panCrb | Jensen Laboratory, University of Massachusetts, Amherst | N/A | Immunohistochemistry (1:100) |
| Antibody | rabbit anti-PDX1 | Beta Cell Biology Consortium | AB1068 | ChIP-seq (4 ug) |
| Antibody | rabbit anti-SOX9 | Chemicon | Cat# 5535, RRID:AB_2239761 | ChIP-seq (4 ug) |

| | | | | |
|---|---|---|---|---|
| Cell line (*Homo-sapiens*, male) | CyT49 | ViaCyte, Inc. | NIHhESC-10-0041, RRID:CVCL_B850 | |
| Cell line (*Homo-sapiens*, male) | H1 | WiCell Research Institute | NIHhESC-10-0043, RRID:CVCL_9771 | |
| Chemical compound, drug | 2-Mercaptoethanol | Thermo Fisher Scientific | Cat# 21985023 | |
| Chemical compound, drug | Accutase | Thermo Fisher Scientific | Cat# 00-4555-56 | |
| Chemical compound, drug | B-27 supplement | Thermo Fisher Scientific | Cat# 17504044 | |
| Chemical compound, drug | Bovine Albumin Fraction V | Life Technologies | Cat# 15260037 | |
| Chemical compound, drug | D-(+)-Glucose Solution, 45% | Sigma-Aldrich | Cat# G8769 | |
| Chemical compound, drug | DAPI | Invitrogen | Cat# D1306 | Immunohistochemistry (1:200) |

| | | | |
|---|---|---|---|
| Chemical compound, drug | DMEM High Glucose | VWR | Cat# 16750-082 |
| Chemical compound, drug | DMEM/F12 [-] L-glutamine | VWR | Cat# 15-090-CV |
| Chemical compound, drug | DMEM/F12 with L-Glutamine, HEPES | Corning | Cat# 45000-350 |
| Chemical compound, drug | DMF | EMD Millipore | Cat# DX1730 |
| Chemical compound, drug | DPBS | Thermo Fisher Scientific | Cat# 21-031-CV |
| Chemical compound, drug | DTT | Sigma | Cat# D9779 |
| Chemical compound, drug | Fetal Bovine Serum | Thermo Fisher Scientific | Cat# MT35011CV |
| Chemical compound, drug | Glutamax | Thermo Fisher Scientific | Cat# 35050-079 |

| | | | |
|---|---|---|---|
| Chemical compound, drug | GlutaMAX | Thermo Fisher Scientific | Cat# 35050061 |
| Chemical compound, drug | Hoechst 33342 | Thermo Fisher Scientific | Cat# H3570 |
| Chemical compound, drug | HyClone Dulbecco's Modified Eagles Medium | Thermo Fisher Scientific | Cat# SH30081.FS |
| Chemical compound, drug | IGEPAL-CA630 | Sigma | Cat# I8896 |
| Chemical compound, drug | Illumina tagmentation enzyme | Illumina | Cat# FC-121-1030 |
| Chemical compound, drug | Insulin-Transferrin-Selenium (ITS) | Thermo Fisher Scientific | Cat# 41400045 |
| Chemical compound, drug | Insulin-Transferrin-Selenium-Ethanolamine (ITS-X) | Thermo Fisher Scientific | Cat# 51500-056 |
| Chemical compound, drug | KAAD-Cyclopamine | Toronto Research Chemicals | Cat# K171000 |

| | | | |
|---|---|---|---|
| Chemical compound, drug | K-acetate | Sigma | Cat# P5708 |
| Chemical compound, drug | KnockOut SR XenoFree | Thermo Fisher Scientific | Cat# A1099202 |
| Chemical compound, drug | LDN-193189 | Stemgent | Cat# 04-0074 |
| Chemical compound, drug | Matrigel® | Corning | Cat# 356231 |
| Chemical compound, drug | MCDB 131 | Thermo Fisher Scientific | Cat# 10372-019 |
| Chemical compound, drug | Mg-acetate | Sigma | Cat# M2545 |
| Chemical compound, drug | mTeSR1 Complete Kit - GMP | STEMCELL Technologies | Cat# 85850 |
| Chemical compound, drug | NEBNext High-Fidelity 2X PCR Master Mix | NEB | Cat# M0541 |

| | | | |
|---|---|---|---|
| Chemical compound, drug | Non-Essential Amino Acids | Thermo Fisher Scientific | Cat# 11140050 |
| Chemical compound, drug | O.C.T. Compound | Sakura Finetek USA | Cat# 25608-930 |
| Chemical compound, drug | Penicillin-Streptomycin | Thermo Fisher Scientific | Cat# 15140122 |
| Chemical compound, drug | Polyethylenimine (PEI) | Polysciences | Cat# 23966-1 |
| Chemical compound, drug | Protease inhibitor | Roche | Cat# 05056489001 |
| Chemical compound, drug | Retinoic Acid | Sigma-Aldrich | Cat# R2625 |
| Chemical compound, drug | RNA ScreenTape Sample Buffer | Agilent Technologies | Cat# 5067-5577 |
| Chemical compound, drug | ROCK Inhibitor Y-27632 | STEMCELL Technologies | Cat# 72305 |

| | | | |
|---|---|---|---|
| Chemical compound, drug | RPMI 1640 [-] L-glutamine | VWR | Cat# 15-040-CV |
| Chemical compound, drug | SANT-1 | Sigma-Aldrich | Cat# S4572 |
| Chemical compound, drug | Sodium Bicarbonate | Sigma-Aldrich | Cat# NC0564699 |
| Chemical compound, drug | Tamoxifen | Sigma | Cat# T5648 |
| Chemical compound, drug | TGF-β RI Kinase Inhibitor IV | Calbiochem | Cat# 616454 |
| Chemical compound, drug | TPB | Calbiochem | Cat# 565740 |
| Chemical compound, drug | Tranylcypromine | Cayman Chemical | Cat# 10010494 |
| Chemical compound, drug | Tris-acetate | Thermo Fisher Scientific | Cat# BP-152 |

| | | | |
|---|---|---|---|
| Chemical compound, drug | TTNPB | Enzo Life Sciences | Cat# BML-GR105 |
| Chemical compound, drug | Vectashield Antifade Mounting Medium | Vector Laboratories | Cat# H-1000 |
| Chemical compound, drug | XtremeGene 9 | Roche | Cat# 6365787001 |
| Commercial assay | High Sensitivity D1000 ScreenTape | Agilent Technologies | Cat# 5067-5584 |
| Commercial assay | RNA ScreenTape | Agilent Technologies | Cat# 5067-5576 |
| Commercial assay | RNA ScreenTape Ladder | Agilent Technologies | Cat# 5067-5578 |
| Commercial assay | BD Cytofix/Cytoperm™ Plus Fixation/Permeabilization Solution Kit | BD Biosciences | Cat# 554715 |
| Commercial assay | ChIP-IT High Sensitivity Kit | Active Motif | Cat# 53040 |

| | | | |
|---|---|---|---|
| Commercial assay | iQ SYBR Green Supermix | Bio-Rad | Cat# 1708884 |
| Commercial assay | iScript cDNA Synthesis Kit | Bio-Rad | Cat# 1708891 |
| Commercial assay | KAPA Library Preparation Kit (Illumina) | Kapa Biosystems | Cat# KK8234 |
| Commercial assay | KAPA Stranded mRNA-Seq Kits | Kapa Biosystems | Cat# KK8401 |
| Commercial assay | MinElute PCR purification kit | QIAGEN | Cat# 28004 |
| Commercial assay | Qubit ssDNA assay kit | Thermo Fisher Scientific | Cat# Q10212 |
| Commercial assay | RNeasy Micro Kit | QIAGEN | Cat# 74004 |
| genetic reagent (*D. rerio*) | *TgBAC(pdx1:eGFP)*$^{bns13}$ | PMID: 16258076 | N/A |

| | | | |
|---|---|---|---|
| genetic reagent (*D. rerio*) | *Tg(ptf1a:eGFP)$^{jh1}$* | PMID: 16258076 | N/A |
| Other | SPRIselect bead | Beckman Coulter | Cat# B23317 |
| Recombinant protein | Activin A | R&D Systems | Cat# 338-AC/CF |
| Recombinant protein | Human AB Serum | Valley Biomedical | Cat# HP1022 |
| Recombinant protein | Recombinant EGF | R&D Systems | Cat# 236-EG |
| Recombinant protein | Recombinant Heregulinβ-1 | Peprotech | Cat# 100-03 |
| Recombinant protein | Recombinant KGF/FGF7 | R&D Systems | Cat# 251-KG |
| Recombinant protein | Recombinant Mouse Wnt3A | R&D Systems | Cat# 1324-WN/CF |

| | | | | |
|---|---|---|---|---|
| Recombinant protein | Recombinant Noggin | R&D Systems | Cat# 3344NG | |
| Sequence-based reagent | Px333 Plasmid | http://www.addgene.org/64073/ | RRID:Addgene_64073 | |
| Sequence-based reagent | LAMA1 Forward | This paper | qPCR primers | GTG ATG GCA ACA GCG CAA A |
| Sequence-based reagent | LAMA1 Reverse | This paper | qPCR primers | GAC CCA GTG ATA TTC TCT CCC A |
| Sequence-based reagent | CRB2 Forward | This paper | qPCR primers | ACC ACT GTG CTT GTC CTG AG |
| Sequence-based reagent | CRB2 Reverse | This paper | qPCR primers | TCC AGG GTC GCT AGA TGG AG |
| Sequence-based reagent | TBP Forward | This paper | qPCR primers | TGT GCA CAG GAG CCA AGA GT |
| Sequence-based reagent | TBP Reverse | This paper | qPCR primers | ATT TTC TTG CTG CCA GTC TGG |

| | | | | |
|---|---|---|---|---|
| Sequence-based reagent | *LAMA1*Enh Upstream Guide | This paper | CRISPR sgRNA | GTC AAA TTG CTA TAA CAC GG |
| Sequence-based reagent | *LAMA1*Enh Downstream Guide | This paper | CRISPR sgRNA | CCA CTT TAA GTA TCT CAG CA |
| Sequence-based reagent | *CRB2*Enh Upstream Guide | This paper | CRISPR sgRNA | ATA CAA AGC ACG TGA GA |
| Sequence-based reagent | *CRB2*Enh Downstream Guide | This paper | CRISPR sgRNA | GAA TGC GGA TGA CGC CTG AG |
| Sequence-based reagent | lama1-ATG | PMID: 16321372 | Morpholino | TCA TCC TCA TCT CCA TCA TCG CTC A Obtained from GeneTools, LLC |
| Sequence-based reagent | crb2a-SP | PMID: 16713951 | Morpholino | ACG TTG CCA GTA CCT GTG TAT CCT G Obtained from GeneTools, LLC |
| Sequence-based reagent | crb2b-SP | PMID: 16713951 | Morpholino | TAA AGA TGT CCT ACC CAG CTT GAA C Obtained from GeneTools, LLC |
| Sequence-based reagent | standard control MO | N/A | Morpholino | CCT CTT ACC TCA GTT ACA ATT TAT A Obtained from GeneTools, LLC |

| | | | |
|---|---|---|---|
| Software, algorithm | Adobe Illustrator v 5.1 | http://www.adobe.com/products/illustrator.html | RRID:SCR_014198 |
| Software, algorithm | Adobe Photoshop v 5.1 | http://www.adobe.com/products/photoshop.html | RRID:SCR_014199 |
| Software, algorithm | BEDtools v 2.26.0 | https://github.com/arq5x/bedtools2 | RRID:SCR_006646 |
| Software, algorithm | Bioconductor | https://www.bioconductor.org/ | RRID: SCR_006442 |
| Software, algorithm | Burrows-Wheeler Aligner v 0.7.13 | http://bio-bwa.sourceforge.net/ | RRID:SCR_010910 |
| Software, algorithm | CENTIPEDE v 1.2 | http://centipede.uchicago.edu/ | N/A |
| Software, algorithm | Cufflinks v 2.2.1 | http://cole-trapnell-lab.github.io/cufflinks/ | RRID:SCR_014597 |
| Software, algorithm | deepTools2 v 3.1.3 | https://deeptools.readthedocs.io/en/develop/content/installation.html | N/A |

| | | | |
|---|---|---|---|
| Software, algorithm | DESeq2 v 3.10 | https://bioconductor.org/packages/release/bioc/html/DESeq2.html | RRID:SCR_015687 |
| Software, algorithm | FlowJo v10 software | https://www.flowjo.com/solutions/flowjo | RRID: SCR_008520 |
| Software, algorithm | GraphPad Prism v 8.1.2 | https://www.graphpad.com/scientific-software/prism/ | RRID: SCR_002798 |
| Software, algorithm | HOMER v 4.10.4 | http://homer.ucsd.edu/homer/ | RRID: SCR_010881 |
| Software, algorithm | Juicebox Tools v 1.4 | https://github.com/aidenlab/Juicebox/wiki/Juicebox-Assembly-Tools | N/A |
| Software, algorithm | MACS2 v 2.1.4 | http://liulab.dfci.harvard.edu/MACS/ | RRID:SCR_013291 |
| Software, algorithm | MEME suite v 5.1.1 | http://meme-suite.org/ | RRID:SCR_001783 |
| Software, algorithm | Metascape | http://metscape.ncibi.org | RRID:SCR_014687 |

| | | | |
|---|---|---|---|
| Software, algorithm | Picard Tools v 1.131 | http://broadinstitute.github.io/picard/ | RRID:SCR_006525 |
| Software, algorithm | R Project for Statistical Computing v 3.6.1 | http://www.r-project.org/ | RRID:SCR_001905 |
| Software, algorithm | SAMtools v 1.5 | http://samtools.sourceforge.net | RRID:SCR_002105 |
| Software, algorithm | STAR v 2.4 | https://github.com/alexdobin/STAR | N/A |
| Software, algorithm | UCSC Genome Browser | http://genome.ucsc.edu/ | RRID:SCR_005780 |
| Software, algorithm | vcf2diploid v 0.2.6a | https://github.com/abyzovlab/vcf2diploid | N/A |
| Software, algorithm | ZEISS ZEN Digital Imaging for Light Microscopy | http://www.zeiss.com/microscopy/en_us/products/microscope-software/zen.html#introduction | RRID:SCR_013672 |
